# Universal consensus 3D segmentation of cells from 2D segmented stacks

**DOI:** 10.1101/2024.05.03.592249

**Authors:** Felix Y. Zhou, Zach Marin, Clarence Yapp, Qiongjing Zou, Benjamin A. Nanes, Stephan Daetwyler, Andrew R. Jamieson, Md Torikul Islam, Edward Jenkins, Gabriel M. Gihana, Jinlong Lin, Hazel M. Borges, Bo-Jui Chang, Andrew Weems, Sean J. Morrison, Peter K. Sorger, Reto Fiolka, Kevin M. Dean, Gaudenz Danuser

## Abstract

Cell segmentation is the foundation of a wide range of microscopy-based biological studies. Deep learning has revolutionized 2D cell segmentation, enabling generalized solutions across cell types and imaging modalities. This has been driven by the ease of scaling up image acquisition, annotation, and computation. However, 3D cell segmentation, requiring dense annotation of 2D slices still poses significant challenges. Manual labeling of 3D cells to train broadly applicable segmentation models is prohibitive. Even in high-contrast images annotation is ambiguous and time-consuming. Here we develop a theory and toolbox, u- Segment3D, for 2D-to-3D segmentation, compatible with any 2D method generating pixel-based instance cell masks. u-Segment3D translates and enhances 2D instance segmentations to a 3D consensus instance segmentation without training data, as demonstrated on 11 real-life datasets, >70,000 cells, spanning single cells, cell aggregates, and tissue. Moreover, u-Segment3D is competitive with native 3D segmentation, even exceeding when cells are crowded and have complex morphologies.

## Main

Instance segmentation is the problem of unambiguously assigning each pixel in a 2D or voxel in a 3D image to unique objects of interest. Near universally, it is the first step in quantitative image analysis for scientific fields including cell biology^1^. Only through segmentation are the objects of interest, such as nuclei^2,3^, organelles^4^, cells^5^, bacteria^6^, plants^7^, organs^8,9^ or vasculature^10^, explicitly identified and delineated within an image. The segmentation subsequently defines the quantitative unit of analysis to extract desired object features such as morphology^11^ (e.g. length, area, and volume) and molecular expression (e.g. mean marker expression^12^, subcellular patterns^13^).

The segmentation of individual cells is easy when they are isolated, well-contrasted and uniformly illuminated, and amenable to binary intensity thresholding and connected component analysis^14^. However, this scenario is rare. In practice, cells of diverse morphologies in culture in-vitro, or within tissues in-situ and in-vivo, may aggregate in clusters that cannot be easily or accurately separated by direct thresholding^1,15^. This is further compounded by variations in the staining and imaging process used to visualize cellular structures, often resulting in weak, partial, sparse, or unspecific foreground intensity signals^4,16^.

Thanks to advancements in GPU architecture and increased availability of publicly available labeled datasets, generalist or ‘foundational’ 2D cell segmentation models have emerged both for interactive segmentation using prompts such as µ-SAM^17^, CellSAM^18^ and for dense segmentation of every cell such as Cellpose^5^ and various transformer models^1^. These methods leverage ‘big data’ and harness diversity in the training data to achieve impressive ability in segmenting 2D cells acquired across modalities and cell types^1^ out-of-the-box or with fine-tuning.

Physiologically, however, cells interact within complex 3D environments. The importance of studying cell biological processes in the relevant physiological 3D environments is well-documented^11,19-22^. Moreover, the emergence of 3D *in-situ* tissue imaging has provided unprecedented insights into the complex nature of the tissue microenvironment and its role in development and disease; including novel cell-cell interaction, tissue organization, and diverse cell morphologies^12,23^. Unlocking the potential of 3D imaging necessarily requires reliable, general, and scalable 3D cell segmentation solutions. Simply replicating the training strategy of 2D foundation models is impossible as this requires significant amounts of well-labeled, diverse 3D cell datasets.

Despite the relative ease of image acquisition, abundance of industrial annotation tools in 2D^24,25^ and ease of crowd-sourcing and proofreading entire 2D image datasets^26^, the Cellpose training dataset comprises merely 540 training images (total ∼70,000 cells, 5 modalities) and the most recent and largest multimodal segmentation challenge^1^ only 1000 training images (total 168,491 cells, 4 modalities). Replicating a densely labeled 3D dataset with comparable levels of cell diversity and numbers, given more complex microenvironments, more variable image quality, and more diverse morphologies and cell packing would be a formidable undertaking^4,12,23,27^. Despite ongoing efforts to develop scalable 3D annotation tools^28-31^ with AI assistance^32,33^ and proofreading^34^, the generation of meaningful training data requires an extraordinary level of manual intervention^35-39^. Even with expert annotators, labelling suffers from inter- and intra-operator variation^40-42^ and is inherently biased towards easy cases. Consequently, both classical^43-46^ and deep- learning based^47-49^ 3D segmentation method development focus primarily on nuclei, which have well-defined round shapes, are separated from neighbors, and can be visualized with high contrast by nuclear dyes^50^. Some densely annotated, proofread datasets of 3D cells^51^ have been generated for plant tissues^7,52^, few cell aggregates^15,53^, and embryos^54^. However, these datasets comprise crops or timepoints of few unique biological samples and therefore are not reflective of the variability for true biological replicates. Synthetic^55,56^, partial^57^ or generative model^58,59^ derived datasets have been proposed to alleviate the need for fully labeled data but have only been demonstrated with star-convex morphologies. It is unclear how these methods generalize to more complex morphologies, microenvironments and future, novel 3D imaging modalities, as they still fundamentally use data-driven supervised learning.

High-quality, annotated datasets with solid ground truth and minimal noise^60^ are not the only limitations for 3D segmentation. The time to train or fine-tune foundation models is already a major consideration in 2D, requiring significant time investment, memory, specialized GPUs^1,9,17,18,61^ and careful dataset curation to ensure diversity^62^. Training comparable 3D models will not only require more time and dedicated resources, but faces additional challenges such as model overparameterization, necessitating more efficient, revised architecture designs^10,15,63^. Lastly, even if trained on a vast dataset, foundation models still cannot guarantee generalization and robustness^64,65^. Consequently, at the expense of reduced recall or accuracy, it is typically more efficient for academic labs with small datasets to adopt human-in-the-loop, interactive segmentation tools like ilastik^66^ or to use segmented nuclei as seeds for classical 3D watershed segmentation^27^.

To address the shortcomings of directly training 3D segmentation models, we revisit the idea of leveraging 2D cell segmentations from orthoviews without data retraining to generate a “consensus” 3D segmentation – a 3D translation of 2D segmentations in orthogonal slices, i.e. orthoslices, which leverages complementary information between orthoslices, and preserves or even improves the accuracy of the 2D inputs. Using 2D predictions to assist 3D inference is a common strategy to minimize computation and training. Primarily, this involves adapting pretrained 2D models to 3D, for example by inflating 2D convolutional kernels followed by fine-tuning in 3D^63^. Alternatively, 3D data are evaluated slice-by-slice and the outputs are processed and integrated by a separately trained 3D model^67^. Few works pursue a no-training approach. Almost universally, existing tools construct a 3D segmentation by matching and stitching 2D segmentations across x-y slices^5,26^ whereby the stitching is controlled by an overlap score. Relying on a single view, these 3D segmentations are notoriously rasterized and erroneously join multiple touching cells as tubes^26,33,50,68-70^. CellStitch^71^ and 3DCellComposer^50^ propose matching across orthogonal x-y, x-z, y-z views to create a more accurate consensus 3D segmentation. However, these discrete matching approaches are inherently difficult to scale-up with cell number, not applicable to branched structures and cannot easily handle missing, undersegmented or oversegmented cells across slices. Alternatively, Cellpose^5^ proposed to average predicted 2D flow vectors along the x-y, x-z and y-z directions to construct a 3D gradient map. By tracing the gradient map to the simulated heat origin, the 3D cell instances are found by grouping all voxels that flow to the same sink. Whilst conceptually elegant, its execution has been restricted to Cellpose predicted gradients and demonstrates limited performance on anisotropic^71^, noisy or morphologically non-ellipsoidal datasets^15^. Moreover, we and others^71^ have observed fragmentation of whole 3D cells into angular sectors, a behavior inconsistent with Cellpose’s representation of whole cells as a 360^0^ angular field, and worse than just stitching the 2D cell masks^71^.

To derive a formal framework for 2D-to-3D segmentation unifying stitching and gradient tracing, we revisited the problem from first principles. We find that 2D-to-3D translation can be formulated generally as an optimization problem, whereby we reconstruct the 3D gradient vectors of the distance transform representation of each cell’s 3D medial-axis skeleton. Notably, 3D cells can then be reconstructed using gradient descent and spatial connected component analysis. Together these two principles allow the implementation of a universal consensus 3D segmentation from 2D segmentations in one, two or all three orthoview stacks (e.g., in x-y, x-z and y-z). Here universality refers to the independence of the framework to cell morphology and which 2D segmentation method is employed to generate pixel-based instance segmentation masks. Named u-Segment3D, our implementation of this framework exposes parameters to account for imperfect input 2D segmentations. Moreover, it includes preprocessing and postprocessing steps to assist the application of pretrained models on unseen datasets and to recover missing 3D segmentation features. We first describe our formalism of 2D-to-3D segmentation. We then validate u-Segment3D by optimal, near perfect reconstruction of reference 3D shapes from 11 real-life datasets, encompassing >70,000 cells from cell aggregates, embryos, tissue, and entire vasculature networks. We use pretrained Cellpose 2D models to demonstrate the efficacy of u-Segment3D in faithfully translating the 2D segmentation performance into consensus 3D segmentations. We also compare the performance of the 2D- to-3D segmentation approach on datasets for which native 3D segmentation models are available. We find that consensus 3D segmentations match, and for crowded cells or complex, branched morphologies even exceed native 3D segmentations. In sum, u-Segment3D’s implementation of 2D-to-3D segmentation guarantees that the better the 2D segmentation, the better the resultant 3D segmentation. Finally, we demonstrate the flexibility and capacity of u-Segment3D to segment unseen 3D volume data sets of anisotropic cell cultures, and unwrapped embryo surfaces^72^; high-resolution single cells and cell aggregates with intricate surface protrusions^73^; thin, sprouting vasculature in zebrafish, and tissue architectures imaged with spatial multiplexing^23^ and electron microscopy^74^.

u-Segment3D is implemented in Python 3 using open-source packages with multiprocessing capability. Example scripts are included to help users get started and leverage widely accessible CPU-based high-performance computing (HPC) clusters for segmenting larger 3D volumes. u-Segment3D is freely available and can be installed from https://github.com/DanuserLab/u-segment3D and the Python Package Index, PyPI.

## Results

### A formalism for 2D-to-3D segmentation

Dense instance segmentation identifies every object in the image and assigns a unique ID to all pixels and voxels comprising an object instance. This is equivalent to two separate tasks: (i) binary labeling every image voxel as foreground (value 1) or background (value 0), and (ii) assigning to each designated foreground voxel, a unique positive integer ID denoting the instance, (Fig. 1a). The critical challenge of 2D-to-3D segmentation is preserving the 2D cell boundaries in all orthoviews, and obtain the correct number of 3D instances after translation, whilst keeping computation low. We formulate an optimal reconstruction using 2D without explicit matching of segmentations, through spatial clustering in an eroded space, motivated from first principles.

**Figure 1.**
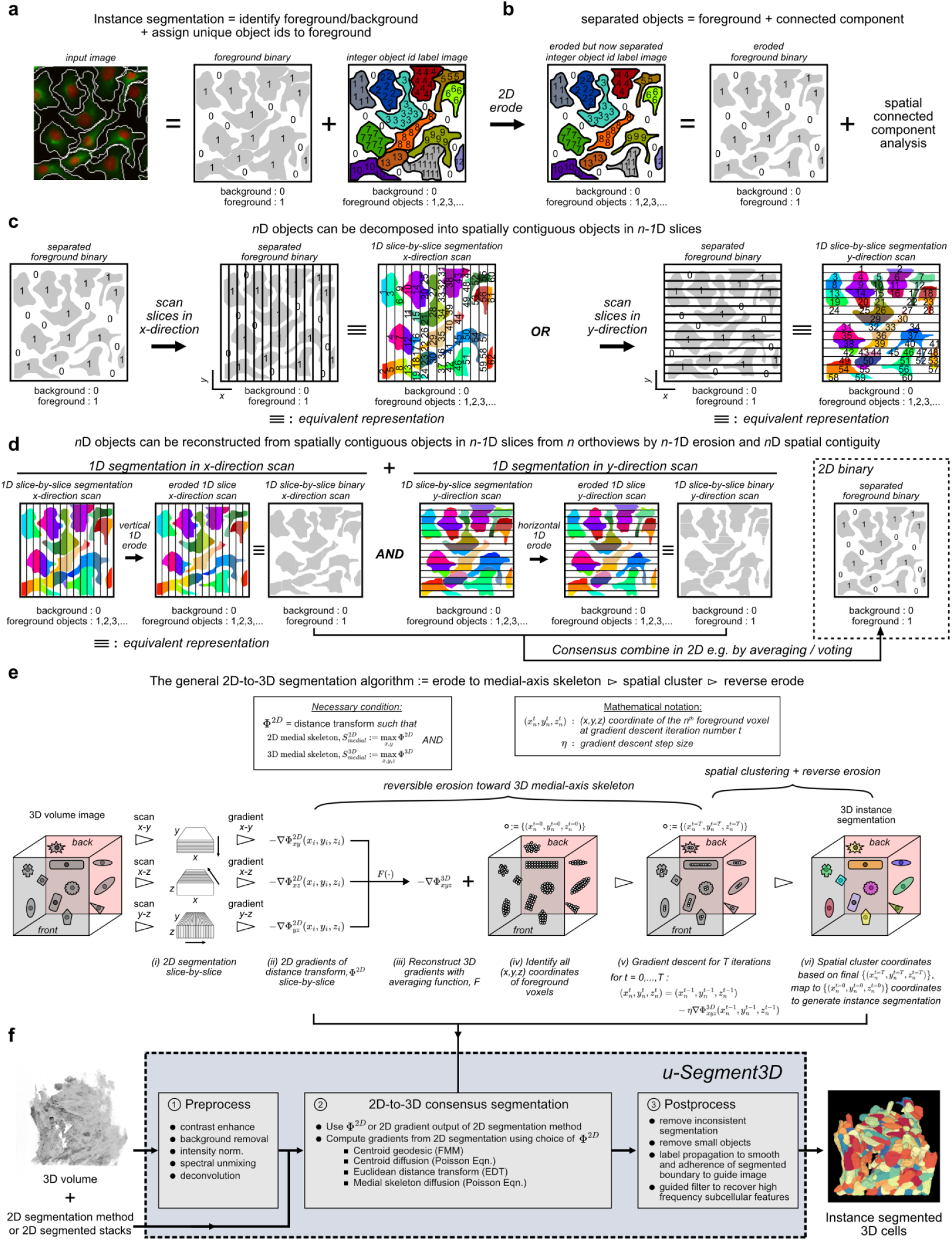
u-Segment3D is a toolbox for generating consensus instance 3D segmentation from 2D segmentations. **a)** Computational representation of the 2D segmentation of densely packed cells as two images, a foreground binary mask and a labelled image where each cell is assigned a unique integer id. **b)** Equivalent representation of the eroded segmentation where individual cells are spatially separated and can be equivalently represented using a foreground binary mask that can be parsed using connected component analysis to recover individual cell ids. **c)** Schematic of decomposing 2D instance cell segmentations into 1D slice segmentations scanning the 2D image in orthogonal *x*- or *y*- directions. **d**) Schematic of the reconstruction of 2D instance cell segmentations using 1D segmentations from orthogonal *x*- and *y*- direction scans of the 2D image to generate a consensus foreground followed by 2D spatial proximity clustering. Segmentations are eroded vertically or horizontally for *x*- or *y*- direction scans respectively. ≡ denotes equivalent representation. **e)** Schematic of the minimal set of algorithmic steps to operationalize the framework in d) for a consensus 3D segmentation of densely packed cells. **f)** u-Segment3D is a toolbox to enable the application of the algorithmic steps in e) to real datasets with additional preprocessing methods to adapt any pretrained 2D segmentation model or 2D method and postprocessing methods to improve and recover missing local features in the reconstructed 3D segmentation such as subcellular protrusions.

Starting with a 2D instance segmentation, if we erode each object border by 1 pixel, then all cell areas become spatially separated. In this scenario, cell instances are equally-well represented by a binary foreground/background image whereby object IDs are parsed by performing connected component analysis (Fig. 1b). However, unlike the discrete label representation, this binary image is now a continuous scalar field which can be decomposed and equally-well computed from its 1D slices in either x- or y-directions (Fig 1c). This is because the unique 2D objects are spatially contiguous across adjacent 1D regions. This implies that a 1D instance segmenter that accurately delineates individual cell boundaries with touching 1D cells can also be applied in reverse to reconstruct the 2D instance segmentation via its equivalent eroded 2D binary foreground, (Fig 1d). The 1D segmenter is run slice-by-slice in the x-direction. Each unique 1D ‘cell’ is eroded vertically within each x-slice then restacked to a 2D binary image. The erosion ensures spatial separation in the x-direction in the 2D binary, however, cells may still touch in the orthogonal y-direction. The binary generated by an x-direction scan must be combined by a consensus operation, e.g. computing a pixel-wise average (and re-binarize), or pixel-wise intersection, with the equivalent binary produced from an orthogonal y-direction scan to reconstruct a single foreground binary representing fully separated 2D cells. Spatial connected components can then identify and label all contiguous 2D regions as unique cells. The erosion in x- and y-slices is reversed for each 2D cell to recover touching cell boundaries. The same first-principle arguments apply to reconstructing an *n*D segmentation from minimally *n* orthogonal *n*-1D segmentations, i.e. for 3D cell segmentations from x-y, x-z, and y-z 2D slices.

In practice, this conceptual framework is suboptimal if directly applied to data. Images rasterize the cell shape, with resolution dictated by the grid size; the lower-dimensional segmenter is imperfect with inevitable missing or inconsistent segmentations across slices whereby a cell splits into multiple or multiple becoming one; and uniform morphological dilation after *n*D reconstruction will not preserve eroded curvature features. Discrete computational processes such as label matching and morphological operations, fundamentally cannot overcome these issues across orthoviews. The former cannot manipulate and correct erroneous inputs, either segmentations are matched or are discarded. The latter does not retain shape manipulation history. A central question is how to erode 2D cells to guarantee framework applicability to heterogeneous cell sizes and morphologies.

u-Segment3D proposes a continuous implementation to address all these issues. First, each cell is represented as a foreground binary and associated dense gradient field. This enables segmentations to be flexibly manipulated with continuous computations: smoothing to impute across slices or to join arbitrary number of neighboring cells; and averaging to obtain consensus, downvoting information unsupported by majority orthoviews. Second, we iteratively erode cell shapes to their medial-axis skeleton^75^ (MAT). By construction, the MAT maximally separates foreground points belonging to separate neighboring cells during erosion and is computable for any cell shape. Crucial for the eventual 2D-to-3D segmentation, the MAT of 2D slices coincides with the MAT of the corresponding 3D object. Resolution permitting, the 2D skeletal slices of each unique 3D object remain spatially proximal after 3D stacking, enabling identification by spatial proximity. Moreover, medial-axis skeletons are attractors of distance transforms^75,76^, Φ. The gradient field of a cell is therefore the distance transform gradient, and gradient descent specifies a reversible, continuous erosion process without shape information loss. A cell shape is eroded by advecting its foreground coordinates with step size *η*, in the direction of the local gradient, ∇Φ (Methods). The history of coordinates is the reverse erosion. The general 2D-to-3D segmentation algorithm (Fig. 1e) is thus:

1. Generate 2D pixel-based instance segmentations independently in orthogonal x-y, x-z, y-z views.
2. Convert cells in x-y, x-z, y-z views to their 2D gradient field representation by computing a 2D distance transform, Φ^2D^ whose attractor is a 2D medial-axis skeleton, and its 2D gradient, 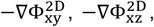, 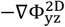 respectively. Should the 2D segmenter already predict suitable 2D gradients, e.g. Cellpose^5^, these can be directly used instead, (Methods).
3. Reconstruct the 3D gradients of the distance transform 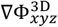 from the 2D gradients, using a consensus averaging function, *F* (Methods).

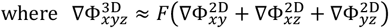

Apply *F* to the 3D foreground stacked from x-y, x-z, and y-z views and binarize to reconstruct the consensus 3D foreground binary, *B* (Methods).
4. Identify all 3D foreground voxels in *B* as initial (*t* = 0) coordinates of gradient descent. The coordinate of the *n*th foreground voxel is 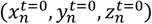.

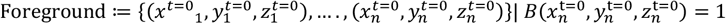
5. Apply gradient descent in 3D to iteratively advect the *n*th foreground point for a fixed number of total iterations, *t* = 1,2, … , *T*, with step size *η* to uncover its 3D medial-axis skeleton attractor

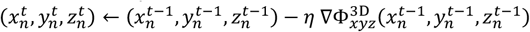

where the gradient field is normalized to unit length vectors, 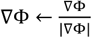, i.e. the parameter *η* defines the true voxel step size.
6. Cluster all points in the eroded space, at final advected coordinates (*t* = *T*) by spatial proximity, labeling each cluster with a unique positive integer object id, with all points part of the same cluster having the same id. This produces a 3D label image, *L*^*t*=*T*^.

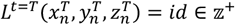

Finally, to obtain the 3D instance segmentation in the initial non-eroded space, generate the label image corresponding to the initial coordinates (*t* = 0), *L*^*t*=0^.

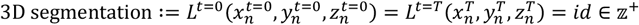

### u-Segment3D is a toolbox to create consensus 3D segmentations from 2D segmentations

u-Segment3D is comprised of robust implementations of each of the outlined steps in the general 2D-to-3D segmentation algorithm (Fig. 1f, Suppl. Movie 1). To accommodate variations in the quality of the 2D segmentations, each step exposes tunable parameters. These are detailed with rationale and tuning advice in Suppl. Table 1.

First, u-Segment3D offers a choice of distance transforms to tune the segmentation with respect to computation speed, accuracy, and compatibility with 2D model outputs (Extended Data Fig. 1). An object’s medial-axis skeleton is not unique^75,76^. u-Segment3D provides a choice of distance transforms categorized into two classes; ‘explicit’ (Extended Data Fig. 1b) or ‘implicit’ (Extended Data Fig. 1c).

Explicit transforms define attractor coordinates that are incorporated as boundary conditions in the computation (Methods). This ensures gradients are zero inside the attractor and enables stable convergence via gradient descent (Suppl. Movie 2). u-Segment3D implements single ‘point’ and multi ‘point set’ source attractors. The single point is the internal medial centroid, whose placement is adjustable by percentile-based thresholding of the cells’ Euclidean distance transform (EDT) (Extended Data Fig. 1d, Methods), extending the definition in Cellpose^5^. The point set is the 2D medial-axis skeleton of the cell shape (Methods). To compute the distance transforms, u-Segment3D considers two different partial differential equations (PDEs); the Eikonal equation, which gives the geodesic solution; and the Poisson equation, which gives the heat diffusion solution, as used in Cellpose^5^. The solution of the Eikonal equation, offers less smooth gradients but can be efficiently computed by the fast marching method^77^. The Poisson equation offers smoother gradients and thereby reduces oversegmentation but is slower to compute, being solved exactly using LU decomposition (Methods).

Implicit transforms specify the medial skeleton as ridges. Consequently, convergence to the attractor is unstable^6,76^ (Suppl. Movie 2), but it is more efficient, requiring only the solution of the PDEs without additional constraints. The Eikonal equation can then be solved using EDT, which is also an intermediary output of many 2D segmentation models^2,6^. Irrespective of the chosen distance transform, it is imperative for 2D-to-3D segmentation that the computational implementation guarantees a proper definition of the distance throughout the cell, regardless of cell shape. Iterative solutions of the PDE such as those implemented by Cellpose can fail to fully diffuse in shapes (Extended Data Fig. 1e), particularly with elongated and tortuous substructures (Extended Data Fig. 1f). Consequently, u-Segment3D relies on exact solutions of the PDEs. Gradients remain properly defined and faithfully capture the shape even for very long and branching structures (Extended Data Fig. 1f).

Second, the pipeline uses a content-based consensus averaging function, *F*, to fuse 2D image stacks, (Extended Data Fig. 2). 2D slice-by-slice segmentation may miss or under-/over-segment a cell across slices. Inspired by multiview image fusion^78,79^, u-Segment3D fuses multiple image stacks using linear inverse local variance weighting (Extended Data Fig. 2a, Methods). Using EDT as an example, segmentation errors across slices cause non-continuity such that erroneous pixels exhibit high local variance. Using inverse weighting the value of pixels from images with high-variance are down-weighted in the final fusion (Extended Data Fig. 2b). Increasing the size of the local pixel neighborhood enables correcting larger errors. For a 1×1×1 pixel neighborhood, *F* corresponds to the mean average fusion also used by Cellpose^5^. In this case no error correction is applied when fusing. With a 5×5×5 pixel neighborhood, binary thresholding on the fused EDT perfectly recovers the foreground nuclei without artifacts (Extended Data Fig. 2c).

Third, 2D-to-3D segmentation requires accurate implementations of gradient descent in 2D and 3D, (Extended Data Fig. 3). For downstream spatial proximity clustering gradient descent must propagate points of the same attractor together whilst retaining spatial compactness (Extended Data Fig. 3b). We verify our implementation using a synthetic 2D image of two objects, a circle within a doughnut (Extended Data Fig. 3a). Though simple, the object’s gradient field is complex with features typical of more nuanced morphologies such as local sinks and separating flows of opposite orientation. Running 100 iterations, u-Segment3D propagates points stably converging towards the two point attractors (Extended Data Fig. 3c), enabling perfect reconstruction of the original objects. In contrast, Cellpose’s implementation generates erroneous additional attractors: orphaning foreground points of the ring and an erroneous line attractor for the circle. Though minor in this example, inaccuracies in gradient descent compound with spatial clustering in the next step to detrimentally impact the final segmentation.

Fourth, to identify attractors in the gradient field the pipeline relies on robust spatial proximity clustering of the propagated points using connected component analysis, (Extended Data Fig. 4). Too many or too few clusters directly translate to over- and under- segmentation. With heterogeneous cell shapes, points do not converge to their attractors at equal speed. Running gradient descent to convergence for all cells is unnecessary and computationally expensive in 3D. Consequently, clustering must generalize to intermediate, irregular shapes represented by more uniform point densities. Density-based clustering methods such as the adaptive local histogram thresholding used by Cellpose^5^ are unsuitable. These implicitly assume and find all local point-like hotspots as attractors. This leads to catastrophic failure, generating grossly fragmented cells.

To overcome this, u-Segment3D exploits the fact that foreground coordinates are on an image grid (Extended Data Fig. 4a). The final floating-point advected coordinates are rasterized by integer flooring (step i). A count of the number of coordinates in each voxel is tabulated (step ii) and smoothed with a Gaussian filter of *σ* to build an approximate kernel density heatmap, *ρ* (step iii). *ρ* is sparse, enabling clusters to be identified using a global threshold, *mean*(*ρ*) + *k* · *std*(*ρ*) with tunable *k*. Connected component analysis then labels all spatially separated regions irrespective of shape and density with unique ids (step iv). The final segmentation is generated by indexing the final advected coordinates of foreground voxels in this label image, and transferring the labeling to the initial coordinates. *ρ* enables probabilistic cluster identification. By increasing *σ* u-Segment3D can ‘fuzzy’ link erroneously split clusters, equivalent to merging segmentations in the final 3D. We validated our implementation by reconstructing the 2D cell segmentation as we propagate foreground coordinates along the gradients of the geodesic centroid distance transform (Extended Data Fig. 4b i,ii). We correctly obtain the equivalent segmentation of applying connected component analysis to the foreground binary at iteration 0. As iterations increase, and attractors emerge, the segmented cells converge on the true number (Extended Data Fig. 4b iii, top). Correspondingly, segmentation quality, measured by the intersection- over-union (IoU) and F1 score, increases to 1 (Extended Data Fig. 4b iii, bottom). These observations also translate to elongated, touching cells (Extended Data Fig. 4c). Moreover, only our clustering recovers the number of clusters present in the final coordinates propagated by either Cellpose or u-Segment3D’s gradient descent for the synthetic image of a circle within a doughnut (Extended Data Fig. 3d, 4d). In contrast Cellpose’s clustering breaks up what should be single clusters (Extended Data Fig. 3d). This negatively impacts Cellpose’s 3D segmentation performance, particularly for low signal-to-noise ratio or out-of- distribution cells (Extended Data Fig. 4e,f). Whereas Cellpose’s gradient descent and clustering grossly oversegments and fractures individual cells, u-Segment3D’s gradient descent and connected component clustering recovers complete cell segmentations when applied to reparse the same predicted foreground binary and 3D gradients (Extended Data Fig. 4e,f, Suppl. Movie 3).

To maximize the utility of pretrained 2D models, u-Segment3D further implements preprocessing and postprocessing modules. The image to segment may not reflect the quality, acquisition, noise distribution and modality of the training dataset a segmentation model was trained on. Preprocessing can help transform input images to improve performance in the 2D segmentation of orthoview slices^27,62^. While dataset- and model-specific, the following order of processing steps has proven broadly applicable: intensity normalization, none or any combination of denoising, deconvolution and ridge feature enhancement, and uneven illumination correction with optional gamma correction (Methods). Postprocessing steps are implemented; to filter out implausible segmentations by object size and consistency with the reconstructed 3D gradients (as in Cellpose^5^), to refine segmentations by spatial-connectivity aware label diffusion to better adhere to cell boundaries within a guide image; and guided filtering to recover missing or intricate subcellular details. None of these postprocessing procedures require further 3D training (Methods).

### Smoothing of reconstructed gradients and suppressed gradient descent are essential for 1D-to-2D segmentation

To understand how the different components of the 2D-to-3D algorithm may impact 3D segmentation, we first empirically investigated 1D-to-2D reconstruction of single cell shapes from the Cellpose^5^ and Omnipose^6^ training datasets, which can be intuitively visualized (Extended Data Fig. 5). We first examined the approximation of 2D gradients using 1D gradients (Extended Data Fig. 5a, b, Methods). Across single cells representing spherical, convex, branched and vessel morphologies, we found that Gaussian filtering of the reconstructed 2D gradients was essential to recover the original 2D shapes (Extended Data Fig. 5b). Without smoothing, the 2D gradients specified by the raw 1D x-, y-gradients (Methods), do not capture the 2D context and specify a single fixed-point attractor per cell. Smoothing with increasing *σ*, trajectories were regularized to converge towards single points. Smoothing effectively conformalizes the cell shape, shifting its 2D centroid towards the centroid of its convex hull. Unfortunately, for concave shapes, the new attractor may lie outside the cell. To examine the implications of this with crowded cells, we performed the same 1D-to-2D reconstruction considering full images. For a Cellpose example (90 cells), the reconstructed 2D gradients smoothed by *σ* = 1 contains more than one attractor per cell (Extended Data Fig. 6a). Consequently, after 50 iterations of gradient descent we oversegment (143 cells). Nevertheless, the shape reconstruction is good (*F*1 = 0.77, IoU=0.91). Moreover, for fragmented cells, the number of fragments per cell was low, and were largely small fragments. Increasing *σ* regularizes the reconstructed gradients. For *σ* = 5, segmentation (93 cells, F1=0.94, IoU=0.93) is on-par with ideal 2D gradients (93 cells, F1=0.98, IoU=1.00). Beyond *σ* > 5, gradients interact across neighboring cells causing undersegmentation, as well as decreased IoU and F1 performance (Extended Data Fig. 6a iv). For long and thin tubular structures in the Omnipose example (86 cells), increasing *σ* shifts attractor centroids into neighboring cells. Thus increasing *σ* improved F1 but also decreased IoU, with an optimal balance at *σ* = 3 (F1=0.49, IoU=0.60) (Extended Data Fig. 6b iv). Due to the sensitivity to cell size and shape, tuning *σ* is impractical. The aim of gradient descent is to separate adjacent cells by clustering points towards their medial-axis skeleton. Iterations beyond this point are unnecessary. Depending on the distance transform (Suppl. Movie 2) they can cause fragmented clusters and thus oversegmentation. To prevent excessive iterations without changing the preset iteration maximum, we define a variable stepsize, 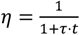, decaying for higher iterations, where *t* denotes the iteration and τ an adjustable decay rate^6^. Applying the suppressed gradient descent^6^ with *σ* = 1 perfectly reconstructed the 2D shapes (86 cells, F1=1.00, IoU=1.00) (Extended Data Fig. 6c).

### Translating gradient smoothing and suppressed gradient descent to the case of 2D-to-3D segmentation

To test if insights from the 1D-to-2D segmentation translate to 2D-to-3D segmentation we conducted the analogous reconstruction experiment for single 3D cells (Extended Data Fig. 7a). These cells were segmented from high-resolution microscopy images as surface meshes with u-Shape3D^73^ then voxelized to binary volumes using u-Unwrap3D^72^. Similar to the 1D-to-2D case, the 2D geodesic centroid distance transform (Methods) was computed slice-by-slice in orthogonal x-y, x-z, y-z stacks, treating spatially contiguous 2D regions as unique ‘cells’. The 3D gradients were then reconstructed by averaging (*F* with 1×1×1 pixel neighborhood). Different, however, is that we additionally incorporate momentum into the suppressed gradient descent to expedite convergence (Methods). 3D cells were selected to represent a spectrum of morphologies ranging from pseudo-spherical, to pseudo-convex and branched, and with different types of surface protrusions (Extended Data Fig. 7b). Applying suppressed gradient descent (τ = 0.1) for 200 iterations, we found similar results as 1D-to-2D, with 3D cell examples of pseudo-spherical, pseudo-convex and branched morphologies (Extended Data Fig. 7b, Suppl. Movie 4). Gaussian smoothing aids regularization and increasing *σ* ensures convergence to a single cell, even for a highly branched cell with filopodia (*σ* = 15). The same cell was fragmented into several regions at branch junctions at lower *σ* = 1. However, we can also keep *σ* = 1 and increase τ = 0.5 to still recover perfect construction of the branched cell.

In summary, these experiments underline the necessity of Gaussian filtering to smooth the reconstructed 3D gradients, and the use of a suppressed gradient descent. These are among the algorithmic key differences to Cellpose that allow u-Segment3D to segment cells of heterogeneous morphologies.

### u-Segment3D reconstructs 3D objects from their orthogonal 2D slices

We assembled 10 published 3D datasets with dense segmentation labels and 1 additional zebrafish macrophage dataset (Suppl. Table 2). This latter dataset was curated in-house by combining thresholding and connected component analysis with u-Segment3D consensus segmentations^80^. DeepVesselNet^10^ is a dataset of simulated binary vasculature networks. To identify disconnected subnetworks as unique ‘cells’, we applied connected component analysis. The total number of cells across all datasets was 73,777. For each cell, we extracted 8 morphological features (Fig. 2a, Methods), chosen to assess cell size (total number of voxels), the extent of elongation (stretch factor) and the topological complexity (# of skeleton nodes). To visualize in 2D the morphological diversity and variation in cell numbers across datasets, we applied UMAP^81^ to the normalized features (Fig. 2b, Methods). Two plant datasets: Arabidopsis (CAM) (24,439) and Ovules (37,027) contributed the majority of cells (83%) and dominate the UMAP. The assembled datasets capture commonly found 3D morphotypes encountered in tissue including thin, complex vessel-like networks (Region 1), pseudo-spherical (Regions 2-4), irregular (Region 5), and tubular or branched (Region 6). Using the per dataset median UMAP coordinate, and colored UMAP by stretch factor, # skeleton nodes and volume, we broadly divide the 11 datasets into three super-morphotypes: complex networks (DeepVesselNet), irregular/branched (Zebrafish macrophages/Platynereis ISH nuclei/MedMNIST3D/Lateral Root Primordia), and convex (*C. Elegans* embryo/mouse organoid/mouse skull nuclei/*Platynereis* nuclei/Arabidopsis (CAM)/Ovules).

**Figure 2.**
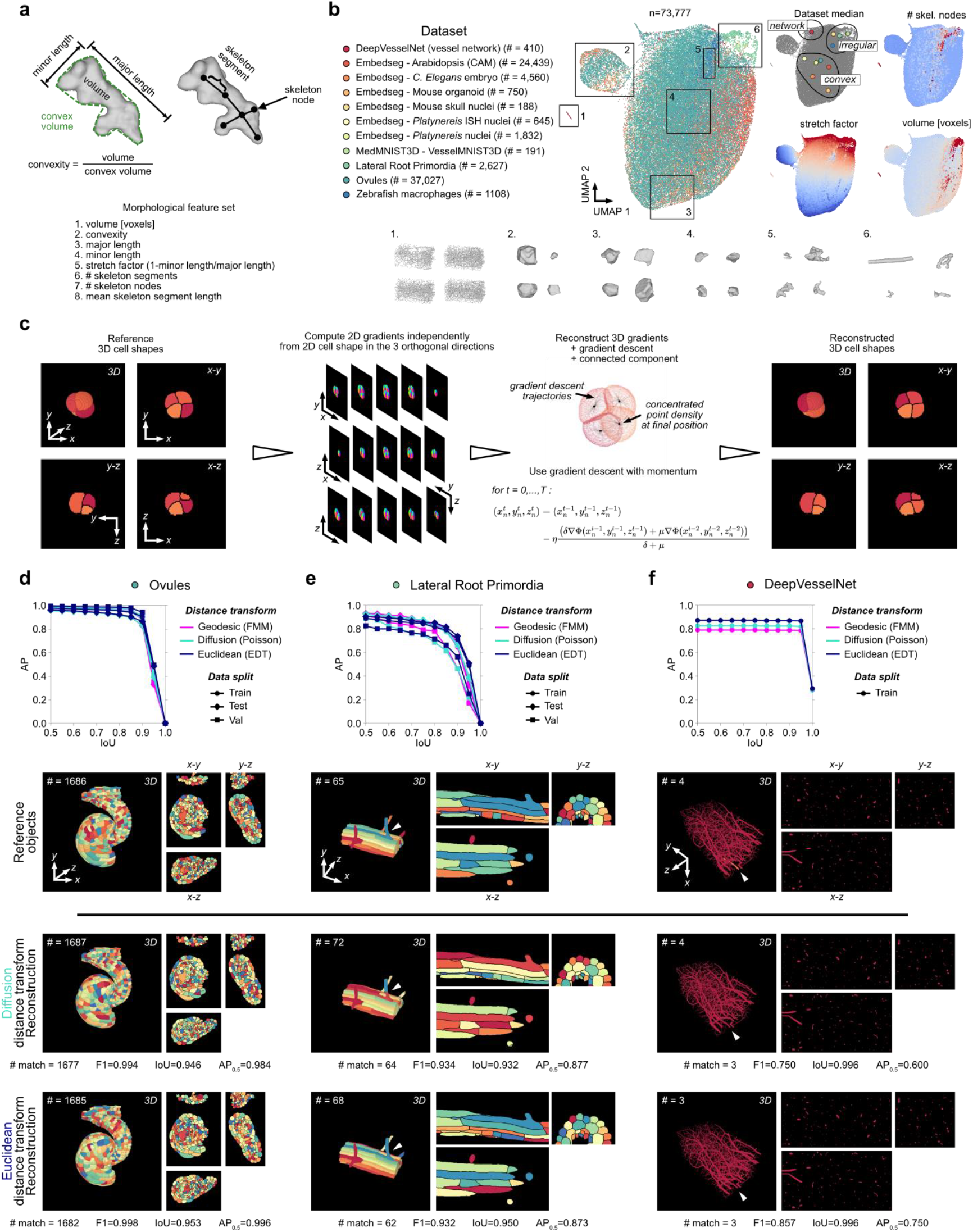
u-Segment3D reconstructs 3D objects from their 2D slices in orthogonal x-y, x-z, y-z views. **a**,**b)** Characterization of test dataset. Illustration of the 8 computed geometrical and topological features to describe the shape complexity of test data (a). UMAP embedding of individual cells from 11 datasets covering the full spectrum of morphological complexity from convex-spherical to branching to networks (b). Left: colormap of individual dataset and total number of uniquely labelled cells in each dataset. Middle: UMAP, each point is a cell, color-coded by their dataset. Right: Median UMAP coordinate of each dataset (top left) and heatmap of three features representing the extent of branching (total number of skeleton nodes, top right), the extent of elongation (stretch factor = 1 – minor length/major length) and their image size (total number of voxels). **c)** Illustration of the experimental workflow to compute 2D slice-by-slice distance transforms in orthogonal directions using a 3D reference shape. These 2D transforms are then integrated by u-Segment3D to reconstruct a 3D cell shape. Comparison of reference and reconstruction is used to assess performance of the 2D-to-3D reconstruction.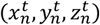 denotes the coordinate of the *n*th foreground voxel at iteration number *t, η* the step size, *δ*, *μ* the weighting of current and previous gradients where *δ* ≥ *μ*. We set *δ* = 1 and *μ* is momentum. **d)** Reconstruction performance measured by the average precision (AP) curve (Methods) for the Ovules dataset using three different 2D distance transforms. From top to bottom: average precision vs intersection over union (IoU) curve; 3D rendering of reference vs reconstructions based on point-based diffusion distance transform and skeleton-based Euclidean distance transform. Shown are the reconstructed 3D objects and their respective mid-slices in the three orthogonal views. **e), f)** Same as d) for the Lateral Root Primordia dataset containing instances of branching morphology and DeepVesselNet representing complex, thin network morphologies. Individual cells are uniquely colored but are not color- matched with the reference object.

For all images in each dataset, we use the provided segmentation labels as reference 3D objects, and tested the reconstruction of each 3D object from their 2D slice projections (Fig. 2c). All 3D objects were scanned slice-by-slice in x-y, x-z, y-z views, with each 2D contiguous region in a 2D slice treated as a unique ‘cell’. For each 2D ‘cell’, the 2D gradients were computed and used to assemble the 3D gradients for reconstruction of the 3D objects by 3D gradient descent and connected component analysis. This experimental setup allowed us to assess 3D shape reconstruction performance based on 2D gradients. Three different 2D distance transforms were tested: Poisson diffusion centroid as an example of an explicit transform and used in Cellpose^5^; Euclidean distance transform as an example of an implicit transform and used within models like Omnipose^6^, StarDist^2^; and geodesic centroid as a second example of an implicit transform, but computed differently (Methods). For all datasets, the number of iterations in the total gradient descent was fixed at 250, and reference 3D objects were resized to isotropic voxels with nearest-neighbor interpolation (Suppl. Table. 2). Temporal decay τ was the only parameter adjusted for each transform and dataset (Suppl. Table 3). Postprocessing was applied to remove cells of less than 15 voxels. Reconstructed 3D objects were evaluated using the average precision (AP) and F1 curve. AP and F1 are two aggregate scores of precision and recall, with AP equally weighting true positives, false positives and false negatives (Methods). Their curves compute the score between reference and predicted shapes as the overlap cutoff (IoU) for a valid match is increased from 0.5 to 1.0 (perfect overlap) (Methods). We use the notation AP_0.5_, *F*1_0.5_ to denote AP, F1 with IoU cutoff=0.5 henceforth. For perfect reconstruction, AP=1, F1=1 at all IoU. In practice, due to limited numerical accuracy, AP, F1 always drops to 0 above an IoU cutoff.

For each of the super-morphotypes, we first analyzed the dataset with the highest number of cells: Ovules for convex (Fig. 2d), Lateral Root Primordia (LRP) for irregular (Fig. 2e) and DeepVesselNet for networks (Fig. 2f) (Suppl. Movie 5). We find near-perfect reconstruction across all distance transforms, morphotypes and data splits, qualitatively and quantitatively: Ovules, AP0.5 ≈1.0, LRP, AP0.5 ≥ 0.8, and DeepVesselNet, AP_0_.5 ≥ 0.8. As expected, increased τ was necessary for thinner, branching cells: using EDT, τ =0.5 for Ovules and LRP, τ =2.0 for DeepVesselNet. These results were reflected in the other 8 datasets (Extended Data Fig. 8), with AP_0.5_ ≥ 0.75. F1 curves support the same conclusion as AP0.5 across all datasets (Extended Data Fig. 9) but with higher scores, Ovules, F10.5 ≈ 1.0, LRP, F10.5 ≥ 0.9, and DeepVesselNet, F10.5 ≥ 0.85. Henceforth, we primarily plot AP for better visualization. Moreover, IoU was high, with both performance metrics decaying prominently only for IoU ≥ 0.85 and for many, IoU ≥ 0.95. Notably, IoU > 0.8 masks are near-indistinguishable from the reference by eye^6,82^. We also compared reconstruction by stitching using CellComposer^50^ and CellStitch^71^. Whereas u-Segment3D demonstrates universal applicability with high performance across all datasets and distance transforms, CellStitch and CellComposer were inconsistent, with varying performance within the same morphotype, for convex, ranging from AP_0.5_ ≈ 0.60 (Mouse organoid) to 1.0 (Ovules). Moreover, both are inapplicable to branched morphologies (AP_0.5_ ≈ 0.4 (VesselMNIST3D), and AP_0.5_ ≈ 0.0 (DeepVesselNet)).

As we fix the foreground, the performance gap to an ideal AP_0.5_ = 1 of u-Segment3D largely reflects errors in the 3D objects, a consequence of artifacts in the reference segmentation labels provided for each dataset. Gradient descent 2D-to-3D aggregation requires a spatially contiguous path in 3D. Therefore, each 3D object must only be a single spatial component. We did not check for nor enforce this to better reflect the nuances of real-life segmentation annotation quality. As a first check, balanced dataset splits should exhibit the same reconstruction performance. However, in LRP, known to contain labelling errors^6,42^, the AP curve of all three transforms on the validation (val) split were notably worse. In only 6/11 datasets (Ovule/Arabidopsis(CAM)/*C*.*Elegans*/mouse organoid/Platynereis nuclei/vesselMNIST3D) did the best distance transform achieve a perfect AP_0.5_ =1.00. Notably, these were majority convex-shaped or had unambiguous cell edges and minimal background. As a second check, across all datasets, zebrafish macrophages, had the lowest reconstruction performance (AP_0_.5 = 0.8 with Poisson distance transform) and these segmentations were also constructed with the least proofreading^80^. To segment the extremely branched macrophages, we generated the segmentations by automatically replacing u-Segment3D consensus segmentations of Cellpose 2D outputs with connected component segmentations using hard- coded rules^80^. Consequently there are small and multi-component ‘cells’. For DeepVesselNet, the reconstruction error is over-estimated quantitatively. The average number of subnetworks is 3, thus our results reflect 1 misidentified small subnetwork on average. Qualitatively, there is no noticeable difference in coverage (Fig. 2f), thus any errors are likely data resolution-related, for example two subnetworks separated by a small gap were joined (Fig. 2f, white arrow), or size filtering removed a small subnetwork or a connecting segment between two subnetworks. In comparison, stitching is extremely sensitive to discretization, thereby only generating many 3D vessel subnetworks, resulting in AP_0.5_ = 0.0.

As pointed out by the Omnipose paper^6^, there are discrepancies between different distance transforms for different morphotypes. The two explicit transforms with point-source attractors, Poisson and geodesic, differed minimally. Both outperformed EDT on convex morphologies, most evidently in the LRP val (Fig. 2e, Extended Data Fig 9b), mouse skull nuclei test (Extended Data Fig 8d, 9g), Platynereis ISH nuclei test (Extended Data Fig. 8e, 9h) and zebrafish macrophages (Extended Data Fig 8h, 9k) datasets. This is primarily due to the increased stability of explicit transforms. However, EDT was superior for thin and complex vasculature networks (Fig. 2f, Extended Data Fig 9c, DeepVesselNet), better described by skeletal-based representations, and minimizing the distance each point propagates. Overall, the quantitative difference was small (<0.5 difference in AP_0.5_) and not as dramatic as suggested by Omnipose^6^. This is because under gradient descent the medial axis 3D skeleton is always an intermediate structure when converging towards a centroid attractor (Suppl. Movie 4). Qualitatively, we visualized both the diffusion and EDT reconstruction on exemplars from Ovules, LRP and DeepVesselNet. Despite similar F1 and IoU, only the EDT fully reconstructed all branching cells. Diffusion fragmented the cell with the longest branch (Fig. 2e, white arrows) into two ‘cells’. Importantly, the fragments are standalone and not erroneously part of neighboring cells.

In summary, u-Segment3D is universally applicable across all morphotypes and empirically achieves near-perfect, consistent 3D shape reconstruction from 2D slice projections in orthogonal views. In the best case, it is perfect. In the worst case, a subset of branching cells might be decomposed into a few standalone segments that could be reassembled. Our results indicate that generally, explicit distance transforms perform more optimally, however these are more computationally expensive than implicit transforms.

### u-Segment3D generates consensus 3D segmentation from any orthogonal 2D slice-by-slice instance segmentation

2D segmentation models either (i) already predict a suitable distance transform or 2D gradients (c.f. Extended Data Fig. 2, Methods), e.g. Cellpose^5^, or (ii) provide 2D instance segmentation masks from which distance transforms may be computed. u-Segment3D accounts for both cases (Fig. 3a). In the former, predicted 2D gradients are directly used to generate the 3D segmentation (the direct method). In the latter, users choose a 2D distance transform to compute the necessary 2D gradients from the 2D segmentation masks, (the indirect method). We demonstrate both methods using pretrained generalist Cellpose models. Unlike the ideal 2D projections derived from given 3D shapes, the morphology of the reconstructed 3D foreground binary is an additional determinant of the overall performance. If the foreground does not provide a contiguous path for gradient descent, the resulting segmentation will be fragmented, even with correct gradients.

**Figure 3.**
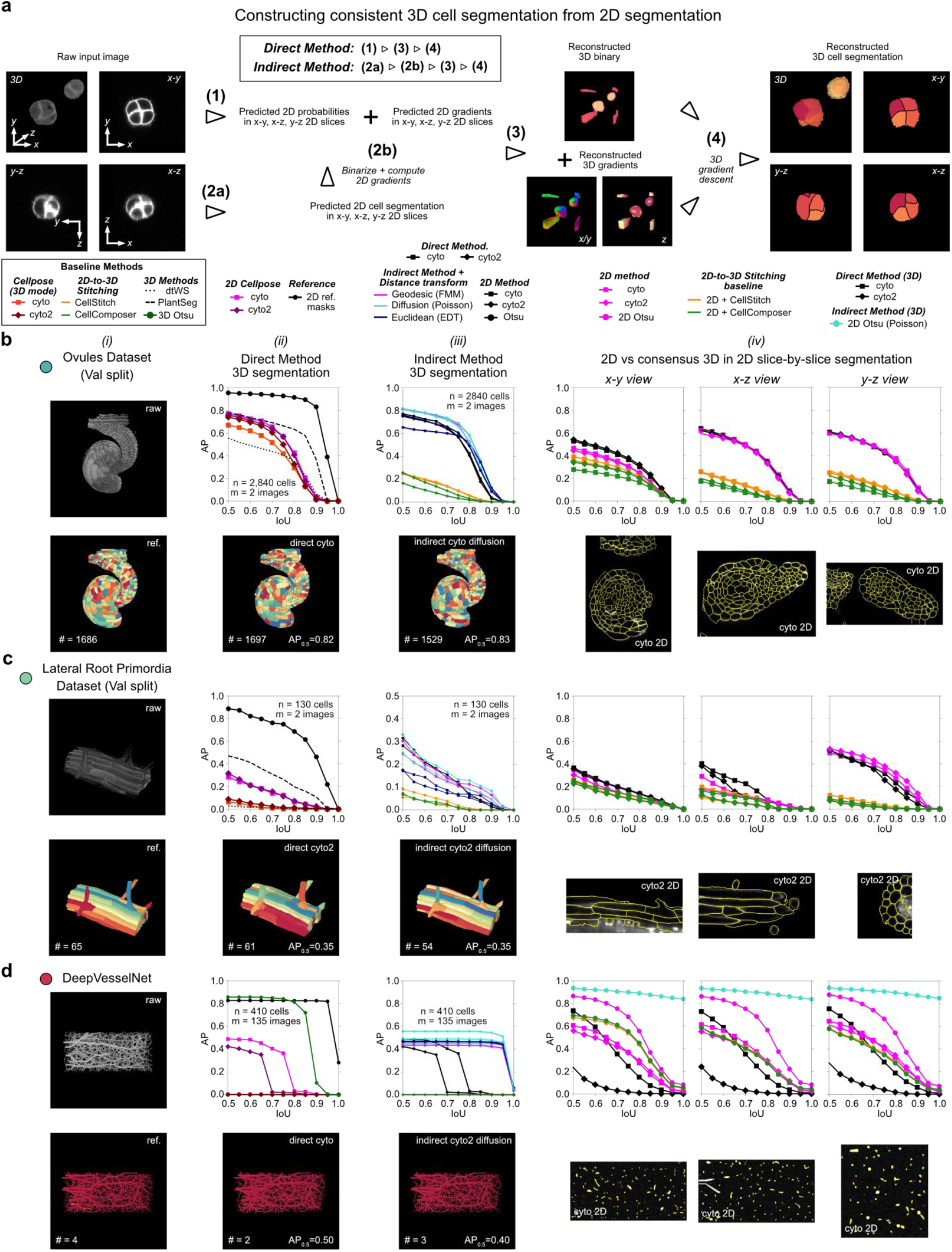
Consensus segmentation of 3D real datasets using pretrained Cellpose2D models applied to orthogonal x-y, x-z, y-z views. **a)** Illustration of the two workflows that can be executed in u-Segment3D to generate 3D cell segmentations. The direct method (steps 1,3,4) allows direct utilization of predicted spatial gradients and cell probability maps output by some 2D segmentation models without generating instance segmentation masks. The indirect method (steps 2a, 2b, 3, 4) operates on instance segmentation masks from x-y, x-z, and y-z and computes spatial gradients from masks based on the user chosen 2D distance transform (see also Fig. 2c). Image panels for step 3 show the Cellpose outputs of the input data which predicts extraneous ‘cells’ at the image border. These are removed during postprocessing and therefore not present in the final segmentations of step 4. **b)** u-Segment3D performance on the Ovules dataset (validation (val) split, n=2840 cells, m=2 volumes) using pretrained Cellpose 2D outputs (i) 3D rendering of raw image (top) and reference 3D labels (bottom). (ii) Average precision (AP) curve for the direct method using pretrained Cellpose 2D cyto or cyto2 models relative to the AP curve of the best reconstruction from ideal 2D slices in Fig. 2d (top). Also shown are AP curves for segmentations by Cellpose 3D. 3D rendering of the segmentation using the Cellpose 2D cyto model (bottom). (iii) Average precision (AP) curve for the indirect method using the 2D segmentation of pretrained Cellpose 2D cyto or cyto2 models and different 2D distance transforms relative to the AP curve of the corresponding direct method constructed 3D segmentation, and alternative 2D-to-3D stitching methods; CellStitch and CellComposer (top). 3D rendering of the segmentation using the Cellpose 2D cyto model and the centroid diffusion distance transform – the highest-performing combination using the indirect method (bottom). (iv) Average precision (AP) curve of the 2D slice segmentations of direct method consensus 3D segmentation (i.e. 3D-to-2D segmentation, black lines), and 2D-to-3D stitched segmentations using CellStitch (orange) or CellComposer (green), compared to Cellpose 2D model independent segmentations of each 2D slice (magenta lines) in each orthogonal view, x-y, x-z, y- z from left-to-right. Representative 2D segmentations from the highest performing Cellpose cyto model are shown below the AP curves. The consensus 3D segmentation consistently improves 2D segmentation, showing u-Segment3D leverages complementary information from individual orthogonal views. **c), d)** Same as b) for the Lateral Root Primordia (validation (val) split, n=130 cells, m=2 volumes) and DeepVesselNet (n=410 network components, m=135 images) dataset with representative visualizations from the best performing combination of Cellpose model and distance transform. For d) we additionally evaluated the performance of binary Otsu thresholding as a native 3D baseline, and as a 2D baseline with the indirect u- Segment3D method (explicit diffusion transform, circle marker) (turquoise), cellstitch (orange) and cellcomposer (green).

When working with pretrained Cellpose 2D models two model parameters crucially influenced performance: (i) the diameter defining the expected cell object size and (ii) the cell probability threshold to determine the foreground binary. Cellpose 2D models enable ‘optimal’ diameter prediction using a pretrained regression model. However, this assumes one size fits all. An image can contain objects of different scales, e.g. cell body vs cell nuclei, cells within an embryo vs the overall embryo shape. Moreover, a trained model cannot guarantee generalization to out-of-sample datasets or consistency across sequential 2D slices. When we examined cell probability and gradients on cross-sections of the LRP dataset predicted by Cellpose, we found seemingly similar results over a broad range of ‘diameter’ settings (Extended Data Fig.10a-c). To set the ‘diameter’ objectively without training, we developed an alternative tuning method based on examining the model’s self-confidence. u-Segment3D runs Cellpose over a test diameter range to compute a ‘contrast score’ per diameter using the local pixel variance within the predicted gradients and cell probability (Extended Data Fig. 11a, Methods). The resulting contrast function uncovers all salient object scales as local maxima, serving as a tuning guide. Cellpose models are trained using a mean diameter of 30 pixels and documented to perform best for diameter=15-45 pixels. Therefore, based on the peak of the contrast function, images are resized in preprocessing (Suppl. Table 1) to match Cellpose’s optimal operating diameter range. For automated operation, u-Segment3D selects the diameter with maximum contrast as optimal. If multiple peaks are present, the optimal diameter can be biased to favor alternative maxima by adjusting the size of the considered pixel neighborhood used to compute the contrast score (Extended Data Fig. 11, Suppl. Table 1) or by constraining the diameter range. In 3D, the cross-sectional appearance of an object often has different aspect ratios and size, even if the image is resized to isotropic voxels. We apply our tuning to set the optimal diameter in each of the x-y, x-z, and y-z views using a representative 2D slice (Methods). As validation, the Cellpose predicted diameter coincides with the predicted maxima of our method on the same 2D image (Extended Data Fig. 10a-c). Moreover, the direct method 3D segmentation using our method (AP_0.5_=0.28) is comparable to using Cellpose’s method (AP_0.5_=0.23), if not better (Extended Data Fig. 10d).

For thresholding the cell probability u-Segment3D uses multi-threshold Otsu to statistically determine a finite choice of thresholds (Methods). Cellpose does not provide automated means. We can use flooring to round thresholds to the nearest decimal point or select a lower threshold to strike a balance between segmentation accuracy and ensuring contiguous foreground regions for gradient descent. This works excellently for both 2D Cellpose and for 3D reconstructed cell probabilities. Given the problems we found with Cellpose’s gradient descent (Extended Data Fig. 3), and spatial clustering (Extended Data Fig. 4), henceforth we always use u- Segment3D’s equivalent to generate segmentations. Thus, we refer to Cellpose 2D outputs only as the predicted gradients and cell probability, whereas Cellpose 2D segmentation refers to the cell masks after applying to Cellpose 2D outputs u-Segment3D’s statistical binary thresholding, suppressed gradient descent with momentum and connected component analysis.

Using our tuning and parsing of Cellpose, we compared the direct and indirect method of u-Segment3D on 9/11 datasets (see Suppl. Table. 4 for parameter details). The zebrafish macrophage dataset was excluded, as labels were derived from u-Segment3D; so was VesselMNIST3D, which only contains binary masks. We also considered two pretrained Cellpose models, ‘cyto’ and ‘cyto2’, both generalist models but trained on different datasets, to assess how different 2D models translate to different 3D segmentation performance. The direct result was assessed relative to Cellpose 3D and native 3D segmentation baselines. For the latter, we used an unsupervised 3D watershed as the lower bound, and dataset-specific native 3D segmentation model as the upper bound (Methods). The indirect result was compared to the stitching methods, CellComposer and CellStitch. To minimize data leakage, we applied models to only the validation or test splits when available.

On Ovules validation split (n=2,840 cells, m=2 images, Fig. 3b i), we found excellent performance with both the ‘cyto’ and ‘cyto2’ models and the direct method, surpassing the watershed and comparable to the specialized native pretrained PlantSeg3D^7^ models (Fig. 3b ii, AP_50_ ≈ 0.80 for both). This was expected as the cells are convex and delineated by high-contrast edges. This is also evidenced by good performance of running Cellpose 3D on the same preprocessed input. Cellpose 3D has no automatic diameter tuning and uses only one diameter across all views. To perform a fair comparison without modifying its source code, we accounted for the oversegmentation tendency of Cellpose 3D and used the maximum of u-Segment3D inferred diameters. We also ran Cellpose 3D twice, the first to obtain 3D cell probabilities to compute the equivalent Otsu thresholds and the second to obtain final segmentations using the same postprocessing parameters (Methods). As expected from algorithm design, for the same model, u-Segment3D consistently outperforms Cellpose 3D. Most impressively, u-Segment3D even boosted the ‘cyto’ model from *AP*_0.5_ = 0.7 to *AP*_0.5_ = 0.8 to be on par with ‘cyto2’. The same was true for the test split (n=10,328 cells, m=7 images, Extended Data Fig. 12). Again u-Segment3D raised the average precision of ‘cyto’ from *AP*_0.5_ = 0.65 to *AP*_0.5_ = 0.7. Compared to the best 3D reconstruction with ideal 2D segmentations (black line, circle marker, (AP_50_ =1.0), however, there remains a noticeable gap of 0.2. Interestingly, the indirect method with either the geodesic (magenta colored) or diffusion (cyan colored) distance transforms for both models was better quantitatively than the direct method (Fig. 3b iii, *AP*_50_ > 0.80). This is likely due to better cell boundary delineation from aggregating on the hard-thresholded 2D segmentations. However, the total number of cells predicted decreased (reference=1686, direct=1697, indirect=1529). In contrast, CellComposer and CellStitch are severely impeded by non-ideal Cellpose 2D segmentations. The stitched 3D segmentation was worse than 3D watershed (AP_0.5_ = 0.2-0.3), a > 0.5 decrease from using ideal 2D slices (Fig. 2d). Lastly, we tested whether the 2D orthoslices of the direct 3D segmentation (black lines) retain the accuracy of 2D segmentations. We compared the native 2D slice-by-slice segmentations in x-y, x-z, and y-z views (magenta lines) to the corresponding cross-sections of the 2D-3D consensus segmentation by u-Segment3D as translated by the direct method (Fig. 3b iv). Not only does u-Segment3D preserve the 2D segmentation but even improved it in x-y view for both models (AP_0.5_ = 0.45 to 0.55). Meanwhile, CellComposer and CellStitch underperform with a strong bias against the x-z and y-z views. This demonstrates the true, consensus integration of complementary information from orthogonal predictions by u-Segment3D.

The segmentation of the LRP data set is more challenging as it contains a mixture of both compact and elongated/branching cells with weakly-defined edges (Fig. 3c). Unsurprisingly, direct u-Segment3D segmentation with both models on the validation split was substantially lower (AP_50_ ≈ 0.30 for both) than that from ideal 2D segmentations (AP_50_ ≈ 0.90). Nevertheless, u-Segment3D significantly outperforms Cellpose 3D and watershed in both the validation and test splits (Extended Data Fig. 12, AP_50_ ≈ 0.05 for both; improving Cellpose 3D cyto (AP_50_ = 0.18 to 0.37) and cyto2 (AP_50_ = 0.19 to 0.40)). As with Ovules, AP_50_ was lower than PlantSeg3D model. However, this AP_50_ with pretrained Cellpose models with u-Segment3D was comparable to an Omnipose model (plant-omni) trained natively in 3D specifically on LRP^6^. We asked whether a specialist Cellpose 2D model (plant-cp) pretrained on 2D slices of LRP^6^ would improve 3D performance. Performing a like-for-like evaluation (Methods), we find 2D plant-cp using indirect u-Segment3D and any distance transform outperformed 3D-trained plant-omni and generalist Cellpose 2D models for both validation (AP_0.5_=0.50, Extended Data Fig. 12c) and test (AP_0.5_=0.50-0.56) splits, comparable to PlantSeg3D (Fig. 3c, Extended Data Fig. 12c,d). Unexpectedly, plant-omni and plant-cp (using Cellpose 3D) were only as good as pretrained cyto2 and direct u-Segment3D in both splits. Closer inspection revealed that though plant-omni looks excellent in 3D renderings, segmentations were incomplete when viewed cross-sectionally. Additionally, plant-omni oversegments despite additional size filtering (Methods). These results highlight the robustness of u-Segment3D and verify that the framework translates improved 2D segmentation models into better 3D segmentations, on-par or better than natively 3D trained models. Again, direct and indirect u- Segment3D segmentations were comparable in AP_50_, and outperformed CellStitch and CellComposer equivalents. Indirect segmentations were better in IoU, with a slightly slower drop-off (Fig. 3c iii). Also for LRP, u-Segment3D exploits complementary information from all orthogonal views: the 2D slice segmentations of the consensus 3D segmentation is marginally worse than Cellpose 2D segmentations in y-z AP curve, but improves in x-y and x-z AP curve performance for both ‘cyto’ and ‘cyto2’ models (Fig. 3c iv). In contrast, despite the y-z view being most convex-shaped and having the best 2D segmentation, it was worst after stitching using CellStitch and CellComposer.

DeepVesselNet, comprised of thin, complex vasculature networks represents the biggest challenge for 2D- to-3D segmentation (Fig. 3d). Both pretrained Cellpose models predict segmentations uniformly larger than the actual vessel radii in 2D slices. Hence, we additionally uniformly eroded consensus 3D segmentations to obtain the final segmentation (Suppl. Table 4). Nevertheless, there was a clear difference between the two models. Using direct segmentation, ‘cyto’ (AP_0.5_ = 0.5, IoU drop-off≈0.75) outperforms ‘cyto2’ (AP_0.5_ = 0.4, IoU drop-off≈0.65) (Fig. 3d ii). Without suppressed gradient descent, Cellpose 3D grossly oversegments (AP_0.5_ = 0). Again, direct and indirect u-Segment3D segmentations were on-par in AP_50_. However, the indirect method is superior in IoU, with drop-off extending to 0.95 with similar AP curves across all distance transforms (Fig. 3d iii). As with ideal slices (Fig. 2f), CellComposer and CellStitch oversegment (AP_0.5_ = 0). Comparing 2D segmentation performance, the direct 3D aggregated cyto outperforms individual 2D segmentations in AP_0.5_ but exhibits faster IoU drop-off (Fig. 3d iv). The direct aggregated cyto2 was significantly worse than its 2D counterpart. This is likely due to the 3D erosion postprocessing removing too many small 2D segmentations in slices. Since the background appeared homogeneous in this dataset, we additionally tested 2D binary Otsu thresholding and connected component analysis instance masks. This yielded the highest AP_0.5_ 2D segmentations in all orthogonal views (Fig. 3d iv, magenta circle lines). Applying u-Segment3D, we achieve the highest AP_0.5_ 3D segmentation (AP_0.5_ ≈ 0.6) with a 2D method (Fig. 3d iii, turquoise circle line) but this was lower than native 3D binary Otsu thresholding and connected component analysis, (AP_0.5_ ≈ 0.8, Fig. 3d (ii), green circle line). Impressively, the consensus u-Segment3D (Fig. 3d iv, turquoise circle line) significantly improves upon the original 2D Otsu segmentations (Fig. 3d iv, turquoise circle line), unlike CellComposer and CellStitch. Therefore, we attribute the discrepancy in performance to discretization, whereby 3 orthoviews are insufficient to fully capture the 3D branching directions.

Altogether, these analyses demonstrate the robust implementation and universal applicability of u- Segment3D to real datasets of different super-morphotypes. We also showed how u-Segment3D can be applied to any 2D segmentation method generating pixel-wise distance transforms and gradients or instance masks using the direct or indirect methods, respectively, with similar AP_0.5_. Thus, we applied only the direct method on remainder EmbedSeg^15^ datasets with 3D watershed and specialist-trained EmbedSeg3D native 3D segmentation baselines (Extended Data Fig. 13, Methods). Except for Arabidopsis (CAM) (best AP_0.5_ = 0.4), which had low image quality and densely-packed cells, all others had one pretrained Cellpose model outperforming watershed and AP_0.5_ ≥ 0.6. Mouse organoids and skull nuclei matched EmbedSeg3D. For mouse organoids, pretrained cyto2 with u-Segment3D (AP_0.5_ =0.93) nearly matched the ideal 2D segmentation (AP0.5=1.0).

### Specialist 2D models with u-Segment3D are competitive to specialist native 3D models

Our experiments with pretrained 2D models suggests u-Segment3D can translate 2D orthoview segmentations with performance rivalling natively trained 3D models. We next formally assessed this to investigate pros and cons of 2D-to-3D segmentation across all 8 datasets (minus DeepVesselNet) amenable to be trained with 3D instance cell segmentation models. For each dataset we constructed training, validation, and test splits and performed (i) distance transform watershed (dtWS) as an unsupervised classical baseline, and (ii) trained specialist 3D models and (iii) counterpart specialist 2D models from 2D slices pooled across orthoviews (Fig. 4a, Methods). We evaluated EmbedSeg3D and 2D^15^ as representative of the state-of-the-art instance embedding approach; StarDist3D^47^ and 2D^2^ as representative of the shape-prior approach; PlantSeg3D^7^ and 2D as the original and state-of-the-art method for segmenting Ovules and LRP datasets; and trained the latest state-of-the-art 2D Cellpose3^62^ models with masks generated by u-Segment3D (when suffixed with (u)). 2D segmentations were then translated to 3D with indirect u-Segment3D (Methods). In Ovules, u-Segment3D consensus AP and F1 curves (lighter-colored dashed lines) match native 3D models (darker-colored solid lines), (Fig. 4b). StarDist2D (AP_0.5_ = 0.7, *F*10.5 = 0.85) is even better than StarDist3D (AP_0.5_ = 0.6, *F*1_0.5_ = 0.75). This highlights 2D-to-3D as a superior approach for complex 3D shapes, whose 2D cross-sectional shape may be overall simpler, i.e. more convex and less elongated. All 2D models outperformed 3D counterparts for LRP by as much as 0.2 in AP_0.5_, *F*1_0.5_ for StarDist (Fig. 4c). Most dramatically, for mouse skull nuclei (Extended Data Fig. 14d), u-Segment3D achieved AP_0.5_ = 0.8, *F*1_0.5_ = 0.9 with StarDist2D, whereas StarDist3D failed (AP_0.5_ = 0.05, *F*1_0.5_ = 0.1). EmbedSeg3D was the best method only for Arabidopsis and nuclei datasets (Extended Data Fig. 14d). The advantage of native 3D models appears primarily in capturing thin and small 3D cells (relative to the image size), that are not well captured by 2D orthoslices. Plotting 2D AP_0.5_ vs 3D AP_0.5_ of its u-Segment3D consensus across datasets and models, u-Segment3D faithfully translates 2D performance (Pearson’s *R* = 0.79) and in most cases, 3D AP_0.5_ is higher (Fig. 4d). The 2D AP, F1 curves verify that u-Segment3D parses Cellpose outputs with additional flexibility without affecting performance (Extended Data Fig. 15). Across datasets and models, 3D AP_0.5_ of native 3D segmentations correlates with the corresponding u-Segment3D consensus segmentation (Pearson’s *R* = 0.57). Individually, except for Platynereis nuclei, there is at least one 2D model where u- Segment3D AP_0.5_ matches or exceeds the native 3D segmentation (Fig. 4e). For Platynereis nuclei, the discrepancy is not due to failure of u-Segment3D to translate 2D performance but insufficient capture of smaller cells in 2D slices. Training an EmbedSeg3D model guided by EmbedSeg2D u-Segment3D consensus segmentation (Methods) restored smaller cells and boosted AP_0.5_ to 0.85, much closer to the native EmbedSeg3D (AP_0.5_ = 0.95), (Fig. 4f). Lastly, u-Segment3D can integrate less than three orthoviews in any combination. Depending on the cell packing and shape symmetry, this reduces computation with results comparable to using all three orthoviews, as we demonstrate for LRP (Fig.4g, Suppl. Movie 6).

**Figure 4.**
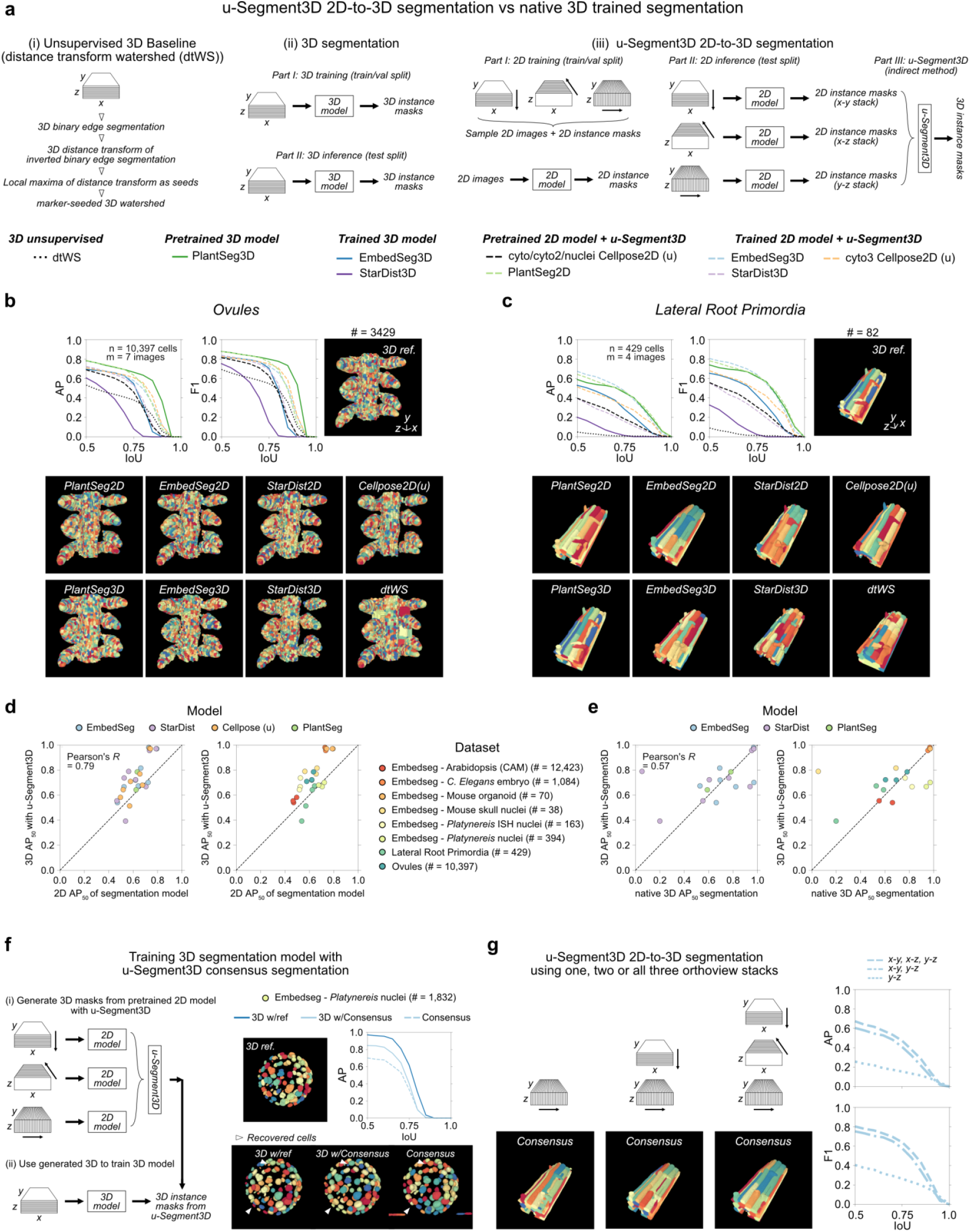
Consensus 3D segmentation from 2D stacks is competitive with native 3D segmentation and superior for densely packed cells and cells with complex morphologies. **a)** Schematic of the steps to construct i) classical unsupervised distance transform watershed (dtWS) 3D segmentation baseline (left), and the training and application for ii) native 3D segmentation (middle) and iii) consensus 3D segmentation from 2D models with u-Segment3D (right). AP and F1 curves on test split and 3D render of reference cell masks of an example image (top), and corresponding output with each 3D segmentation method (bottom) for **b)** Ovules and **c)** Lateral Root Primordia datasets. Each trained model is colored uniquely with solid line, darker hue for native 3D segmentation and dashed line, lighter hue for consensus segmentation using their trained 2D counterpart model. **d)** AP_50_ of trained 2D model in 2D segmentation (x-axis) plotted against matching AP_50_ of 3D performance after u-Segment3D. Each point is uniquely colored by 2D model (left) and dataset (right). In b)-d) Cellpose 2D segmentations were generated from predicted outputs using u- Segment3D’s method, indicated by the (u) annotation. **e)** AP_50_ of training native 3D model in 2D segmentation (x-axis) plotted against AP_50_ of training its 2D model counterpart and consensus segmentation with u- Segment3D. Each point is uniquely colored by model (left) and dataset (right). In d) and e) dashed black line represents identical performance. **f)** Schematic of training an EmbedSeg3D model using u-Segment3D generated consensus segmentation of EmbedSeg2D segmented stacks (left). Render of the reference 3D ground-truth masks and average precision curve of native 3D training with reference 3D masks (w/ref), with consensus segmentation derived from 2D (w/Consensus), and the consensus generated by u-Segment3D from 2D (Consensus) (top row, right). 3D renders of the segmentation for the same image as the reference, with each method (bottom row, right). **g)** 3D render of the consensus segmentation from using only one (y-z), two (x-y and y-z) and all three (x-y, x-z and y-z) EmbedSeg2D segmented stacks of an example image (left). Corresponding average precision and F1 curve evaluated across all test images using one, two and all three orthoviews.

### For anisotropic 3D data u-Segment3D can construct consensus 3D segmentation from 2D slice-by-slice instance segmentations from a single orthoview

Due to the microscope or culture conditions, 3D images cannot always be acquired isotropically or be interpolated to be near-isotropic with an image quality similar in x-y, x-z, and y-z orthoviews. In these cases, application of 2D segmentation models, trained solely on the equivalent of in-focus ‘x-y’ slices, to x-z and y-z views may yield worse 3D segmentations. As demonstrated above, to accommodate this scenario, u- Segment3D can use only x-y slices. u-Segment3D then operates similarly to slice-by-slice stitching method but still has the advantage of adjusting parameters such as smoothing to interpolate missing segmentations (Suppl. Table 1). Looking top-to-bottom through an epidermal organoid culture^83^ (Methods), cells are flat and elongated in the suprabasal layers transitioning to cuboidal in the basal layer (Fig. 5a). Even when interpolated to isotropic voxels, suprabasal cells remain flat, and stretched in appearance (Fig. 5b). Instead of training models for each view, we applied pretrained Cellpose 2D to segment only x-y slices, using an optimal predicted diameter set per slice. u-Segment3D then translated the 2D segmentations into 3D. We compared the 3D segmentation from 2D segmentations based on Cellpose predicted optimal diameters (Fig. 5c,d) and u-Segment3D contrast score diameters (Fig. 5e,f). Qualitatively, both looked similar. Without ground-truth, and ambiguity in manual labeling without a nuclear marker, we assessed the segmentation consistency between consecutive x-y slices, slice *i* and slice *i* + 1 with AP0.5. This revealed AP0.5 variation correlated with morphology, with a systematic drop in AP_0.5_ as cell morphology changed from squamous to cuboidal. Overall, the contrast score method appears more stable, with a higher mean AP_0.5_ = 0.59. We plotted the predicted mean cell diameter per slice (green line) with the measured cell diameter of the resultant segmentation (black line) for each method (Fig. 5g,h). Whilst Cellpose better predicts the absolute mean diameter per slice, its correlation across x-y slices was moderate (Pearson’s *R* = 0.47). In contrast, u- Segment3D’s contrast-score method exhibits strong correlation (Pearson’s *R* = 0.89). This consistency likely translated to the improved slice-to-slice AP_0.5_.

**Figure 5.**
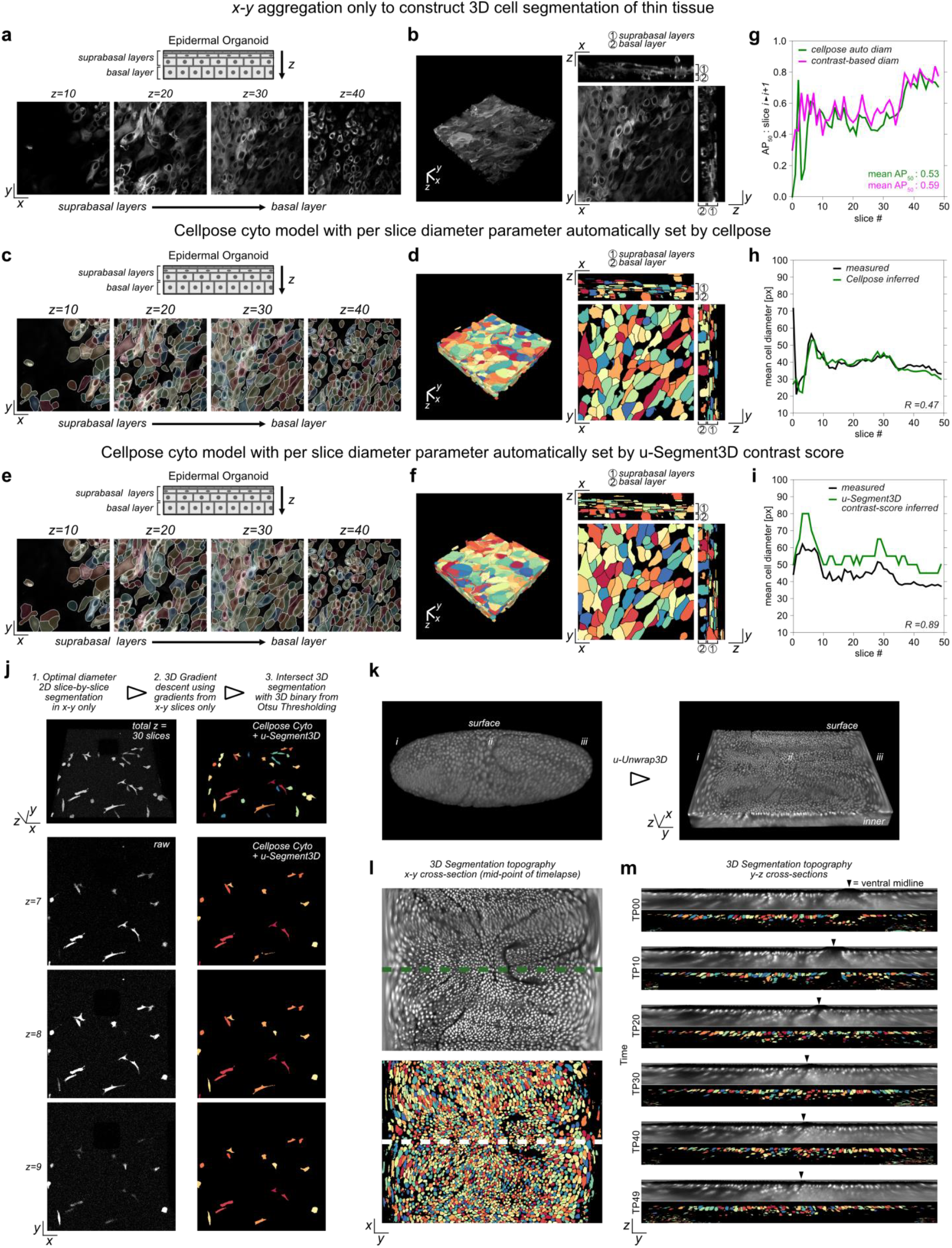
u-Segment3D consensus segmentation using only x-y 2D stacks. **a)** Four equi-sampled x-y image slices from top to bottom, capturing the suprabasal to basal layer transition within an epidermal organoid culture. **b)** 3D rendering of culture with axial interpolation to isotropic voxel resolution (left) and corresponding mid-section orthoslices (right). **c)** Cellpose 2D cell segmentations using the ‘cyto’ model and diameter automatically determined per-slice by Cellpose. Cells are individually colored and overlaid onto the four x-y image slices in a). White boundaries delineate individual cell boundaries within a slice. **d)** 3D rendering of the u-Segment3D consensus segmentation of the x-y 2D segmentation stacks in c) (left) and corresponding x-y slice at z = 20, mid-section x-z, y-z orthoslices (right). **e)** Cellpose 2D cell segmentations using the ‘cyto’ model and diameter automatically determined per-slice by u-Segment3D contrast score. Cells are individually colored and overlaid onto the four x-y image slices in a). White boundaries delineate individual cell boundaries within a slice. **f)** 3D rendering of the u-Segment3D consensus segmentation of the x-y 2D segmentation stacks in e) (left) and corresponding x-y slice at z = 20, mid-section x-z, y-z orthoslices (right). **g)** 2D cell segmentation consistency measured by AP_50_ between consecutive z-slices as a function of z-slice id for per-slice Cellpose model diameter auto-determined by Cellpose (green line) or u-Segment3D contrast score (magenta line). **h)** Mean cell diameter inferred by Cellpose (green line) and the mean cell diameter computed from the 2D cell segmentation (black line) for each x-y slice. **i)** Mean cell diameter inferred by peak position in the u-Segment3D contrast score (green line) and the mean cell diameter computed from the 2D cell segmentation (black line) for each x-y slice. *R* denotes the Pearson’s *R* in panels g)-i). **j)** Time-point by time-point segmentation of MDA231 human breast carcinoma cells from the 3D Cell Tracking Challenge using u-Segment3D to aggregate x-y slice segmentations based on on Cellpose 2D ‘cyto’ models with optimal diameter selection by contrast score. 3D rendering of raw image volume and segmented cells (top) and in consecutive 2D x-y slices (bottom). **k)** Timepoint by timepoint segmentation of cells embedded in a surface proximal tissue section of a developing drosophila embryo. The surface of the embryo body was segmented and the surface proximal tissue section projected into a 3D thin volume using the u-Unwrap3D^72^ framework. **l)** Consensus segmentation of individual cells by u-Segment3D in the unwrapped tissue section at timepoint (TP) 25 (top) using Cellpose 2D ‘cyto’ models with the diameter optimized via u-Segment3D’s contrast score (bottom). **m)** Snapshots of the mid y-z cross-section (position indicated by green dashed line in l) of the raw (top) and segmented (bottom) unwrapped tissue volumes at 6 timepoints. Black arrowheads indicate the position of the ventral midline towards which cells converge from the two sides.

A second example is a video of MDA231 human breast carcinoma cells embedded in a collagen matrix from the single cell tracking challenge^84^ (Fig. 5j). These cells have small area, thin, protrusive morphologies and were imaged with a noisy background. The 3D image has only 30 z slices, each cell spanning <5 slices. Again, applying pretrained Cellpose with contrast-score diameter determination on x-y slices only, we successfully generated consistent 3D cell segmentations. Visual inspection confirmed the same cell was consistently segmented across slices. Applying our strategy to every timepoint, we also observed consistent segmentation of cells across time (Suppl. Movie 7).

Finally, we also segmented cells on the surface of a developing drosophila embryo, a second data set from the single-cell tracking challenge^84^. Due to the curved embryo surface, cell dynamics are better visualized in cartographic surface projections^85^. We first segmented the embryo surface u-Segment3D (Methods) and applied the u-Unwrap3D^72^ pipeline to define a cartographic projection of the surface proximal subvolume (Fig. 5k). We then applied u-Segment3D to this computationally flattened tissue section to extract individual 3D cells. Again, due to the relatively shallow depth of the section we used x-y slices only. Despite the distortions introduced by the tissue projection, u-Segment3D still produced consistent 3D cell segmentations (Fig. 5l). This enabled us to visualize the migration of cells toward the ventral midline (black arrows) from the side in relation to cells underneath the embryo surface (Fig. 5m, Suppl. Movie 8).

### Refinement of 3D segmentations and recovery of subcellular structures not captured by pretrained models

In 3D, cells exhibit a rich spectrum of protrusive, subcellular surface morphologies. Biologically, these protrusions are classified into recurring morphological motifs^73^ such as blebs, lamellipodia, filopodia, and villi. Existing 2D cell training datasets represent these fine-grained structures incompletely, compounded by the spectral bias of neural networks^86^, i.e. low-frequency modes are learned faster and more robustly than high-frequency modes. Rectifying the bias requires revising the model architecture and additional training on fine-grained higher quality labels^87^. To circumvent this formidable and often impossible task, u-Segment3D proposes a two-stage solution to be applied in postprocessing after filtering out implausible cells by size and gradient consistency (Fig. 6a, i-iii). The first stage (Fig. 6a, iv) is label diffusion, a semi-supervised learning^88^ technique to improve adherence to the cell boundaries within a guide image, whilst enforcing spatial connectivity. Each cell in the input segmentation defines unique ‘sources’ that simultaneously diffuse to neighbor voxels for *T* iterations based on an affinity graph combining the local intensity differences in the guide image with spatial proximity (Methods). The final segmentation is generated by assigning each voxel to the source with highest contribution (Methods). We control the extent of diffusion using a ‘clamping’ factor such that if ‘clamped’, diffusion only modifies voxels initially assigned as background. We observe improved boundaries for *T*<50 iterations. The guide image can be the intensity-normalized raw image or any image enhancing the desired features to be segmented.

**Figure 6.**
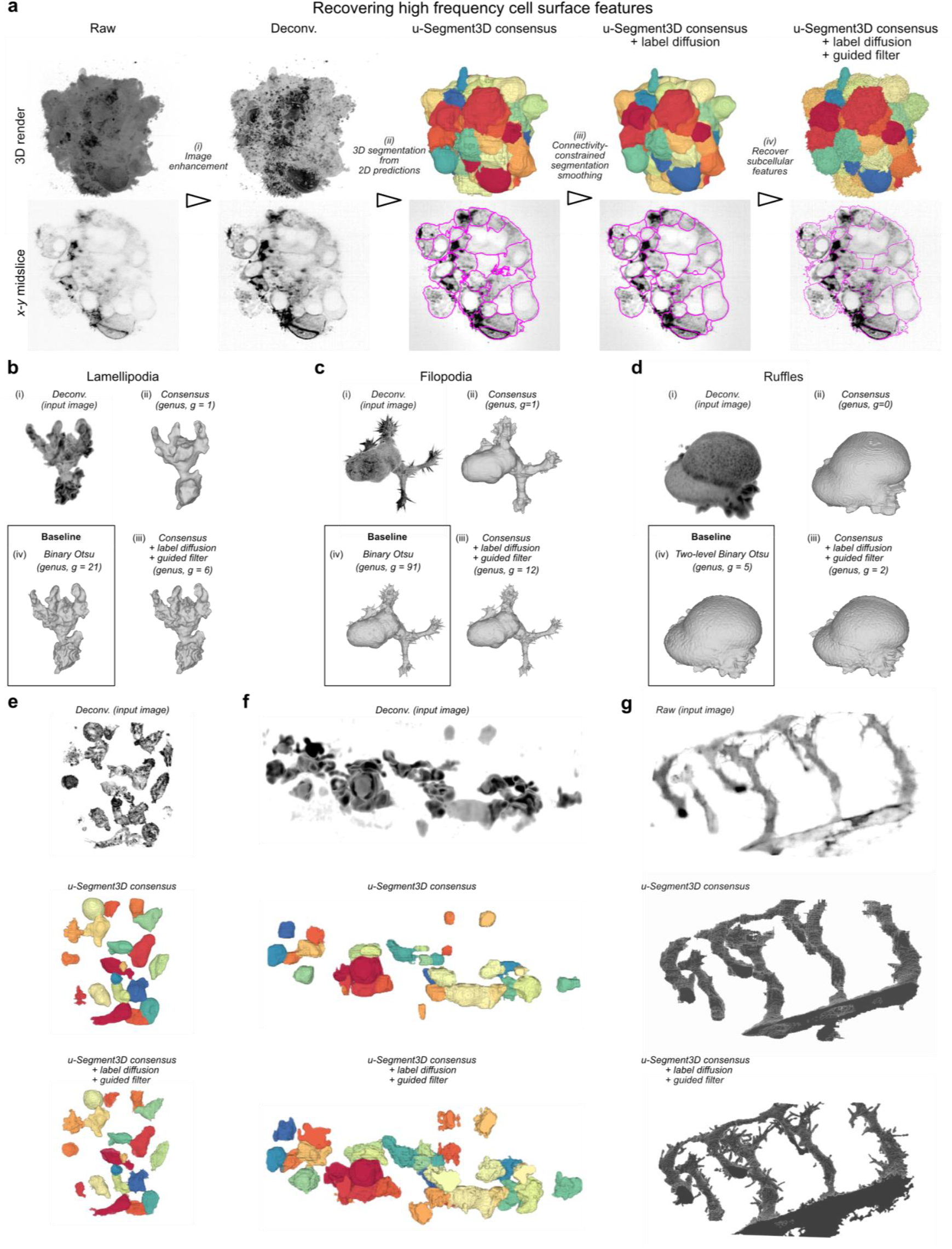
u-Segment3D postprocessing recovers missing high-frequency, high-curvature subcellular structures. **a)** Workflow with postprocessing to segment individual cells and recover subcellular features of each cell. 3D rendering (top) and x-y midslice (bottom) of the output at each step. **b)** Binary segmentation and recovery of lamellipodial features on a dendritic cell using u-Segment3d postprocessing. 3D rendering of the (i) deconvolved input, (ii) initial 3D consensus segmentation integrating Cellpose 2D cell probability maps (after step ii of a)), (iii) final postprocessed 3D segmentation (after step iv of a)) and comparison with the segmentation from binary Otsu thresholding on the 3D image intensity. *g* = genus of extracted surface mesh. **c), d)** Binary segmentation and recovery of filopodial and ruffle features on a HBEC and COR-L23 cell using u-Segment3d postprocessing as in b). **e)** Segmentation of T-cells with u-Segment3D postprocessing. 3D rendering of deconvolved image volume (top), initial 3D consensus segmentation integrating Cellpose 2D outputs (middle) and final 3D segmentation with recovered subcellular protrusions (bottom). **f)** Segmentation of zebrafish macrophages using Cellpose 2D outputs with u-Segment3D postprocessing. 3D rendering of deconvolved image volume (top), initial 3D segmentation from aggregated Cellpose 2D outputs (middle) and final 3D segmentation with recovered subcellular protrusions (bottom). **g)** Binary 3D segmentation of developing zebrafish vasculature using Cellpose 2D outputs with u-Segment3D postprocessing. 3D rendering of raw image volume (top), initial 3D segmentation from aggregated Cellpose 2D cell probability maps (middle) and final 3D segmentation with recovered sprouting vessels (bottom).

The second stage uses a linear guided filter^89^ to transfer detailed features from a guide image to the segmentation, constrained to the local neighborhood around the cell border (Methods). Conceptually, this filter is analogous to an interpolation between the binary cell mask and the intensities in the corresponding spatial region of the guide image. The neighborhood size may be fixed for all cells or set proportionate to cell diameter. For guided filtering, we find a good guide image to be *I*_*guide*_ = *α*. *I*_*norm*_ + (1 − *α*)*I*_*ridge*_, i.e. a weighted sum of the normalized input image, *I*_*norm*_ and its ridge filter-enhanced counterpart, *I*_*ridge*_, which amplifies subcellular protrusions (Suppl. Table 1).

Applying this workflow, we recovered the majority of missing surface protrusions for cells tightly packed as an aggregate whilst simultaneously retaining the benefits of the shape prior from Cellpose (Fig. 6a, Suppl. Movie 9). We expected that the workflow would allow us to segment cells imaged with high-resolution lightsheet microscopy even when membrane staining is inhomogeneous or sparse, situations that challenge thresholding-based techniques^73^. We tested this on single cells with different morphological motifs. As these image volumes contain only one cell, we applied a threshold directly to the 3D reconstructed cell probability (Fig. 6b-d i). The result captures well the global morphology but cell protrusions only approximately (ii). After all postprocessing steps, all protrusions are recovered (iii), with comparable fidelity to binary thresholding (iv). However, this segmentation is better suited for surface analysis, as measured by a lower genus, *g*, of the extracted surface mesh. We can further recover protrusive features on touching cells in a field-of-view as shown for T-cells (Fig. 6e) and zebrafish macrophages (Fig. 6f). Lastly, we tested the segmentation of zebrafish vasculature undergoing angiogenesis (Fig. 6g). The combination of using pretrained Cellpose 2D as shape prior and guided filtering recovered complete, thin sprouting vessels, despite the noisy background and inhomogeneous staining (Suppl. Table 5, Suppl. Movie 10)).

### Consensus 3D cell segmentations of tissue by parallel computing

Recent advances in high-resolution thick-section tissue imaging generate image volumes easily containing 10,000’s of cells^23^. The time for gradient descent increases with iteration number and the number of foreground pixels (related to image size). To allow segmentations to be computed in a reasonable time, we implemented a multiprocessing variant of the 2D-to-3D segmentation pipeline that takes advantage of the wide availability of CPU-based cluster computing (Methods). Fig. 7a illustrates the key steps: (i) Running pretrained 2D models by fast GPU inference^5,62^ on stacks of orthogonal views of the full volume; (ii, iii) applying gradient descent in parallel in local spatially-overlapped subvolumes whilst generating globally- indexed foreground coordinates. Spatial overlap ensures border cells across subvolumes retain the same global attractor, avoiding additional stitching in postprocessing (Methods); and (iv) using an efficient parallelized connected component analysis^90^ to generate the full image 3D instance segmentations from advected coordinates. Any segmentation postprocessing is subsequently applied in parallel to individual cells. The segmentation of a metastatic melanoma CyCIF multiplexed tissue sample using fused nuclear and membrane signal composites (Methods), imaged with an equivalent isotropic voxel size of 280 nm resolution and dimension 194 × 5440 × 4792 pixels took ≈ 2h for preprocessing and running Cellpose slice-by-slice in x-y, x-z, y-z; ≈ 2h to generate the initial 3D segmentation from 250 gradient descent iterations and using subvolumes of 128 × 256 × 256 pixels with 25% spatial overlap; ≈ 1 h for size filtering and gradient consistency checking; and ≈ 2h for label diffusion refinement. Thus, with a total of 7h processing time we extracted the volumes of 43,779 cells on a CPU cluster with 32 physical cores, 72 threads, 1.5TB RAM and a single A100 GPU (40GB) for initial 2D model inference. Notably, run on the full volume the gradient descent alone would be > 20x slower. Moreover, the obtained segmentations are free of stitching artifacts (Suppl. Movie 11). Inspecting the zoom-ins of the mid-slices from each of the three orthoviews, our segmentation agrees well with the fused cell nuclei and membrane markers (Fig. 7b, Suppl. Movie 11). Functionally, these segmentations enabled us to improve the accuracy of 3D cell phenotyping and to show, for example, how mature and precursor T cells in metastatic melanoma engage in an unexpectedly diverse array of juxtracrine and membrane-membrane interactions^23^.

**Figure 7.**
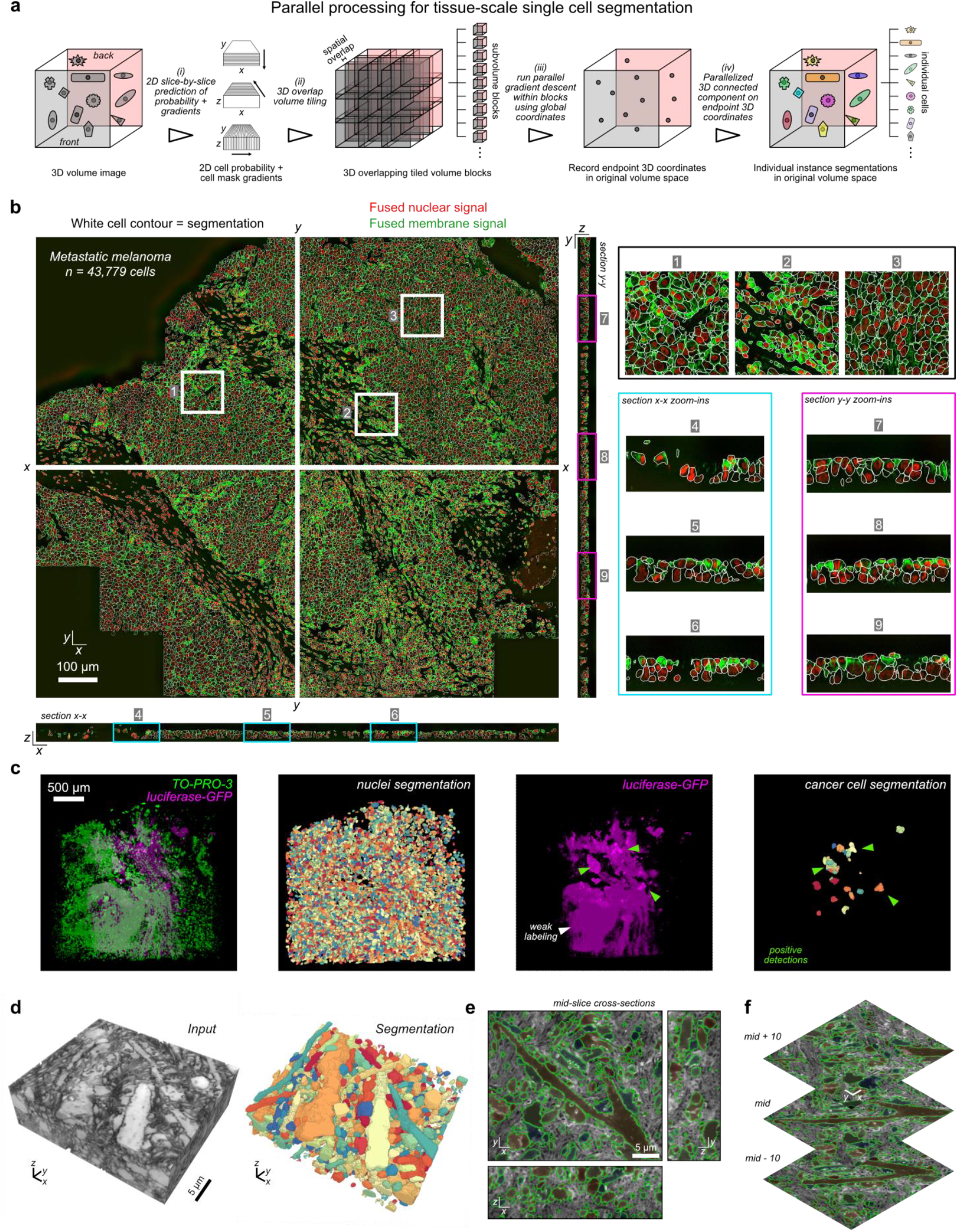
u-Segment3D implements parallel computing for tissue-scale segmentation. **a)** Schematic of the parallelized gradient descent in overlapped subvolume tiles used by u-Segment3D to facilitate consensus single cell 3D segmentation in large tissue volumes. **b)** x-y, x-z, y-z midslice cross-sections of the fused nuclear (red) and membrane (green) signal channels from multiple biomarkers (see Methods) for a CyCIF multiplexed patient biopsy of metastatic melanoma with white boundaries to delineate the individual cells of a u-Segment3D consensus segmentation using the Cellpose cyto model in each view (left). Zoom-ins of 3 subregions in x-y (black box, regions 1-3), x-z (cyan box, regions 4-6), y-z (magenta box, 7-9) cross-sections (right). **c)** u-Segment3D consensus segmentation of nuclei channel (TO-PRO-3, green) using the Cellpose nuclei model and cancer cells (luciferase-GFP, magenta) using the Cellpose cyto2 model in a cleared lung tissue of a mouse xenografted with YUMM 1.7 melanoma cells (Methods). Left-to-right: merged input volume image, individual segmented nuclei from nuclei channel, cancer luciferase-GFP channel only image showing weak, non-specific staining (white arrow) compared to the specific positive label of bona fide metastatic colonies (green arrow), and final u-Segment3D representation of metastatic cells post-filtered by mean cell luciferase-GFP intensity. **d)** 3D rendering of the input coCATs volume (left) and u-Segment3D consensus segmentation of salient tissue architecture using Cellpose cyto model (with model inverted) (right). **e)** Mid-slice cross-sections in x-y, x-z, y-z with individual segmentation boundaries in green and overlaid to the input image. **f)** Mid ± 10 z-slice x-y cross-section with individual segmentation boundaries in green overlaid on the input image.

Axially-swept lightsheet microscopy^101^ can image cleared tissue volumes at subcellular resolution over sections as thick as 2mm. This enabled us to visualize single cells within micrometastases in lung tissue. We found u-Segment3D could readily extract all nuclei in the invaded lung tissue stained by TO-PRO-3 and even segment luciferase-GFP cancer cells of the micrometastatic colonies, despite the weak GFP fluorescence (Fig. 7c, Suppl. Movie 12). To generate this cancer segmentation both channels were conjointly input to Cellpose. With the cancer channel alone, individual cells could not be resolved due to insufficient stain contrast between cell boundaries. The nuclei channel helped provide complementary information to localize individual cancer cells. Even so, 2D slice-by-slice segmentations are noisy and consequently u-Segment3D consensus 3D segmentation contained many extraneous, spurious segmented cells. However, because u- Segment3D implements robust methods for each algorithmic step, cells were not catastrophically fragmented, and invalid cells could be readily filtered out by mean luciferase-GFP intensity to identify only the bona fide cancer cells (Extended Data Fig. 16b).

Finally, we tested u-Segment3D on an image stack of brain tissue acquired by CATS^74^, a recent technique that labels the extracellular space of tissue. Tissue structure based on this label is difficult to discern (Fig. 7d). We hoped Cellpose and u-Segment3D could provide an exploratory tool that ‘scans’ the volume to generate consensus representations of larger pockets in extracellular space. These spaces are heterogeneous, different in size, and morphotype, which challenges Cellpose’s implicit approach to produce homogeneously scaled segmentations. Applying the default u-Segment3D pipeline led to fragmentation of the thick, dominant, branching dendrites (data not shown). However, by adjusting parameters of the u- Segment3D 2D-to-3D segmentation process we can overcome limitations of the 2D model and successfully preserve the multiscale tissue architecture, both in 3D and in 2D cross-sections (Fig. 7d-f, Suppl. Movie 13, Suppl. Table 1,5). Specifically, we applied increased Gaussian smoothing (*σ* = 2) to the reconstructed 3D gradients, utilizing smoothing across cells to correct noisy vectors that otherwise would split single cells. We ran suppressed gradient descent with a larger decay (τ = 0.25) to reduce the splitting of branched structures. Finally, we set *σ* =1.2 to compute the point density after gradient descent used to identify unique cells with binary thresholding and connected component analysis. The slightly larger *σ*(= 1 in rest of manuscript) merges clusters separated by a small distance that are part of the same 3D cell, without affecting those clusters that genuinely represent single cells and located further apart.

## Discussion

By considering shape reconstruction, we present a formalism to generate optimal, consensus 3D segmentations from stacks of 2D segmented slices. Our formalism unifies existing proposals of 2D-to-3D segmentation and shows near perfect 3D segmentations are achievable across single cells, dense tissue contexts, and morphotypes. We refer to the proposed consensus segmentation as ‘universal’ to describe how it (i) is applicable to diverse morphologies, with minimal adjustment of parameters (e.g. by specifying a gradient decay in the gradient descent to account for significant branching in shapes), (ii) exhibits no bias to any particular orthoview, and (iii) encapsulates and generalizes existing stitching and gradient tracing approaches for 2D-to-3D segmentation.

Conceptually our work reformulates the ad hoc procedure of stitching discrete label segmentations into a continuous domain problem with well-behaved mathematical properties. This not only enabled true integration of complementary information from orthoviews without bias (Fig. 3) but also empowers users to perform well-rationalized fine-tuning of the pipeline rather than empirical trial-and-error (Suppl. Table 1), which is prohibitive with hard-to-visualize 3D datasets.

These theoretical insights led us to develop a general toolbox, u-Segment3D, to robustly implement 2D-to- 3D segmentation universally for any 2D method that generates pixel-wise instance segmentation masks. Other representations should be first converted to labeled images to use u-Segment3D. For example, using connected component analysis to label individual contiguous regions in 2D binary masks, and rasterization to convert polygonal contour segmentations to pixel-wise masks. Through extensive validation on public datasets, we showed with proper parameter setting, u-Segment3D faithfully translates 2D model performance without further data training to 3D, with results competitive to equivalent native 3D segmentation models (Fig. 4). The better the 2D model, the better the 3D segmentation. u-Segment3D opens up the development of creative 3D solutions beyond the scope of this paper including training of individual 2D models for each orthoview, and extension to more than three views. Moreover u-Segment3D provides fine-tuning and postprocessing methods for further improving 3D segmentations. We also implemented multiprocessing to enable scalable 2D-to-3D segmentation on CPU clusters. Further speed improvements could be made such as implementing a multi-scale scheme to run u-Segment3D, which we leave for future work.

With the successes of foundation models such as ChatGPT in natural language processing and Segment Anything for object segmentation in image analysis, there is a prevalent notion that everything should be learnt from data, that more data is better and models should be ‘turnkey’, working directly out-of-the-box or if not, be ‘fine-tuned’ on more data. In the quest for generality, we must not neglect the value of grounded formalism and robust design. Our analyses provide multiple cautionary tales. First, optimally parsing the outputs of neural network models is just as important as training. By identifying and rectifying the spatial proximity clustering of Cellpose, we significantly reduced over-segmentation and boosted performance on noisy and out-of-distribution datasets. Second, considering extremal morphotypes and the simpler 1D-to-2D segmentation problem showed the critical importance of suppressed gradient descent to enable 2D-to-3D segmentation to be applied to branched and network structures. Third, running Cellpose with optimal diameters in different views is necessary to capture general 3D shape, irrespective of voxel anisotropy. Because Cellpose was trained using a fixed size diameter, we exploited Cellpose and its ability to learn a strong cell shape prior to ‘scan’ and infer all salient diameters of objects in the image. This enabled us to set optimal diameters in orthogonal views for 3D segmentation training-free. Lastly, by recognizing the spectral bias of neural networks and annotation bias, we developed simple label diffusion and guided filter postprocessing to recover intricate surface morphologies of 3D cells. This enabled us to extend any pretrained neural network to segment high-resolution single cells comparable to state-of-the-art classical methods but with 3D surfaces more suited for mesh-based applications.

Individually, each point is a minor detail. However, in practice, these implementation details truly make the difference. Together they compound, leading to inaccurate segmentations, and more importantly unpredictable performance for non-ideal inputs. This is why u-Segment3D carefully considered all practicalities of a consensus segmentation pipeline and implemented rationalized solutions to ensure guaranteed performance with respect to theory.

In sum, our experiments question the proposition value of directly training 3D segmentation models when a viable 2D segmentation model is available, and more broadly, the reliance on ‘big data’ data-driven approaches to solve problems without proper mathematical formulation. Using only pretrained Cellpose models equipped with automated parameter tuning, we demonstrate an unprecedented capacity to 3D segment cells from diverse microenvironments, from single cells through to entire tissues, acquired in-vitro, in-vivo and in-situ, from different modalities and with different resolutions. With widespread availability of diverse generalist and specialized 2D segmentation models, u-Segment3D paves a way towards accessible 3D segmentation for all, translating time-consuming annotation and training towards more impactful time spent on analyzing the acquired 3D datasets to provide biological insights.

## Author Contributions

Conception: AJ, FYZ (using pretrained 2D for 3D segmentation), FYZ (u-Segment3D); Investigation and Analysis: FYZ, CY, ZM. Data generation: CY (CyCIF tissue), SD (Zebrafish macrophages), ZM, MTI, JL, HMB, KMD, SJM (lung micrometases), BN (epidermal organoid), EJ (T cell co-culture), GMG, BJC (COR-L23 single cell with ruffles), ZM, JL (ctALSM microscope development), HMB (tissue clearing), AW, HMB, KD (Septin cleared tissue), RF (HBEC cell aggregate). Supervision: GD; Funding acquisition: GD, KMD, RF, PKS; Writing – Original Draft: FYZ, GD; Writing – Review and Editing: all authors.

## Acknowledgments

Funding for this work in the Danuser lab was provided by the grants R35 GM136428 (NIH) and U54CA268072 (NIH). Software dissemination by the Danuser lab is funded by RM1 GM145399. GMG is a HHMI Hanna H. Gray Fellow (GT16003). EJ was funded by the Wellcome Trust (grant #224040/Z/21/Z). Funding for the 3D data acquisition in the Sorger Lab was provided by the Ludwig Cancer Research, PCA (U2C-233262) and HTA (U2C-233280). The Fiolka lab is supported by the grants R35 GM133522 (NIGMS) and R01EB035538 (NIBIB). T-cell culture imaging was performed using the Oxford-Zeiss Centre of Excellence in Biomedical Imaging and the Kennedy Trust for Rheumatology Research (Grant #202117 and 202103, respectively). For epidermal organoid imaging, the authors would like to acknowledge the Quantitative Light Microscopy Core, a Shared Resource of the Harold C. Simmons Cancer Center, supported in part by an NCI Cancer Center Support Grant, 1P30 CA142543-01. This research was supported in part by the computational resources provided by the BioHPC computing facility located in the Lyda Hill Department of Bioinformatics, UT Southwestern Medical Center, TX. URL: https://portal.biohpc.swmed.edu. The research within this work complies with all relevant ethical regulations as reviewed and approved by the University of Texas Southwestern Medical Center. Zebrafish husbandry and experiments described here have been approved and conducted under the oversight of the Institutional Animal Care and Use Committee (IACUC) at UT Southwestern under protocol number 101805.

## Data Availability

All data used in this study except for the imaging of Septin cleared tissue (Fig. 1f), HBEC cell aggregate (Fig. 5a), and zebrafish vasculature (Fig. 5g) are publicly available. The public segmentation datasets are available from their sources as documented in Suppl. Table 1 and in the Dataset section of Methods. The other exemplar data used are provided as example data with the code, with the download link provided in the u- Segment3D GitHub readme. Any non-available data will be made available on request to the corresponding author.

## Code Availability

u-Segment3D is available both through the Python Package Index, PyPI and at https://github.com/DanuserLab/u-Segment3D. The GitHub repository includes installation instructions, our code to automate parameter tuning of Cellpose models, a download link to example data and code for a MATLAB based GUI for running u-Segment3D.

## Methods

### Datasets

#### Validation Datasets

10 independent public datasets with reference 3D segmentation labels and 1 dataset collected in-house with 3D segmentations generated with the aid of u-Segment3D were used to evaluate the ability of u-Segment3D to reconstruct 3D objects from ideal and predicted orthogonal x-y, x-z, y-z 2D slices (Fig. 2). 9 of the public datasets with both images and reference segmentations were used to assess the performance of u- Segment3D with pretrained Cellpose models (Fig. 3). Details of all datasets are given in Suppl. Table 1.

#### Demonstration Datasets

The following datasets were collected largely in-house and segmented using either the pretrained Cellpose cyto, cyto2 or nuclei 2D segmentation models, whichever was observed qualitatively to perform best after using u-Segment3D (if not stated otherwise). Parameter details are provided in Suppl. Table 5.

##### Cleared tissue Septin imaging (Fig. 1f)

2 million MV3 melanoma cells expressing SEPT6-GFP^21^ were subcutaneously injected into the flank of a NOD/SCID mouse, as previously described^91^. The xenograft tumor was grown until palpable (approximately 3 weeks), at which point the mouse was sacrificed and the tumor was excised along with nearby surrounding tissue within ≈5 mm. The tumor was cut into quadrants, cleared using the CUBIC method^92^, and melanoma cells within the invasive margin were imaged via ctASLM^93^).

##### 3D epidermal organoid culture (Fig. 5a-i)

###### Cell culture

Human keratinocyte Ker-CT cells (ATCC #CRL-4048) were a kind gift from Dr. Jerry Shay (UT Southwestern Medical Center). Ker-CT cells stably expressing mNeonGreen-tagged keratin 5 (K5) to label intermediate filaments was created as previously described^94^.

###### Epidermal organoid culture

We adapted an epidermal organoid culture model from existing protocols^94-96^. Polycarbonate filters with 0.4*μ*m pore size (0.47cm^2^ area, Nunc #140620) were placed in larger tissue culture dishes using sterile forceps. On day 0, a cell suspension of 5×10^5^ keratinocytes expressing K5-mNeonGreen in 400*μ*L of keratinocyte serum free media (K-SFM) was added to each filter well, and additional K-SFM was added to the culture dish to reach the level of the filter. On day 10, culture medium was aspirated from above the filter to place the cultures at an air-liquid interface. At the same time, medium in the culture dish was changed from K-SFM to differentiation medium^94^. On day 13, an additional 0.5 mM calcium chloride was added to the differentiation medium in the culture dish. Mature epidermal organoids were processed for imaging on day 20, after 10 days of differentiation at the air-liquid interface. Throughout the procedure, culture media were refreshed every two to three days.

###### Epidermal organoid imaging

Mature epidermal organoids were transferred to a clean dish, washed three times with PBS, then fixed in 4% paraformaldehyde (Electron Microscopy Sciences #15713) for 1 hour at room temperature. Filters with the organoids were cut out of the plastic housing using an 8 mm punch biopsy tool and inverted onto glass-bottom plates. Throughout imaging, PBS was added one drop at a time as needed to keep each organoid damp without flooding the dish. Organoids were imaged using a Zeiss LSM880 inverted laser scanning confocal microscope equipped with a tunable near-infrared laser for multiphoton excitation and a non-descanned detector optimized for deep tissue imaging. Images were acquired using an Achroplan 40×/0.8NA water-immersion objective resulting in an effective planar pixel size of 0.21 µm, and z-stack volumes with 1 µm step size.

##### Single cell tracking challenge datasets^84,97^ (Fig. 5j)

###### MDA231 human breast carcinoma cells (Fluo-C3DL-MDA231) (Fig. 5j)

Cells infected with a pMSCV vector including the GFP sequence, embedded in a collagen matrix were imaged with an Olympus FluoView F1000 microscope with Plan 20x/7 objective lens, sampling rate of 80 min and *xyz* voxel size 1.242 × 1.242 × 6.0 µm.

###### Drosophila Melanogaster embryo (Fluo-N3DL-DRO) (Fig. 5k-m)

Developing embryo imaged on a SIMView light-sheet microscope^98^ with a sampling rate of 30s, 16x/0.8 (water) objective lens and *xyz* voxel size 0.406 × 0.406 × 2.03 µm. We used ‘Cell01’ from the test dataset containing 50 timepoints. We used the pretrained Cellpose *cyto* model and u-Segment3D to segment the surface for each timepoint (Suppl. Table 4). Using the binary segmentation, we unwrapped a proximal surface depth using u-Unwrap3D^72^.

##### Human bronchial epithelial (HBEC) cell aggregate (Fig. 6a)

Transformed HBEC cells expressing eGFP-Kras^V12^ were cultured and imaged using meSPIM and published previously^*99*^.

##### Single dendritic cell with lamellipodia (Fig. 6b)

Conditionally immortalized hematopoietic precursors to dendritic cells expressing Lifeact-GFP were cultured, imaged and published previously^73^.

##### Single HBEC cell with filopodia (Fig. 6c)

HBEC immortalized with Cdk4 and hTERT expression and transformed with p53 knockdown, Kras^V12^ and cMyc expression cultured, imaged and published previously^73^.

##### Single COR-L23 cell with ruffles (Fig. 6d)

###### Culture

COR-L23 cells (Human Caucasian lung large cell carcinoma) were resuspended in 2mg/mL bovine collagen (Advanced BioMatrix 5005) and incubated for 48 hours in RPMI 1640 medium (Gibco 11875093) supplemented with 10% fetal bovine serum (PEAK SERUM PS-FB2) and 1% antibiotic-antimycotic (Gibco 14240062). Cells were incubated in a humidified incubator at 37°C and 5% carbon dioxide.

###### Imaging

Images were acquired with our home-built microscope system that generates equivalents to dithered lattice light-sheet through field synthesis^100^. Briefly, the system employs a 25X NA 1.1 water immersion objective (Nikon, CFI75 Apo, MRD77220) for detection, and a 28.6X NA 0.7 water immersion (Special Optics 54-10-7) for illumination. With a 500 mm tube lens, the voxel size of the raw data is 0.104 µm × 0.104 µm × 0.300 µm. The volumetric imaging was performed by scanning the sample along the detection axis. We used a Gaussian light-sheet and optimized the light-sheet properties so the confocal length was enough to cover the cell size without sacrificing too much the axial resolution^101^. Typically, the light-sheets are about 20 µm long and 1 µm thick. In each acquisition, we optimized the laser power and the exposure time to achieve fast acquisition without introducing too much photo-bleaching. Usually, the time interval for our volumetric acquisition is chosen to be either 5 or 10 s.

##### T-cell coculture (Fig. 6e)

Blasted human CD8^+^ T cells, were produced by activating naïve T cells isolated from PBMCs using anti- CD3/CD28 Dynabeads for 2 days, then rested for 5 days after removing the beads. Cells were frozen on day 5 of resting and thawed 48 hours before use. T cells were grown in complete RPMI-1640 (10% FCS, 1% Pen/Strep, 1% Glutamine, 1% HEPES) + 50 U/ml of IL-2. For migration-based imaging of multiple T cells, 8- well glass-bottom IBIDI chambers were coated with 1*μ*g/ml of hICAM-1-6xHis linker and hCXCL11 (Peprotech) for 1 h at room temperature, washed, then coated with 1% BSA. 0.5×10^6^ blasted human CD8^+^ T cells were labelled with CellMask DeepRed diluted to a 1x working solution in imaging buffer (i.e., colorless RPMI with 1% added Pen/Strep, 1% glutamine, 1% HEPES) for 30 minutes at 37^0^C. Cells and the glass slides were washed and resuspended in pre-warmed (37°C) imaging buffer. 0.1-0.2×10^6^ Cells were gently added to the coated glass slides and left to settle for 30-minutes before imaging. Cells were imaged using the Lattice Lightsheet microscope 7 (LLSM7) from Zeiss using the 641nm laser at 4% power with 4ms of exposure. A large field of view was used for imaging multiple cells at once, with a complete volume taken every second. Deconvolution was performed using the Zeiss software.

##### Zebrafish macrophages (Fig. 6f)

Zebrafish larvae with fluorescent macrophages, labeled with Tg(mpeg1:EGFP) were imaged as published previously^80^ (Suppl. Table 2).

##### Zebrafish vasculature (Fig. 6g)

Zebrafish (*Danio rerio*) embryos, larvae and adults were kept at 28.5°C and were handled according to established protocols^102,103^. All zebrafish experiments were performed at the larval stage and therefore the sex of the organism was not yet determined. To visualize the growing vasculature at around 34 h post fertilization (hpf), zebrafish larvae expressing the vascular marker *Tg(kdrl:Hsa*.*HRAS-mCherry)*^*104*^ in a casper background^105^ were used. To immobilize the zebrafish larvae for imaging, they were anesthetized with 200 mg/l Tricaine (Sigma Aldrich, E10521)^106^ and mounted in 0.1% low melting agarose (Sigma Aldrich, A9414) inside fluorinated ethylene propylene (FEP) tubes (Pro Liquid GmbH, Art: 2001048_E; inner diameter 0.8 mm; outer diameter 1.2 mm), coated with 3% methyl cellulose (Sigma Aldrich, M0387)^107^. The mounted zebrafish larvae were imaged on a custom multi-scale light-sheet microscope with axially-swept light-sheet microscopy^80^.

##### Multiplexed CyCIF tissue (Fig. 7b)

A primary melanoma sample from the archives of the Department of Pathology at Brigham and Women’s Hospital was selected. The protocol was adapted from Nirmal et al.^108^. Briefly, a fresh 35 µm thick FFPE tissue section was obtained from the block and de-paraffinized using a Leica Bond. The region in Fig. 6b was selected and annotated from a serial H&E section by board-certified pathologists as a vertical growth phase. The 35 µm thick section underwent 18 rounds of cyclical immunofluorescence (CyCIF)^109^ over a region spanning 1.4 mm by 1.4 mm and sampled at 140 nm laterally and 280 nm axially. Image acquisition was conducted on a Zeiss LSM980 Airyscan 2 with a 40x/1.3NA oil immersion lens yielding a 53-plex 3D dataset^23^. A custom MATLAB script was used to register subsequent cycles to the first cycle, which was stitched in ZEN 3.9 (Zeiss). The quality of image registration was assessed with Hoechst across multiple cycles in Imaris (Bitplane). For segmentation, multiple channel markers were combined to create fused nuclei and cytoplasmic channels. Hoechst and lamin B1 were combined for nuclei. MHC-II, CD31, and CD3E were combined as a cytoplasm marker to cover all cells including tumor, blood vessels, and T cells.

##### Cleared tissue lung micrometastases (Fig. 7c)

###### Cancer growth

Lung tissue containing a metastatic tumor was harvested from mice injected with YUMM 1.7 GFP-luciferase melanoma cells^110^ and grown as previously described^111^.

###### Lung tissue staining and clearing

Lung tissue was fixed in 4% paraformaldehyde at 4^0^C for less than 24 h and then washed three times with PBS with 0.02% sodium azide for 2 h per wash. The tissue was sliced into 2 mm thick sections. Tissue slices (∼2mm) were permeabilized and blocked in buffer (0.5% NP40, 10% DMSO, 0.5% Triton X-100, 5% donkey serum, 1X PBS) overnight at room temperature (RT). Tissues were incubated in anti-GFP (1:100) for 72 h at room temperature in a tube revolver rotator. After incubation, samples were washed with wash buffer (0.5% NP40, 10% DMSO, 1X PBS) three times for 2 h each and then left rotating in wash buffer overnight. Tissues were immersed in the AF488-conjugated secondary antibody solution (1:250) for 72 h at RT. Then, the secondary antibody was removed with wash buffer for at least two days changing the solution: the first day three times every 2 h and on the second day refreshed once. Finally, tissues were stained for nuclei with TO-PRO-3 647 (1:500) in PBS for 24 h at room temperature. Nuclear dye was washed out with wash buffer three times for 10 min each. Lung tissue was cleared using BABB clearing protocol (also known as Murray’s clear). Lungs were dehydrated in a methanol gradient (25%/50%/75%/100% for 20 min each). The final clearing was achieved with fresh Benzyl Alcohol and Benzyl Benzoate (BABB, 1:2). Prior to starting dehydration, 5 grams of aluminum oxide was added to 45 mL BABB and rotated at RT for at least an hour to remove peroxides. BABB was kept protected from light and air. The samples were quickly washed with BABB three times, then left standing in fresh BABB for 15 min. Sample BABB was refreshed and left overnight. The sample BABB was refreshed again shortly before imaging.

###### Lung tissue imaging

Lung tissue slices were imaged on a ctASLMv2^93^ microscope chamber controlled by navigate^112^. Nuclei were imaged using the TO-PRO-3 647 via illumination with a LuxX 642 nm, 140 mW at 100% laser power and a Semrock BLP01-647R-25 filter in the detection path. Cancer cells were imaged via illumination with a LuxX 488-150, 150 mW at 100% laser power and a Semrock FF01-515/30-32 bandpass filter in the detection path. Images were acquired with a Hamamatsu ORCA-Flash 4.0 v3 with 200 ms integration time in lightsheet readout mode.

##### coCATS labelled volume (Fig. 7d)

We used a coCATS^74^ imaging volume recorded with z-STED at near-isotropic resolution in neuropil of an organotypic hippocampal brain slice published in Michalska et al.^74^ (c.f. Fig. 3). This volume was downloaded already denoised with Noise2Void.

#### UMAP to map morphological diversity of different cell datasets

##### Morphological features

Eight features were extracted for each cell based on their 3D reference segmentations.

1. *Volume* - the total number of voxels occupied by the segmented volume.
2. *Convexity* - the ratio of total volume to total volume occupied by the convex hull. Convex hull was computed with Python Scipy, scipy.spatial.ConvexHull using the 3D coordinates of the binary volume.
3. *Major length* - Length of the longest axis of an ellipse fitted to the cell. Computed by Python Scikit- Image, skimage.measure.regionprops as the largest eigenvalue of the inertia matrix.
4. *Minor length* - Length of the shortest axis of an ellipse fitted to the cell. Computed by Python Scikit- Image, skimage.measure.regionprops as the smallest eigenvalue of the inertia matrix.
5. *1 – minor length/major length* - Measure of the extent of elongation with value 0-1. When spherical, minor length = major length and the measure is 0. When highly elongated, minor length << major length and the measure is 1.
6. *# skeleton segments* - Number of line segments composing the skeleton of the 3D binary volume.
7. *# skeleton nodes* - Number of branch point nodes, where a node is defined as at least three line segments meeting at a junction.
8. *Mean skeleton segment length* - Mean number of voxels in each segment of the 3D binary skeleton.

The 3D binary skeleton was computed using Python Scikit-Image, skimage.morphology.skeletonize. The decomposition of the skeleton into nodes and segments was performed using the Python sknw library (https://github.com/Image-Py/sknw). Non-dimensionless measurements such as volume were not converted to metric units as only the number of raw voxels is relevant for segmentation.

##### UMAP parameters

The 8 morphological features were power transformed to be more Gaussian-like using the Yeo-Johnson method^113^ (Python Scikit-learn, sklearn.preprocessing.power_transform). Then z-score normalization was applied to create normalized features. Uniform Manifold Approximation and Projection (UMAP) (using the Python umap-learn library) was used to project the 8 features after normalization to 2 dimensions for visualization (n_neighbors=15, random_state=0, spread=1, metric=‘Euclidean’). The median UMAP coordinate for each dataset was computed by taking the median of the 2D UMAP coordinates of individual cells comprising the respective dataset. The heatmap coloring of the UMAP uses the normalized feature value and the ‘coolwarm’ color scheme, clipping values to the range [-2,2].

#### 1D-to-2D segmentation and the effects of gradient smoothing

To compute 1D gradients, each spatially disconnected 1D region in a 1D slice is treated as a unique ‘cell’. For each ‘cell’, we computed the distance of each point to the cells’ centroid, computed the 1D gradient using central differences and unit length normalized the vectors. The 2D gradients are reconstructed by stacking the 1D gradients computed from x- and y- directions, and smoothed by applying an isotropic 2D Gaussian filter, width *σ* to x- and y- components. Gradient descent is iteratively performed to compute the trajectory, and the 2D segmentation was obtained by applying the image-based connected component analysis of u-Segment3D to the final advected 2D coordinates.

### u-Segment3D

u-Segment3D is a toolbox that aims to provide methods that require no further training to aggregate 2D slice-by-slice segmentations into consensus 3D segmentations. It is provided as a Python library (https://github.com/DanuserLab/u-Segment3D). The methods within can be broadly categorized into modules based on their purpose; module 1: image preprocessing; module 2: general 2D-to-3D aggregation using suppressed gradient descent with choice of different 3D distance transforms; and module 3: postprocessing to improve the concordance of segmentation to that of a guide image. Postprocessing helps achieve a tighter segmentation and recover missing local high-frequency surface protrusions.

### Module 1: Preprocessing

Described below are the image preprocessing functions included in u-Segment3D to address the primary problems of intensity normalization, image feature enhancement and uneven illumination that can greatly affect pretrained segmentation models, like Cellpose. Generally, the order of operation or the inclusion/exclusion of a step is dependent on the input data. We have found the basic workflow of i) rescaling to isotropic voxels and resizing for the desired segmentation scale, ii) uneven illumination correction, adaptive histogram equalization or gamma correction, iii) deconvolution, and iv) intensity normalization applied to the 3D raw image, to work well for Cellpose models. For Omnipose^6^ models we only use intensity normalization. Any other preprocessing led to worse performance. When both nuclei and cytoplasm channels are both available, we find Cellpose cell segmentation can be significantly better if both channels are used together as a RGB image (red channel – nuclei, green channel – cytoplasm) instead of nuclei or cytoplasm as a single grayscale image.

#### Rescaling to isotropic voxels and resizing to the desired segmentation scale

Pretrained segmentation models work best when input images contain object types and object sizes reflective of the original training dataset. If images are upscaled to be bigger, segmentation models may be biased towards segmenting physically smaller objects. Correspondingly if images are downscaled to be smaller, larger objects become enhanced and easier to segment as smaller objects become oversmoothed. Cellpose models are trained at a fixed diameter of 30 pixels and with isotropic ‘x-y’ images. We find empirically, the u-Segment3D tuning performs best for each orthoview if the input image volume is first rescaled to isotropic voxels and resized using linear interpolation so the desired feature to segment such as cell / vessel results in a peak around 30 pixels (c.f. Extended Data Fig. 10). The rescaling and resizing is implemented as one function using Python Scipy, scipy.ndimage.zoom function with a Python Dask tile-based accelerated variant for large volumes.

#### Contrast enhancing intensity normalization

Image intensities are normalized such that 0 is set to the *p*_*lower*_ percentile and 1 is the *p*_*upper*_ percentile of the image intensity. By default, *p*_*lower*_ = 2 and *p*_*upper*_ = 99.8. This contrast enhances the image by clipping out sporadic high intensities caused by saturated probe aggregates and zeroing small, but non-zero background intensities common to fluorescent microscopy.

#### Image deconvolution

For 2D fluorescent microscopy images or anisotropic 3D images, we use blind deconvolution with the unsupervised Wiener-Hunt approach^114^ (2D slice-by-slice for 3D) where the hyperparameters are automatically estimated using a Gibbs sampler (implemented using Python Scikit- image, skimage.restoration.unsupervised_wiener). The initial point-spread function is specified as a 15×15 pixel sum normalized Gaussian (*σ* = 1) squared kernel. For 3D lightsheet imaging we use Wiener-Hunt deconvolution, and our previously published experimental PSF^73^ as a ‘synthetic’ PSF.

#### Model-free uneven illumination correction

The raw image intensity of 2D or 3D images, 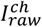 is corrected for uneven illumination ratiometrically, 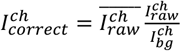 where 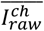 denotes the mean image intensity of the input image and 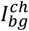 is an estimate of the uneven background illumination. 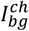 is estimated by downsampling the image by a factor of *ds*, isotropic Gaussian smoothing of *σ* then resizing back to the dimensions of the input image. For 2D images, the downsampling factor does not need to be used as Gaussian smoothing is fast and *σ* is specified as a fraction of the actual image dimension, typically 1/4 or 1/8 is a good starting point. For 3D images a default *σ* = 5 is used, with a *ds* = 8 or 16. If segmentation is worse at the higher *ds*, we decrease *ds* by factor of 2. If *ds* = 1, Gaussian smoothing is applied at the original image resolution. The resultant enhanced image should have even illumination with minimal artifactual enhancement of border background intensities. A more sophisticated background correction is the N4 bias correction available in SimpleITK, originally developed for MRI image and has been successfully applied to 3D cleared-tissue imaging^27^.

#### Adaptive histogram equalization (AHE)

Contrast limited AHE or CLAHE (Python Scikit-image, skimage.exposure.equalize_adapthist) can also be used as an alternative to our model-free uneven correction. The image is divided into non-overlapping tiles and the pixel intensity is histogram equalized within each tile. Whilst this obtains good results, we find that the method is computationally more memory-intensive and slower for large 3D volumes if the size of individual tiles is required to be small, thus increasing the overall number of tiles to be processed. However, there is less artifact for originally low-valued intensities compared to our faster ratiometric method.

#### Gamma correction

Transforms the input image, *I*_*in*_ pixelwise, raising the intensity to a power γ (float between 0-1) according so that the output image, 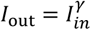 after scaling the image pixel intensity linearly to the range 0 to 1. Used to nonlinearly amplify low-intensity pixels to create a more uniform illumination for segmentation that is computationally inexpensive.

#### Ridge, vessel-like feature enhancement

Neurites, tubes, vessels, edges of cell surface protrusions all represent ridge-like structures that are both thin and long or exhibit high curvature and tortuous morphologies that are often only weakly stained and visualized from raw image intensities. Ridge image filters use the eigenvalues of the Hessian matrix of image intensities to enhance these ridge-like structures assuming the intensity changes perpendicular to but not along the structure. Many ridge filters have been developed. u- Segment3D uses the Meijering^*115*^ filter (Python Scikit-image, skimage.filters.meijering) which enhances ridge image features by pooling the maximum filter responses from multiple Gaussian *σ*. We observe empirically good performance for a diverse range of objects including vessels and cells, without requiring additional hyperparameters and hyperparameter tuning unlike Frangi filtering^116^.

#### Semi-automated diameter tuning for pretrained Cellpose models

The tuning process is illustrated in Extended Data Fig. 11a. Given a 2D image, the cellpose outputs: the non-normalized cell probability, *p* and predicted 2D gradients in x- (∇_*x*_Φ) and y- (∇_*y*_Φ) directions are computed. *p* is clipped to a range of [-88.72, 88.72] to avoid overflow for IEEE float32 and normalized to a value in the range [0,1], 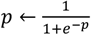. These ouputs define the pixelwise contrast score, *w*. {*σ*_𝒩_(∇_*x*_Φ) + *σ*_𝒩_(∇_*y*_Φ)}, where *w* is a pixelwise weight. We set *w* = *p* but observe no significant difference if *p* = 1 for cellpose models. *σ*(.) is the local standard deviation at each pixel, computed over the local pixel neighborhood of width *P* × *P* pixels. The mean score over all pixels, 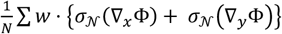 is computed over a range of equisampled diameters e.g. 15 to 120 at 2.5 increments. A centered moving window average (default window size 5) using symmetric padding at edges is then applied to smoothen the diameter vs score plot. Prominent peaks in this plot highlight potential segmentations at different size scales. The more sizes, the more peaks. Users may use this plot to inform the setting of a preferred diameter. For automatic operation, the diameter with highest contrast score is used. The neighborhood size acts like an attention mechanism (Extended Data Fig. 11c). The larger the neighborhood size, the more is the segmentation result corresponding to larger objects favored. If there is no larger salient segmentation, the optimal diameter will be the same as that found with a smaller neighborhood size.

#### Semi-automated cell probability thresholding for pretrained Cellpose models

We observe for out-of-distribution images and noisy input images, pretrained Cellpose 2D models can perform well using an appropriate threshold for cell probability combined with u-Segment3D’s gradient descent and spatial connected component analysis (c.f. Extended Data Fig. 4f vs Fig. 5j). The choice of threshold is particularly important. If the threshold is too high, there is no continuous path for the gradient descent, resulting in over-segmentation. It is therefore better to veer on the side of caution and use a lower threshold to get a more connected foreground binary. However, if the threshold is too low, the foreground binary will be larger than the region with predicted gradients. The additional foreground voxels have zero gradients and may segment as erroneous, extraneous cells. To automate the threshold, u-Segment3D applies multi-class Otsu thresholding to the normalized cell probability (*p* ∈ [0,1]) output of Cellpose, 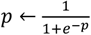. u-Segment3D further performs morphological closing to fill small holes. If only one object is known to be present, further operations such as extracting the largest connected component and binary infilling can be conducted. The default Otsu thresholding used in this paper is 2-class. If the segmentation partially captures the cells, we use 3-class Otsu and the lower of the two thresholds. Vice versa, if too much area is segmented, we use 3-class Otsu and the higher of the two thresholds. Optionally, we cast the threshold to the nearest decimal point, rounding down (*threshold* ← ⌊*threshold* ∗ 10⌋/10 where ⌊.⌋ is the floor operator).

### Module 2: Gradient descent and distance transforms to assemble 2D slice-by-slice segmentation stacks into a 3D consensus segmentation

Methods in this module are used to implement the core 2D-to-3D segmentation algorithm outlined in Fig. 1e. If 2D segmentations are not provided as a normalized cell probability (0-1) and 2D gradients in the manner of Cellpose^5^, then a 2D distance transform is specified to generate the necessary 2D gradients for consensus 3D segmentation.

### 2D Distance transforms

u-Segment3D categorizes the distance transforms according to whether the limit or attractor of propagating points using gradient descent over an infinite number of steps is implicitly or explicitly defined (Extended Data Fig.1). Explicitly defined transforms are further categorized by the type of attractor: a single fixed point source or a source that comprises a set of points.

u-Segment3D implements distance transforms, Φ that are solutions within the cell interior, of the Eikonal equation (*‖*∇Φ*‖*^2^ = 1, which gives the shortest geodesic solution) or Poisson’s equation (∇^2^Φ = −1, which gives a smooth harmonic solution). The Eikonal equation finds the shortest time of propagation for a point. Poisson’s equation can also be viewed as solving the shortest time of propagation but with the additional constraint of minimizing curvature, yielding smoother solutions.

### Implicit attractor distance transforms

With only the boundary condition Φ = 0, the Eikonal and Poisson equation conceptually propagates a wave inwards symmetrically from the cell boundaries. The limit solution is the definition of the medial axis skeleton, the locus of the centers of all inscribed maximal spheres of the object where these spheres touch the boundary at more than one point^75,117,118^.

#### Euclidean distance transform (EDT in text)

Solves the Eikonal equation using fast image morphological operations. u-Segment3D uses the memory and speed optimized implementation in the Python *edt* package released by the Seung Lab (https://github.com/seung-lab/euclidean-distance-transform-3d).

#### Poisson distance transform (Diffusion in text)

Solves the Poisson equation for each cell shape using LU decomposition (Python Scipy, scipy.sparse.linalg.spsolve). We parallelize the solving for all cells in an image using the Python Dask library.

### Explicit attractor distance transforms

The implicit attractor solves the equations everywhere in the cell interior. The explicit attractor variants modify the equations to have different source terms (right hand side of equation) in different parts of the cell interior. For the Eikonal equation, Φ = 0 at the cell boundary and outside, non-source points obey *‖*∇Φ*‖*^2^ = 1 whilst source points act as obstacles with vanishing speed, so that *‖*∇Φ*‖*^2^ = 0. For the Poisson equation, Φ = 0 at the cell boundary and outside, non-source points obey the Laplace equation, ∇^2^Φ = 0 whilst source points obey ∇^2^Φ = −1.

#### (i) Point sources

A single interior point is designated as a point source. u-Segment3D finds the interior point with Euclidean distance transform value greater than the percentile threshold (default: 10^th^ percentile) nearest the median coordinate of all points.

##### Eikonal equation solution (Geodesic centroid distance)

At the interior point, *‖*∇Φ*‖*^2^ = 0. The modified equations are solved using the Fast Marching Method (FMM)^77^, with the constraint enforced by the Python scikit-fmm library using masked arrays. Central first order differences are used to compute the unit normalized 2D gradient.

##### Poisson equation solution (Poisson or diffusion centroid distance)

Only at the interior point, ∇^2^Φ = −1. The modified equations are solved using LU decomposition as before. To apply power transformation with exponent *p* > 0, the minimum is first subtracted from Φ to ensure positivity, Φ^*p*^ ≔ (Φ − Φmin)^*p*^. Central first order differences are used to compute the respective unit normalized 2D gradient.

#### (ii) Point set sources

Any number of interior points are designated as point sources. u-Segment3D computes the 2D medial axis skeleton as the point set attractor. The binary skeleton of a binary cell image is computed by iteratively removing border pixels over multiple image passes^119^ (Python Scikit-image, skimage.morphology.skeletonize). This raw result often produces skeletons with extraneous branches that are too close to a neighboring cell. To improve the skeleton quality, the binary image is first Gaussian filtered with *σ* = 3 pixels (default), rebinarized by mean value thresholding and then skeletonized.

##### Eikonal equation solution (Geodesic centroid distance)

For all points part of the skeleton, *‖*∇Φ*‖*^2^ = 0. The modified equations are then solved using the Fast Marching Method (FMM)^77^ as above with central first order differences for computing the unit normalized 2D gradient. The gradients for all points part of the skeleton are set to zero to enforce the limiting behavior under gradient descent.

##### Poisson equation solution (Poisson or diffusion centroid distance)

For all points part of the skeleton, ∇^2^Φ = −1. The modified equations are solved using LU decomposition as above with central first order differences for computing the unit normalized 2D gradient. The gradients for all points part of the skeleton are set to zero to enforce the limiting behavior under gradient descent.

### Content-based averaging function, *F*

u-Segment fuses 3D volume images, *I*^*i*^ from *i* = 1, … , *N* multiple views using a content-based average function, *F*, with pixelwise weighting of the contribution of each view *i* given by the inverse local-variance, 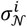, evaluated over an isotropic neighborhood, *N* of width *P* pixels

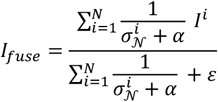

with *α* acting as a pseudo count. If *α* is small, 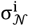 dominates. If *α* is large, 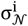 has little effect and all views are equally weighted. *ε* is a small value (10^−20^) to prevent infinity. For a neighborhood of width *P* = 1 pixels, *F* is equivalent to the simple mean used by Cellpose^5^ (Extended Data Fig. 2a). Compared to potentially more accurate approaches such as solving the multi-view reconstruction problem^79^, entropy-based averaging^120^ or using Gaussian filters^78^, the proposed *F* can be implemented more efficiently with uniform filters.

### Fusing normalized 2D cell probabilities (0-1) from orthoviews and binary thresholding

Stacked normalized 2D cell probabilities (0-1) are fused using the content-based averaging function, *F* above with neighborhood, *P* = 1 (default) pixels same as the fusion of 2D gradients below. For Cellpose models, the raw cell probability output, *p* are first clipped to the range [-88.72, 88.72] to prevent underflow/overflow in IEEE float32 and transformed, 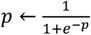 before fusing. For methods yielding only 2D segmentations, either i) fuse using the binary then apply appropriate Gaussian filtering to smooth, ii) use the intermediate cell probability image, which is always available for deep learning methods, or iii) generate a proxy cell probability image e.g. using a normalized Euclidean distance transform with values 0-1.

### Fusing 2D gradients from orthoviews

Stacked 2D gradients from x-y, x-z, y-z are pre-filtered with an isotropic Gaussian of *σ*_*pre*_ = 1. The fused 3D gradients combine three separate fusions: the fusing of x-component from x-y and x-z views, y-component from x-y and y-z views and z-component from x-z and y-z views. The constructed 3D gradients are post-filtered with *σ*_*post*_ = 1 (default) and unit length normalized. The greater *σ*_*post*_ is, the greater the regularization effect, reducing the number of attractors and preventing oversegmentation. This is helpful when using pretrained Cellpose models to segment cells that are larger and more branched than the majority of cells in an image. However, a large *σ*_*post*_ can also merge smaller cells. For fusion, we use *α* = 0.5 and in general *P* = 1 for *F* to maximize segmentation recall and use postprocessing to remove any erroneous segmentations. Larger *P* improves segmentation precision but may miss cells with lower contrast. These settings are generally not modified from the default. Preventing oversegmentation can be more controllably carried out by adjusting the temporal decay parameter in the gradient descent (see next, and Suppl. Table 1.).

### Gradient descent

Given the reconstructed 3D gradients, ∇Φ, gradient descent is applied to the set of all foreground image coordinates, {(*xn*, *yn*, *Zn*)}. The iterative update for gradient descent with momentum in 3D for iteration number, *t* = 0, … , *T*, with *T* = 250 defining the total number of iterations implemented by u-Segment3D, is

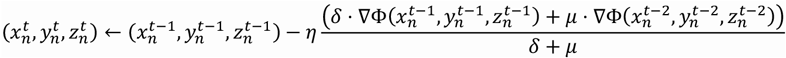

where *μ* is the momentum parameter governing the extent of influence of the previous gradient, ranging from 0-1 (default *μ* = 0.95), and *δ* > *μ* is the weighting of the current gradient and the step-size. *δ* is usually fixed with*δ* = 1 and only *μ* is adjusted. *μ* = 0 recovers the standard gradient descent iteration. Nearest neighbor interpolation is used for computational efficiency. As such, 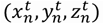 is always integer valued. *η* defines the step-size and varies as a function of the iteration number,

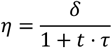

where τ ∈ ℝ^+^ is a floating point number that controls the step-size decay^6^. The greater τ is, the less the points are propagated. When τ = 0, the step-size is constant *η* = *δ*.

### Parallelized gradient descent implementation using subvolumes

For the CyCIF segmentation (Fig. 7a,b), the image volume and associated data: the foreground binary and reconstructed 3D gradient map were tiled with subvolumes of (256, 512, 512), with 25% spatial overlap between adjacent subvolumes. Within each subvolume, 3D gradient descent with momentum (*μ* = 0.98) and gradient decay (τ = 0.0) was run for 250 iterations, and stepsize *δ* = 1 to propagate foreground coordinates towards their attractor. At each iteration, coordinates were clipped to lie within the bounds of the subvolume. The final coordinates were converted to global image coordinates by adding the offset position of the subvolume.

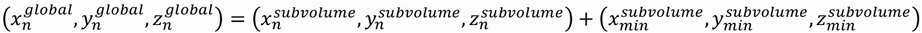

where 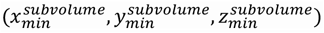 is the corner corresponding to the (0,0,0) origin of the subvolume. Binary thresholding and optimized connected component analysis^90^ is applied to the global coordinates, pooled from all subvolumes, to generate a single globally consistent 3D segmentation. We find this procedure requires no further stitching across subvolumes provided the subvolume size covers a few cells and the spatial overlap enables the attractor basin to be represented in adjacent subvolume tiles.

### Image-based connected component analysis for identifying the unique number of cell centers for instance segmentation

The method is depicted in Extended Data Fig. 4 for a 2D image and described here for a 3D image. Step (i), the final (*t* = *T*) gradient descent advected foreground coordinate positions, 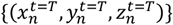 is rasterized onto the image grid by flooring, i.e. 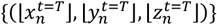 and clipping coordinate values to lie within the bounds of the *L* × *M* × *N* image volume i.e. 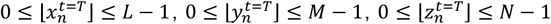 in Python. Step (ii), the number of points at each voxel position is tabulated, each point contributing +1 count. Step (iii), the counts image is Gaussian filtered with *σ* = 1 as a fast approximation to the Gaussian kernel density to produce a point density heatmap, *ρ*(*x*, *y*) for 2D and *p*(*x*, *y*, *Z*) for 3D. This step accounts for gradient errors and spatially connects points into a cluster in a soft manner. The larger the width of the Gaussian filter *σ* the more nearby points will be grouped into the same cluster. This is helpful when segmenting branching structures as highlighted in the coCATs labelled example in Fig. 7d-f. (iv) The density heatmap is sparse, allowing all unique clusters to be identified using a mean threshold with an optional tunable offset specified as a constant multiplicative factor, *k* of the standard deviation (std) of *ρ, threshold* = *mean*(*ρ*) + *k*. *std*(*ρ*). For examples in this paper we set *k* = 0. Image-based connected component analysis is then applied to the binary segmentation of *ρ* to create the distinct spatial cluster segmentation image at *t* = *T* ; *L*^*t*=*T*^(*x*, *y*) for 2D and *L*^*t*=*T*^(*x*, *y*, *Z*) for 3D. Each foreground coordinate, 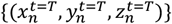 is then assigned to a cluster id by image indexing and the final cell segmentation is computed by mapping the labels of points to their initial positions, 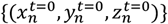. For connected component analysis, u-Segment3D uses the optimized, parallel implementation developed by the Seung Lab (https://github.com/seung-lab/connected-components-3d^90^).

### Module 3: Postprocessing the 3D consensus segmentation

Described below are the implemented postprocessing methods that can be applied to the initial consensus 3D segmentation (module 2). The recommended sequential u-Segment3D workflow is: i) removal of implausible predicted cells involving ia) removal of predicted cells below a user-specified size limit (in voxels), ib) the removal of segmented cells with recomputed 3D gradients inconsistent with that of the 2D-to-3D reconstructed gradients and ic) the removal of cells that are statistically too large (volume > mean(volumes) + *k*. std(volumes) where *k* is a multiplicative factor, default *k* = 5); ii) label diffusion to smooth, enforce the spatial connectivity constraint of segmentation and propagate the initial consensus segmentation to better adhere to the desired features within the given guide image; iii) guided filter to refine and transfer missing local, high-frequency subcellular structures to the segmentation.

The guided image used in label diffusion and guided filtering does not need to be the same as the raw image. Generally, it is a version of the raw where desired cellular features are enhanced.

### (i) Removal of implausible predicted cells

#### (ia) Removal of predicted cells that are too small

Volume of individual cells are computed as number of voxels. The ids of all cells with volume less than the user-specified threshold (default 200) are removed by setting their voxels to 0. Additionally, each cell is checked whether they comprise multiple spatially disconnected components. If so, only the largest component is retained as each segmented cell should be spatially contiguous.

#### (ib) Removal of predicted cells inconsistent with the reconstructed 3D gradients

The reconstructed 3D gradients, ∇Φ_3*D* segmentation_ are computed from x-y, x-z, y-z views of the assembled consensus 3D segmentation. The mean absolute error with the predicted 3D gradients, ∇Φ_3*D*_ used as input in gradient descent is computed per cell, *MAE*_*cell*_ = *mean*(|∇Φ_3*D* segmentation_ − ∇Φ_3*D*_|)_*cell*_. If *MAE*_*cell*_ > user-defined threshold (default 0.85 for *σ*_*post*_ = 1). If the post Gaussian filter *σ*_*post*_ used when fusing gradients from orthoviews is >1, the threshold may need to be relaxed i.e. threshold > 0.85.

#### (ic) Removal of predicted cells that are statistically too large

Ratiometric uneven illumination correction may unduly amplify background at the borders of the image, potentially resulting in the erroneous segmentation of very large background regions. Also in dense tissue, when staining is inhomogeneous and weak, multiple closely packed cells may be segmented as one in the initial 2D segmentation. Assuming cell volumes are approximately normally distributed, we filter out improbably large cells by using the mean and standard deviation (std) of all segmented cell volumes to set a cutoff. Only cells with a volume smaller than mean(volume) + *k*. std(cell volumes) are then retained, with *k* = 5 as default.

### (ii) Labels diffusion to smooth and propagate cell segmentation with spatial connectivity constraint

Labelspreading^88^ is a semi-supervised learning method developed to infer the label of objects in a dataset given the labels of a partial subset of the objects. It works by diffusing labels after one-hot encoding on an affinity graph between objects. u-Segment3D adapts this algorithm for cell segmentation. To be computationally scalable for large cell numbers, for each cell mask, *M*_*i*_, a subvolume, *V*_*i*_, is cropped with the size of its bounding box isotropically padded by a default of 25 voxels. Every label in *V*_*i*_ is one-hot encoded to form a label vector *L* ∈ ℝ^*N*×*p*^ where *N* is the total number of voxels and *p* the number of unique cell ids, including background. We then construct an affinity matrix, *A* between voxels as a weighted sum (weight, *α*) of an affinity matrix constructed from the intensity differences in the guide image, *I* between 8-connected voxel neighbors, *A*_*intensity*_ and another based solely on the voxel spatial connectivity, *A*_*laplacian*_:

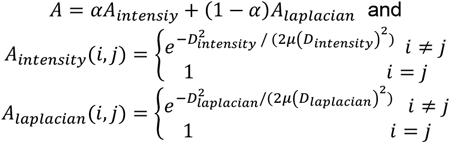

*D*_*intensity*_ is the pairwise absolute difference in intensity values between two neighboring voxels *i* and *j. D*_*laplacian*_ is the graph Laplacian with a value of 1 if a voxel *i* is a neighbor of voxel *j*, and 0 otherwise. *μ*(*D*) denotes the mean value of the entries of the matrix *D*. The iterative label diffusion is then given by

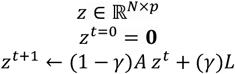

where *t* is the iteration number, 0, the empty label vector and γ is a ‘clamping’ factor controlling the extent by which the original labeling is preserved. The final *Z* is normalized using the softmax operation, and argmax is used to obtain the final cell ids. The refined cell mask, 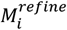 for cell id *i* includes all voxels where the final *Z* is assigned to the same cell id *i*. Multiprocessing is employed to refine all individual cells in parallel. It is recommended to set the parameters per dataset, depending on the extent of correction required. We typically start with a conservative *α* = 0.5, γ = 0.75, and run the propagation for 25 iterations, and adjust accordingly. The guide image, *I* is usually the normalized input image (after any preprocessing) to the 2D segmentation but can be any processed image that enhances the desired cell features. For additional speed, particularly for tissue, we would typically treat each cell mask, *M*_*i*_ independently as binary without considering the multi-label setting of jointly refining neighboring cell masks in the cropped volumes.

### (iii) Guided image filtering to recover missing high-frequency features and subcellular protrusions

The guided filter^89^ is a filter that can be implemented in linear time, to efficiently transfer features in a guidance image, *I* to the input image to be filtered, *P*. Setting *I* to the ridge-filtered input image which enhances high-frequency cellular protrusion and vessel features, and *P* to be the binary mask of cell *i*, the resulting filtered output *Q* is a ‘feathered’ binary, where transferred image features appear as an alpha matte locally around the mask boundaries. The radius of the boundary that is refined is controlled by a radius parameter, *r* = 35 voxels (by default), and the extent of transfer by a regularization parameter, *ϵ* = 1 × 10^−4^. We find the binary mask can encapsulate the cell relatively coarsely. The stronger the features are enhanced in *I* the more prominent the transferred structure. *Q* is then re-binarized using multi-level Otsu thresholding. Typically, we use the two-class binary Otsu. As for label diffusion, guided filtering is applied to cropped subvolumes, *V*_*i*_ of individual cells, with the size of individual bounding boxes isotropically padded by a default 25 voxels. For computational efficiency, for touching cells, we perform the guided filter segmentation independently for each cell and mask out spatial regions occupied by surrounding cell ids. More accurately, we could obtain the guided filter response for all cell ids in the subvolume and use argmax to define the maximum filter response. Multiprocessing is used to perform the guided filter refinement to all cells in parallel. The radius *r* sets the maximum protrusion length that can be recovered. If the cell density is high, it may not be possible to adjust *r* to recover long protrusions without erroneously incorporating features of neighboring cells. Nevertheless, the guided filter result may assist the application of matching algorithms or serve as an improved seed image for watershed algorithms in further specialized downstream processing.

### Semi-automatic tuning of diameter parameter in Cellpose models

The process is illustrated in Extended Data Fig. 11a. for 3D and described below.

#### Determining the optimal diameter for 2D image

Given a pixel neighborhood size with isotropic width, *P* pixels, we conduct a parameter scan of diameter = [*d*_*low*_, *d*_*high*_] (typically *d*_*low*_ = 10, *d*_*high*_ = 120) at equal increments of 2.5 or 5. For each diameter, a contrast score is computed measuring the ‘sharpness’ of the Cellpose model predicted 2D x- and y- gradients (∇_*x*_Φ and ∇_*y*_Φ respectively) and optionally the normalized cell probability map, *p* (0-1).

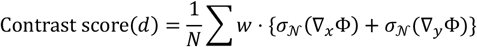

where *N* is the total number of image pixels, *w* is a pixelwise weight set to be *p* and *σ*_*N*_(*I*) is the pixelwise local standard deviation of the image *I* evaluated over the isotropic local neighborhood of width *P* pixels. *p* is computed from the unnormalized raw cell probabilities after clipping to range [-88.72, 88.72] (to prevent overflow or underflow in IEEE float32) by applying the transformation,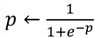. The result is a contrast score function of *d*. A centralized moving average of 5 (if diameter increment is 2.5) or 3 (if diameter increment is 5) is applied to smooth the contrast score function. The diameter *d* that maximizes the contrast score is used as the optimal diameter, *d*_*opt*_ in the Cellpose model. We generally observe no difference in *d*_*opt*_ between *w* = 1 or *w* = *p* for Cellpose models.

#### Determining the optimal diameter for 3D volume

If cells exhibit large size variations slice-by-slice, the optimal diameter determination for 2D should be applied slice-by-slice (Fig. 5). For large numbers of slices this is slow. As compromise, we find good performance for many datasets, if we set an optimal diameter using a single representative 2D slice and apply it to the 3D volume for each orthoview. This representative 2D slice is set automatically using (i) the most in-focus slice as determined by the highest mean sobel magnitude, (ii) the slice with highest mean intensity, (iii) the mid-slice, or (iv) be user-defined.

### Compared 2D-to-3D stitching methods

#### CellComposer^50^

We modified the *process_segmentation_masks* from the official code repository (https://github.com/murphygroup/3DCellComposer/blob/main/run_3DCellComposer.py) to allow for stitching from only x-y, x-z and y-z cell mask inputs. We do not use the unsupervised segmentation metric to set optimal Jaccard index overlap (JI) for stitching. Instead, we perform a search, stitching for JI at 0, 0.1, 0.2, 0.3, 0.4, the same as hard-coded in the code, and report the AP curve with highest mean value.

#### CellStitch^71^

We used the *full_stitch* function from the official code repository to stitch input x-y, x-z and y-z cell masks, https://github.com/imyiningliu/cellstitch/ as illustrated in the provided example notebooks, https://github.com/imyiningliu/cellstitch/tree/main/notebooks. This function is parameterless.

### Other tested segmentation methods

#### Cellpose 3D mode with pretrained models

We ran pretrained cellpose models in 3D mode to generate 3D segmentations by setting do_3D = True. As we find this mode prone to oversegment and Cellpose 3D only allows one diameter for all orthoviews, we used the largest diameter inferred by our contrast score function. Models were run twice. The first time was to obtain the raw, unnormalized cell probability image, which was then used to determine the binarization threshold. We then ran a second time using the determined threshold to generate the 3D segmentation. We then additionally removed all cells with volume < 2500 voxels to get a segmentation that maximizes the measured average AP.

#### Omnipose 3D

We ran the pretrained *plant_omni* model following the example in the documentation (https://omnipose.readthedocs.io/examples/mono_channel_3D.html). This model operates on the raw image downsampled by a factor 1/3 in all dimensions and does not reinterpolate the raw image to isotropic resolution. No other preprocessing was used. We found the raw output to predict many small objects leading to an artificially low AP when compared with qualitative assessment. We therefore removed all objects with volume < 2500 voxels to get a segmentation with maximum average AP. Specifying a higher size cutoff led to lower mean AP, as it detrimentally affected lateral primordial images comprising largely smaller cells in the dataset.

#### Cellpose 3D mode with Omnipose trained ‘plant_cp’ model

We ran the 2D pretrained Cellpose *plant_cp* model using the same function call as the example in the Omnipose documentation for *plant_omni* but with omni=False and do_3D=True. As in the case of the Omnipose 3D *plant_omni* model, we found many small objects were predicted and additionally postprocessing the output segmentation, removing all objects with volume < 2500 voxels.

#### PlantSeg 2D and 3D

We applied PlantSeg^7^ models pretrained on 2D slices and 3D volumes of the Lateral Root Primordia (LRP) and Ovules dataset as-is without any custom preprocessing of downloaded images. The ‘lightsheet_2D_unet_root_ds1x’ and ‘lightsheet_3D_unet_root_ds1x’ models were used for 2D and 3D segmentation of Lateral Root Primordia respectively. The ‘confocal_2D_unet_ovules_ds2x’ and ‘confocal_3D_unet_ovules_ds2x’ models were used for 2D and 3D segmentation of Ovules respectively. We use a tile size of (1,96,96) for 2D and (128,160,160) for 3D with ‘MultiCut’ and watershed in 2D or 3D respectively with postprocessing ‘on’ to produce segmentations. Multiple non-zero labels corresponded to image background in 2D predicted segmentations. To not unfairly penalize PlantSeg, for 2D evaluation and consensus segmentation with u-Segment3D, we reassigned all 2D predicted labels with more than 0.7 spatial overlap with the reference background as ‘background’. This problem was not present PlantSeg3D where we could assign the largest contiguous label as ‘background’. PlantSeg 2D and 3D were evaluated on the test splits of Ovules and LRP. 2D test were constructed from provided 3D splits as described below for the training of segmentation models.

### Training of 3D segmentation models

#### Dataset construction

Datasets were resized to be isotropic voxels with the same dimensions as used for the pretrained Cellpose experiments in Fig 3. We use the existing train/test/val split for each dataset when provided. For datasets without test split, and those without a val split, all of which come from EmbedSeg^15^, we followed the splitting procedure used in the official EmbedSeg code repository for each dataset, (https://github.com/juglab/EmbedSeg/tree/main/examples/3d). This procedure first splits 10% of the train data as test, if required and the 15% of the remainder train data as validation. All 3D models were trained on the constructed train/val datasets and evaluation reported on the test dataset.

#### Embedseg3D

We trained EmbedSeg 3D models following the provided notebooks for each dataset (https://github.com/juglab/EmbedSeg/tree/main/examples/3d), using the model with best IoU after 200 epochs. For Ovules and Lateral Root Primordia, we followed the example for the Arabidopsis CAM dataset.

#### StarDist3D

We trained StarDist3D^47^ models for 400 epochs and 96 rays following the example code, (https://github.com/stardist/stardist/tree/main/examples/3D). For datasets with high shape anisotropy - Mouse skull nuclei and Lateral Root Primordia or large cell sizes - Arabidoposis and Ovules, we isotropically downsampled the volume and if required, additionally doubled the subsampled grid to ensure the neural network field of view was greater than median object size.

### Training of 3D segmentation models guided by u-Segment3D consensus segmentations

A new Embedseg3D model was trained for Platynereis nuclei using the same train/val data splits but using the u-Segment3D consensus segmentation of the trained Embedseg2D model as the reference 3D instance masks. We do not want to train to convergence. Instead, we stopped training as the IoU on the val split begins to plateau (25 epochs). The best IoU model was then evaluated on the test data split using the actual reference segmentation.

### Training of 2D segmentation models

#### Dataset construction

2D train/test/val splits were derived based on the 3D train/test/val volumes constructed for 3D segmentation. For each 3D volume, we equisampled 15% of the 2D slices between the slice containing the first cell and that containing the last cell. For example if the reference segmentation x-y slices has a cell starting from z=15 and has a cell up to z=225, we would sample every 15^th^ from 15. This was done for x-y, x-z, y-z views independently for each volume. 2D slices sampled from train/test/val 3D volumes formed the respective 2D train/test/val datasets that all 2D models were trained on.

#### EmbedSeg2D

We trained EmbedSeg2D models for each dataset following the example codes for dsb-2018, (https://github.com/juglab/EmbedSeg/tree/main/examples/2d/dsb-2018). In 2D, which yields thousands of cells (Extended Data Fig. 15), we found that the instance embedding-based training approach of EmbedSeg2D was time-consuming and observed marginal performance gains after just a few epochs. Therefore we only ran the full 200 epochs on the smaller datasets with fewer cells: Mouse organoids, Mouse skull nuclei, Platynereis ISH nuclei and Platynereis nuclei. For the remainder, we ran sufficient epochs (minimum 35 epochs) to observe plateauing and slow-down of the validation IoU. We also rederived the train/val at 5% equisampling, filtering out 2D slices that did not contain a sufficient number of foreground pixels. Performance was still evaluated on the original constructed 2D test dataset equisampled at 15% for all datasets.

#### StarDist2D

We trained StarDist2D models following the tutorial notebook, https://github.com/stardist/stardist/blob/main/examples/2D/2_training.ipynb with 32 rays. A unique feature of StarDist models is the ability to enforce a star-convex shape prior. Therefore, we trained StarDist2D for all datasets first with the parameter *train_shape_completion=True*. However we found this was detrimental for mouse skull nuclei, lateral root primordia and Ovules whose cells in 2D slices were presumably too oblong and/or concave. ‘Trained’ models only predicted cell centroids with a circle of the same radius for each cell. For these datasets we set *train_shape_completion=True*, and further filtered out all 2D images in train/val that did not contain a sufficient number of foreground pixels. Performance was still evaluated on the original constructed 2D test dataset.

#### Cellpose2D

We trained the latest Cellpose3^62^ ‘cyto3’ model and current default model using the *train_seg* function in single channel mode, (channels=[0,0]), following the official API documentation, (https://cellpose.readthedocs.io/en/latest/train.html) for the default 100 epochs. For ovules, we trained for 250 epochs. To generate 2D instance segmentation masks, (Extended Data Fig. 15), we used the default parameters for Cellpose. The equivalent u-Segment3D generation (Cellpose(u)) uses the same parameters but now with u-Segment3D’s gradient descent and connected component clustering implementation with parameter setting of 50 gradient descent iterations, gradient decay 0.1, and automatically inferring the binary cell foreground threshold as the higher of a minimum threshold 0.25 and the higher threshold from applying 3-class Otsu thresholding to the normalized predicted cell probability. The minimum threshold is used to suppress generating spurious segmentations when the input 2D image is of only background without cells.

### u-Segment3D with trained 2D segmentation models

For a dataset, we used the same parameter settings for each trained 2D model and PlantSeg2D. The main u-Segment3D parameters to consider when using the indirect method with 2D segmentation masks is i) choice of distance transform, ii) gradient decay, iii) gradient descent iterations and iv) minimum cell size. For these we used i) the explicit medial point diffusion distance transform, ii) gradient decay of 0.01, iii) 50 gradient descent iterations and iv) minimum cell size cutoff of 50. These base parameters were rationally adapted to generate the consensus 3D cell segmentation in individual datasets as follows. In case of sporadic segmentation across consecutive 2D slices we used a neighborhood size of 3×3×3 pixels for content-based averaging of foreground (Extended Data Fig. 2) for all datasets and not varied.

#### Ovules

Cells are densely packed. We used an increased 250 gradient descent iterations.

#### Lateral Root Primordia

Cells are densely packed. There are both cells that are small and convex-like and be highly elongated with long branches. We use as distance transform the explicit Poisson diffusion distance with medial skeleton as source (Extended Data Fig. 1), gradient decay, τ = 0.1, and an increased 100 gradient descent iterations.

#### Arabidopsis-CAM

Cells are densely packed and have small volume in voxels. We used an increased 100 gradient descent iterations.

#### Mouse organoids

Cells are not densely packed, and not branching. Base parameters were used without modification.

#### C.Elegans

Cells are more densely packed than mouse organoids. We used an increased 100 gradient descent iterations.

#### Mouse skull nuclei

Nuclei are not densely packed, and not branching. Base parameters were used without modification.

#### Platynereis ISH nuclei

Nuclei are small, but not densely packed, and not branching. Base parameters were used without modification. We used a decreased minimum cell size cutoff of 25.

#### Platynereis nuclei

Nuclei are small, some touching larger nuclei, and not branching. We used an increased 100 gradient descent iterations, and a decreased minimum cell size cutoff of 25.

### Evaluation of segmentation quality

#### Segmentation quality in single images

For single 2D and 3D images, we find the optimal matching between predicted and reference cell segmentations. Given a total number of *M* predicted cells, and *N* reference cells, we iterate and find for each predicted cell *i*, its *K*-nearest reference cells according to the distance between their centroids. For each of the *K*-nearest reference cells, we compute the intersection-over-union (IoU) metric (0-1) (see below). This produces an *IoU*(*i*, *j*) ∈ ℝ^*M*×*K*^ matrix. We convert this to a distance cost matrix, *dist*(*i*, *j*) = 1 − *IoU*(*i*, *j*) ∈ ℝ^*M*×*K*^. The optimal matching between predicted and reference cells is then found by solving the linear sum assignment using a modified Jonker-Volgenant algorithm^121^ implemented by Python Scipy, scipy.optimize.linear_sum_assignment and retaining only the pairings that overlap spatially (*IoU*(*i*, *j*) > 0). The segmentation quality for an image was then assessed by (i) the mean IoU, to measure the spatial overlap of matched predicted and reference cells and (ii) the F1 score (see below), i.e. the harmonic mean of precision and recall to measure how accurately the reconstructed segmentations detected only reference cells.

#### Intersection-over-union (IoU)

Also called the Jaccard index, is defined as the total number of pixels in the intersection divided by the total number of pixels in the union of two binary segmentation masks *A* and *B*, 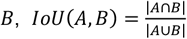.

#### F1 score

Predicted cells that are validly matched to a reference cell (IoU>0) were defined as true positives, TP. Predicted cells that are not matched to a reference cell are false positives, FP, and reference cells that are not matched are false negatives, FN. The precision is the number of matched cells divided by the total number of predicted cells, 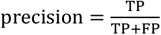 .The recall is the number of matched cells divided by the total number of reference cells, 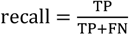. F1 score is the harmonic mean of precision and recall, 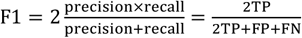

### Average precision curve

We evaluate the quality of cell segmentation using average precision, consistent with popular segmentation models such as StarDist^2^, Cellpose^5^ and Omnipose^6^. Each predicted cell label mask is matched to the reference cell label mask that is most similar, as defined by IoU. The predictions for an image are evaluated at various levels of IoU. At a lower IoU, a predicted cell can overlap a reference cell with fewer pixels to determine a valid match. For a given IoU threshold, the valid matches define the true positives, TP, the predicted cells with no valid matches are false positives, FP, and the reference cells with no valid matches are false negatives, FN. Using these definitions, the average precision metric (AP) for a segmented image is:

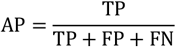

The average precision (AP) curve is reported for a dataset by averaging over the average precision metric for each image in the dataset for a range of IoU thresholds. Optimal matching of predicted and reference cells is too computationally demanding in 3D even when restricting the search to nearest neighbors. Consequently, we use the same approximate matching as in Cellpose derived from the fast matching functions in StarDist. We find this fast matching is not invariant to cell id permutation. To compute the correct AP, we first relabel all cells sequentially after performing an indirect stable sort based on their (*x*, *y*, *Z*) centroids for both reference and predicted cell segmentation independently. In line with Cellpose, the AP curve is reported for 11 IoU thresholds equisampling the range [0.5,1.0]. Many datasets e.g. Ovules do not rigorously label every cell in the image but only the cells of the primary, single connected component object in the field of view. In contrast, pretrained Cellpose models predict all cells in the field-of-view. For fair evaluation, for these datasets (all except for Embedseg skull nuclei, *Platynereis* nuclei and *Platynereis* ISH nuclei), we use the reference segmentation to define the foreground connected components to evaluate AP and include all predicted cells part of binary foreground spatial connected components that share at least 25% overlap with a reference connected component. For DeepVesselNet, we use at least 1% overlap due to the thinness of vasculature.

### F1 curve

We use the F1 curve as an additional segmentation performance measure. We compute the F1 curve for a dataset similar to the average precision curve, by averaging the F1 score metric for each image in the dataset for a range of IoU thresholds for a range of IoU thresholds. We use the same matching procedure and IoU thresholds as for the AP curve. The F1 is correlated to AP that assigns greater weight to true positives.

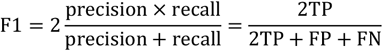

### Visualization

The Fiji ImageJ^122^ 3D viewer plugin was used to render 3D intensity and segmentation image volumes. To visualize the intensity in Fig. 6, we acquired a snapshot of the rendering, then applied an inverse lookup table to the snapshot. Surface meshes in Fig. 6 were extracted using u-Unwrap3D^72^ and visualized using MeshLab^123^. Rotating surface mesh movies were created using ChimeraX^124^.

## Supplementary Figures

**Extended Data Figure 1.**
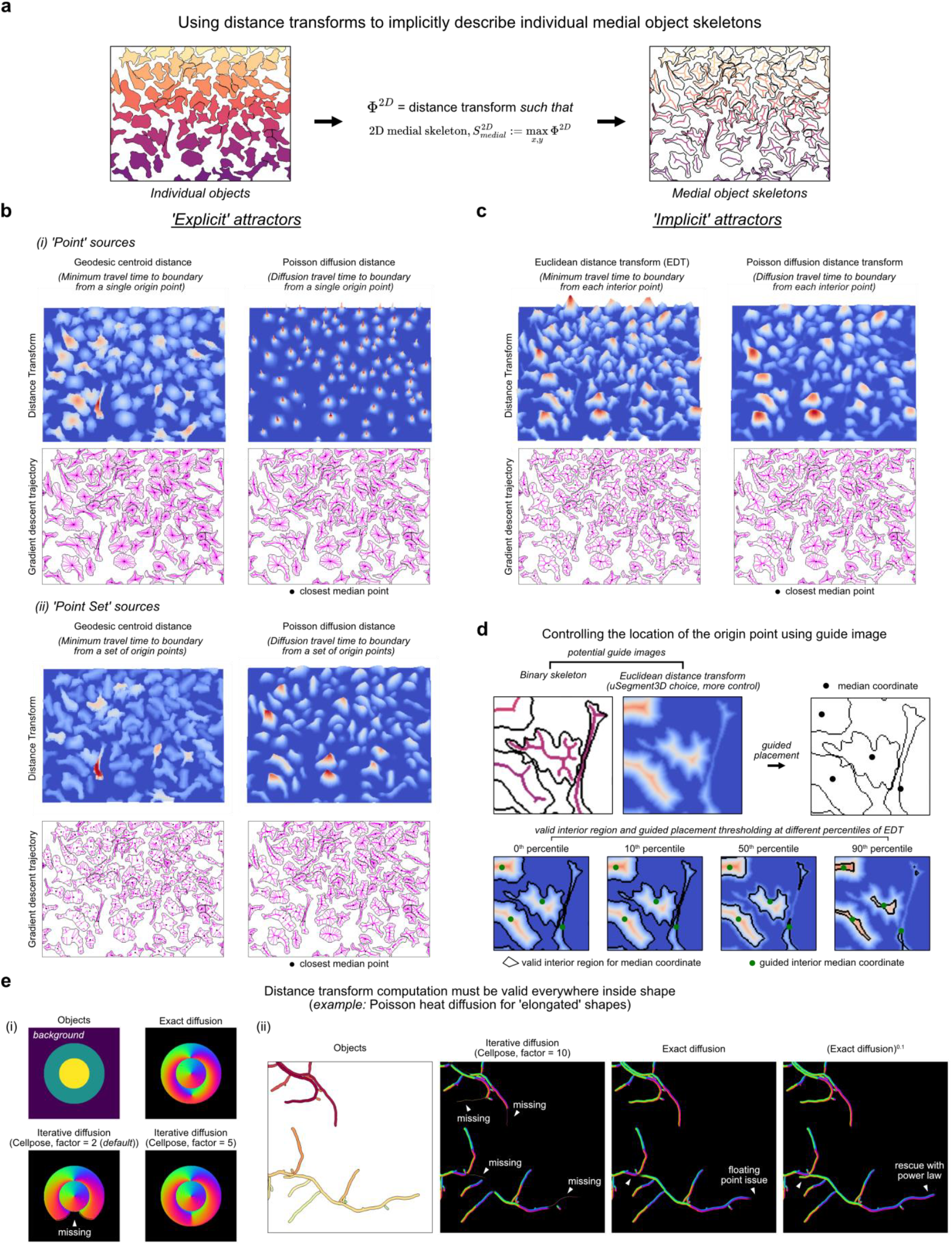
2D distance transforms for consensus 2D-to-3D segmentation in u- Segment3D. **a)** Necessity of 2D distance transforms to sample the 2D medial skeleton of individual shapes. **b)** Example of first-order (geodesic, left) and second-order (diffusion, right) distance transforms with ‘explicit’ attractors implemented in u-Segment3D. The limit of a gradient descent is explicitly defined as either (i) a single point-source on the 2D medial skeleton or (ii) a set of points along the 2D medial skeleton. **c)** Example of first-order (geodesic, left) and second-order (diffusion, right) distance transforms implemented in u- Segment3D with ‘implicit’, skeletal-based attractors. Contrary to transforms with explicit attractors, here the gradient descent may not necessarily converge to the limit. For each example in **b), c)**, distance transforms are represented as a topographic relative heat map, colored blue (lowest) to red (highest) (top) by distance from the cell boundary, and by trajectories (magenta) of equi-sampled boundary points converging to the attractor under gradient descent (bottom). Black point = closest internal point to the median shape coordinate. **d)** Illustration of using percentile-based thresholding of the Euclidean distance transform of individual cells as a soft constraint to find the medial centroid 2D coordinate for convex and concave shapes used in computing the 2D point-source distance transforms in c). **e)** Unit-normalized 2D gradients for circle in doughnut synthetic shape. **f)** Unit-normalized 2D gradients for (i) elongated touching bacterial shapes using (ii) the simulated diffusion approach as implemented in Cellpose versus (iii) the exact solution of diffusion equation with boundary conditions as implemented in u-Segment3D (see Methods). Using an exact solver, computed gradients remain defined and are never zero, even in very long structures. However, due to floating-point precision, the gradient magnitude far from the source is no longer unit length, affecting the convergence of gradient descent. Thanks to the non-zero gradient magnitudes, u-Segment3D can rectify this situation, by (iv) raising the distance transform by an exponent *p*, Φ^*p*^ before computing the gradient. 2D gradients in panels (ii-iv) are colored according to direction. The number of simulated diffusion steps in Cellpose varies per shape and equals the factor multiplied by the number of pixels covered by the shape.

**Extended Data Figure 2.**
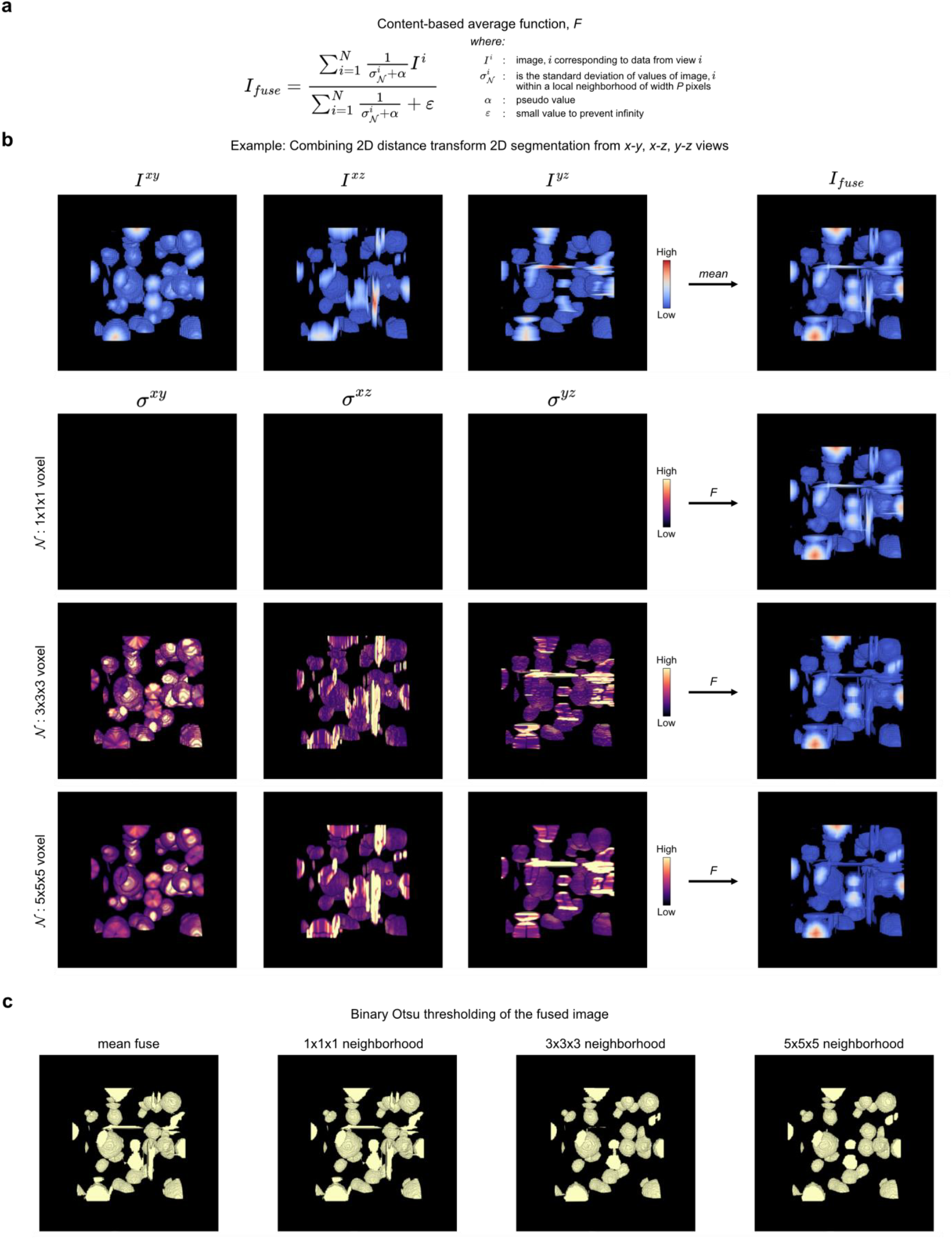
Content-based average function for combining data from multiple views for consensus 2D-to-3D segmentation. **a)** Mathematical definition of the content-based average function as the inverse local variance weighted mean of input image values in an isotropic neighborhood of width *P* pixels. **b)** Example of applying the average function defined by a) to fuse the stacked 2D distance transforms after 2D slice-by-slice segmentation of a volume of spherical cells from x-y, x-z and y-z views. 1^st^ row: Left-to-right, the Euclidean distance transform colored blue (low) to red (high) from the three orthoviews and fused distance transform using pixelwise mean. 2^nd^ row: Left-to-right, the per-pixel local variance weight image *σ* for each orthoview for a neighbourhood of width *P* = 1 pixel and resultant fused distance transform using content- based averaging. 3^rd^ row: Left-to-right, the per-pixel local variance weight image *σ* for each orthoview for a neighborhood of width *P* = 3 pixel and resultant fused distance transform using content-based averaging. 4^th^ row: Left-to-right, the per-pixel local variance weight image *σ* for each orthoview for a neighborhood of width *P* = 5 pixel and resultant fused distance transform using content-based averaging. **c)** Result of applying binary Otsu thresholding on the fused distance transform based on pixelwise mean, and content-based averaging using a neighborhood of width *P* = 1, 3, 5 pixels from left-to-right.

**Extended Data Figure 3.**
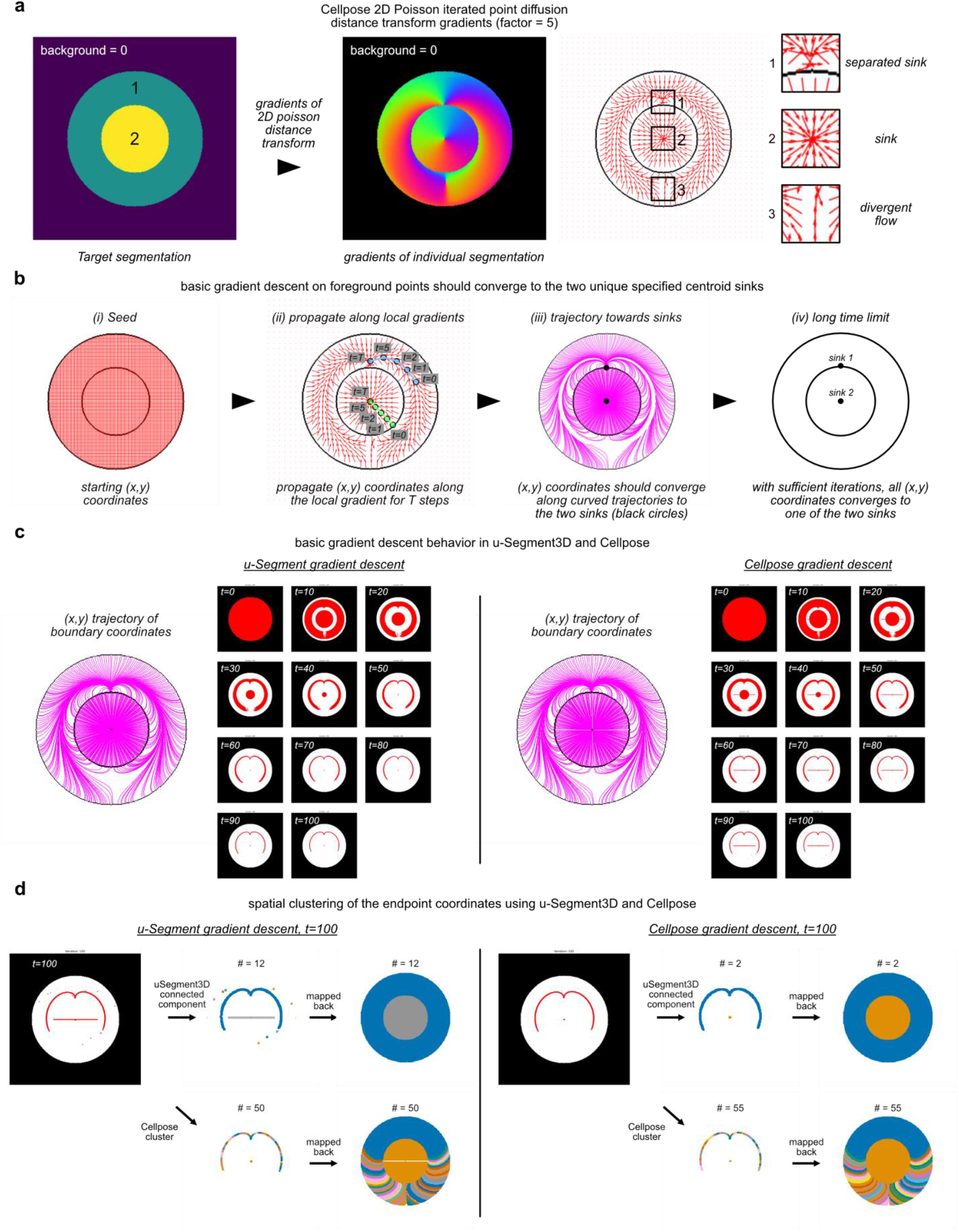
Illustration of 2D Gradient descent for reversible 2D shape erosion. **a)** Cellpose computed 2D gradients for a synthetic shape of a doughnut (labelled 1) surrounding a circle (labelled 2) (left) with the 2D gradients visualized as red arrows (right) and zoomed in at three regions with different types of flow behavior. The number of simulated steps equals the factor = 5 multiplied by the number of pixels occupied per shape. **b)** Schematic of the expected behavior when gradient descent is iteratively applied to propagate the initial foreground (x,y) coordinates with the 2D gradients, the limit being convergence to the 2 black centroids. **c)** Observed point trajectories (magenta lines) and snapshots of coordinate positions (red points in images) running gradient descent in u-Segment3D (left) vs Cellpose (right) for 100 iterations. Red features indicate the point aggregates that emerge after the indicated number of gradient descent iterations. **d)** Recovered cell shapes based on applying u-Segment3D (top) or Cellpose (bottom) spatial proximity clustering on the final coordinate positions after 100 iterations of u-Segment3D (left) or Cellpose (right) gradient descent. The maps demonstrate the critical role optimal numerical implementation of gradient descent plays in avoiding generating severely over-segmented objects.

**Extended Data Figure 4.**
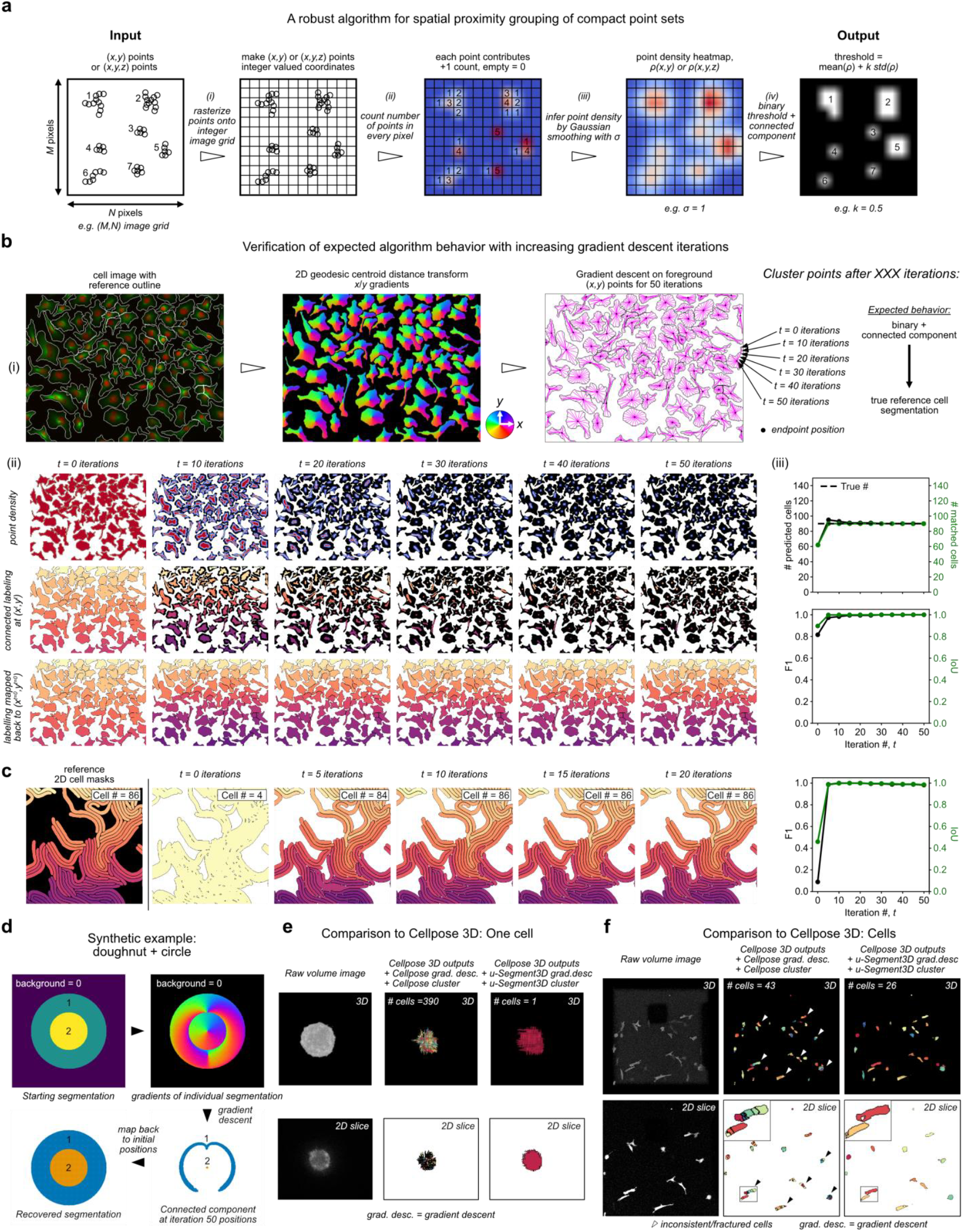
Using image-based spatial connected component analysis to robustly identify distinct spatially compact point sets. **a)** 2D Illustration of the image-based connected components spatial clustering approach in u-Segment3D involving left-to-right, rasterization of floating-point coordinates onto a discrete image pixel grid, building a count of the number of points within each pixel, approximate Gaussian kernel density estimation using a Gaussian filter of *σ*, binary thresholding on the mean density and subsequent connected component analysis to identify distinct spatial clusters. **b)** Verification of algorithmic stability by application to foreground (x,y) coordinates after propagation of 50 iterations with gradient descent. 2D gradients are computed using the 2D geodesic centroid point source distance transform. (i) Schematic of the experimental setup for a 2D cell segmentation with densely touching cells (top panel). (ii) The point density, connected component labelling at current (x^*t*^,y^*t*^) coordinates and labels mapped back to initial (x^*t=0*^,y^*t=0*^) coordinates (top-to-bottom) after t=0, 10, 20, 30, 40, 50 iterations of gradient descent (left-to-right). (iii) Plot of the number (#) of unique cells predicted (left, black-colored y-axis and line) and number of matched cells with reference cell shapes (right, green-colored y-axis and line) with iteration number, *t* (top). Dashed black horizontal line indicates the true cell number. Plot of F1 score of matching with reference cells (left, black-colored y-axis and line) and the mean intersection-of-union (IoU) of matched cells with reference (right, green-colored y-axis and line) with iteration number, *t* (bottom). **c)** Reference 2D cells of elongated touching bacteria (left), identified unique cells by spatial connected component at gradient descent propagated coordinates after t = 0, 5, 10, 15, 20 iterations (middle), and plot of F1 score matching with reference cells (left, black-colored y-axis and line) and the mean intersection-of-union (IoU) of matched cells with reference (right, green-colored y-axis and line) with iteration #, *t*. **d)** Image-based connected component labeling applied to recover a doughnut (region 1) surrounding a circle (region 2) after 50 iterations of gradient descent. **e)** Comparison of segmentations of an isolated noisy single cell^84^ between Cellpose 3D and u-Segment3D based gradient descent and connected component labeling. Left-to-right: 3D rendering of raw volume, 3D segmentation using Cellpose’s gradient descent and density-based clustering, and using u-Segment3D’s gradient descent and image-based connected component clustering (top row) with respective mid x-y slice (bottom row). **f)** Comparison of segmentations of multiple cells with elongated morphologies^84^ between Cellpose 3D and u- Segment3D based gradient descent and connected component labeling. Left-to-right: 3D rendering of raw volume, 3D segmentation using Cellpose’s gradient descent and density-based clustering, 3D segmentation using u-Segment3D’s gradient descent and image-based connected component clustering (top row) with respective mid x-y slice (bottom row). Black arrowheads highlight examples of fractured single cells due to unstable spatial clustering in Cellpose. Inset: zoom-in on two fractured cells.

**Extended Data Figure 5.**
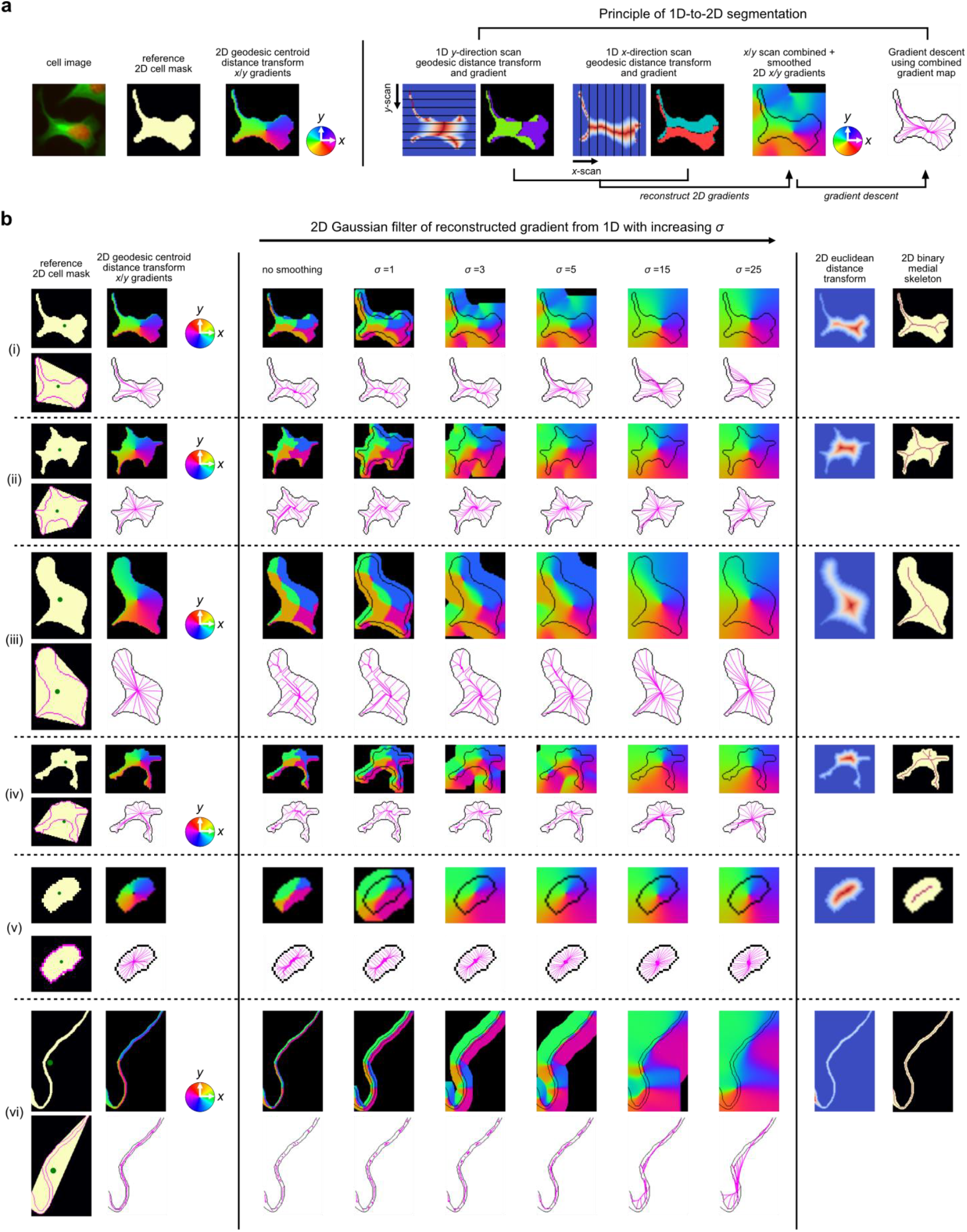
1D-to-2D reconstruction of single cells from ideal 1D segmented stacks. **a)** Schematic of the workflow to investigate 1D-to-2D segmentation. Reference cell shape and ideal 2D geodesic distance transform to reconstruct (left). Reconstruction of 2D gradient field from computed 1D gradients (right). Left-to-right: Determination of the y-direction and x-direction gradient by computing the distance transform of each pixel in 1D slices to the respective slice centroid and taking the gradient; combining the x- and y- gradient into a 2D gradient field and smoothing with a Gaussian *σ*; performing gradient descent on the smoothened 2D gradients to propagate all interior points to a unique centroid (magenta line trajectories). **b)** Gradient descent behavior using Gaussian filter of increasing *σ* on the initial 2D gradients reconstructed for 6 examples of individual cells with diverse morphologies. (i-iii), examples of approximately star-convex shapes; (iv) example of a cell with branches of lengths comparable to cell body; (v) example of convex shaped cell; (vi) example of a thin vessel-like shaped cell. Left column, montage of 4 images depicting reference cell shape and its convex hull image with green point representing the centroid coordinate and the cell’s exact 2D geodesic centroid distance transform with associated gradient descent trajectory to reconstruct. Middle column, observed gradient descent trajectory (magenta line) when the reconstructed 2D gradients are smoothed with an isotropic Gaussian filter of increasing *σ*. Right column, comparison of the gradient descent trajectory with the implicit medial skeleton specified by the 2D Euclidean distance transform and explicit medial skeleton from morphological operations.

**Extended Data Figure 6.**
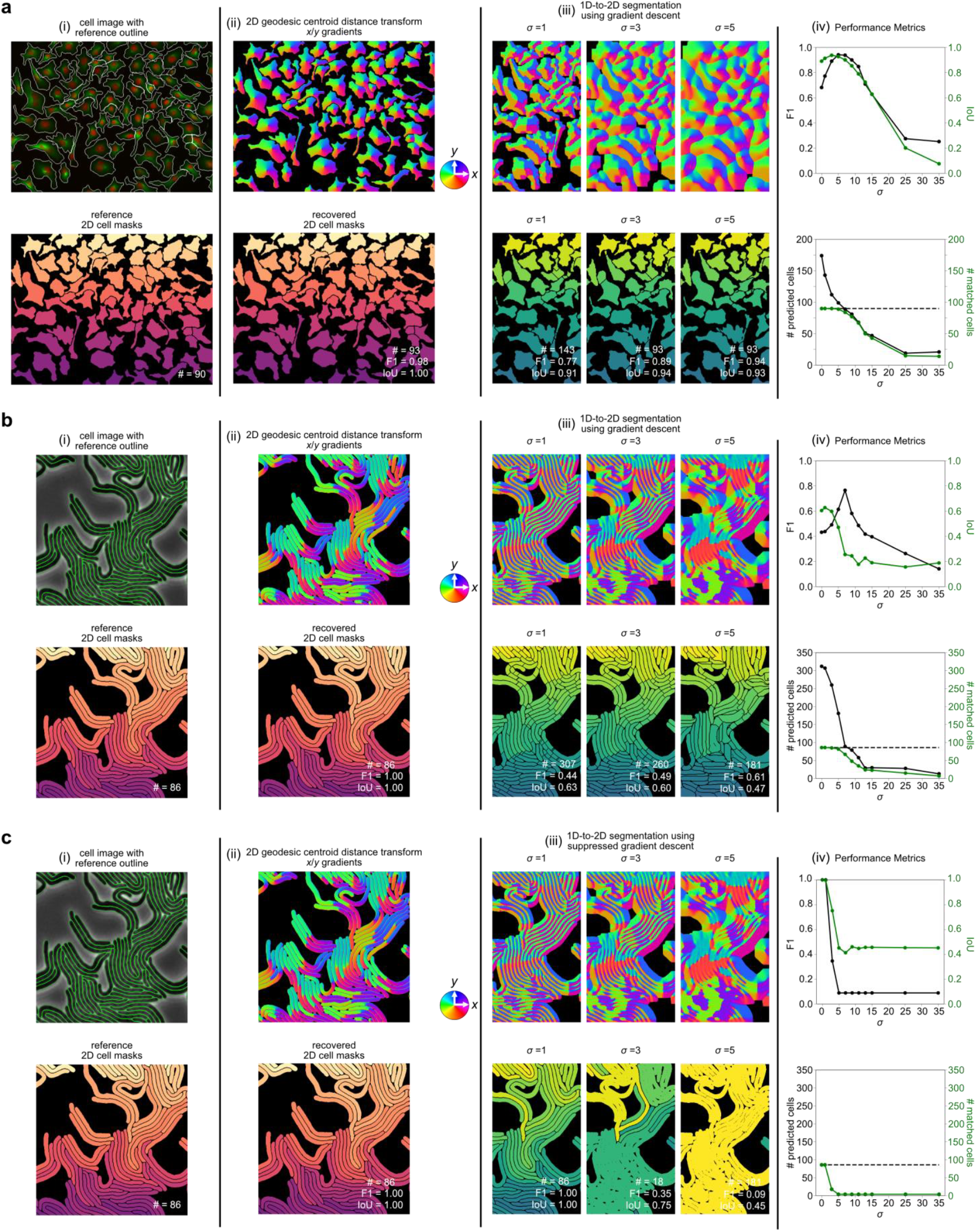
1D-to-2D reconstruction for densely packed cell shapes. **a)** Representative example of a dense 2D cell culture with diverse morphologies with (i) reference cell boundaries overlaid and delineated in white (top) and uniquely colored cell masks (bottom). (ii) Exact unit-normalized 2D gradients of the geodesic centroid distance transform colored by direction (top) and the recovered cell masks using connected component labeling after 100 iterations of gradient descent with step-size of one pixel. (iii) Reconstructed 2D gradient from 1D after Gaussian filtering with increasing *σ* left-to-right (top) and corresponding recovered cell masks using connected component labeling after 100 iterations of gradient descent. (iv) Performance of the 1D-to-2D reconstruction with increasing *σ*. F1 score of matching with reference cells (left, black-colored y-axis and line) and mean intersection-of-union (IoU) of matched cells with reference (right, green-colored y-axis and line) (top). Number (#) of distinct predicted cells (left, black-colored y-axis and line) and # of those that could be matched with reference cells (right, green-colored y-axis and line) (bottom). Dashed black horizontal line indicates the true cell number. **b)** Representative example of a dense 2D cell culture with highly elongated, vessel-like cells. Panels (i)-(iv) as in a). **c)** Same example image and panels (i)-(iv) as in b) but using suppressed gradient descent where the step-size 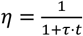, is attenuated with increasing gradient descent iteration number, *t* and τ = 1 (Methods).

**Extended Data Figure 7.**
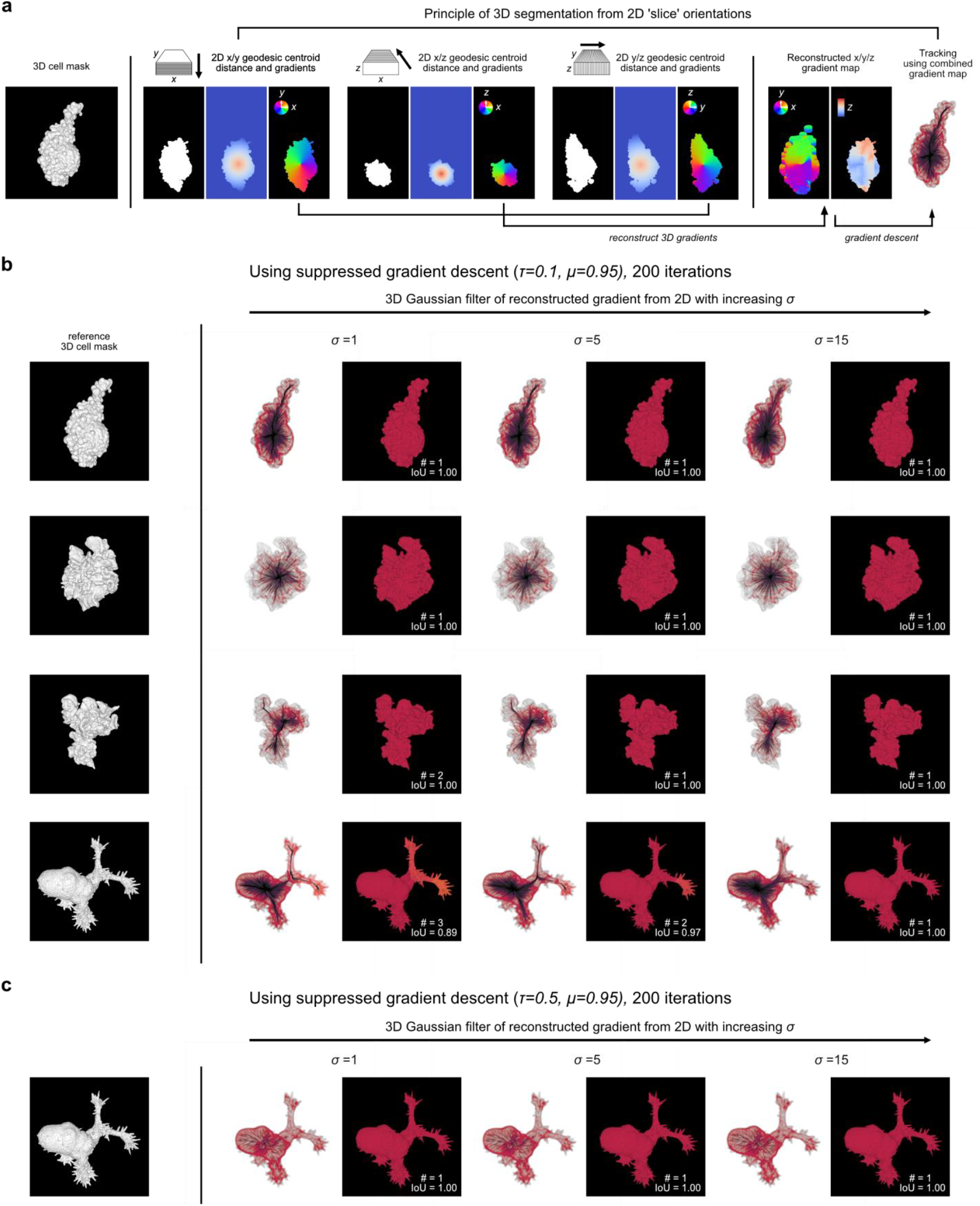
2D-to-3D reconstruction of single cells from ideal 2D segmented stacks. **a)** Illustration of the reconstruction experiment given the 3D shape of a single cell (left), by generating 2D gradients slice-by-slice in x-y, x-z, y-z views, treating each disconnected spatial component as a unique 2D cell (middle) and performing gradient descent on the reconstructed 3D x/y/z gradient followed by connected component labeling on the final advected 3D coordinates (right). **b)** Reconstruction of cells with blebs (1^st^ row), lamellipodia (2^nd^, 3^rd^ rows) and filopodia (4^th^ row). In each row, left-to-right: reference binary 3D cell shape, the 3D gradient descent trajectory (left) and reconstructed 3D shape (right) for Gaussian filtering of the 3D reconstructed gradients with *σ* = 1,5,15, using suppressed gradient descent (τ = 0.1) with momentum (*μ* = 0.95). **c)**. Reconstruction of the same cells with filopodia in b) for post- Gaussian filtering with *σ* = 1,5,15 (left-to-right), using suppressed gradient descent with greater decay (τ = 0.5) with momentum (*μ* = 0.95).

**Extended Data Figure 8.**
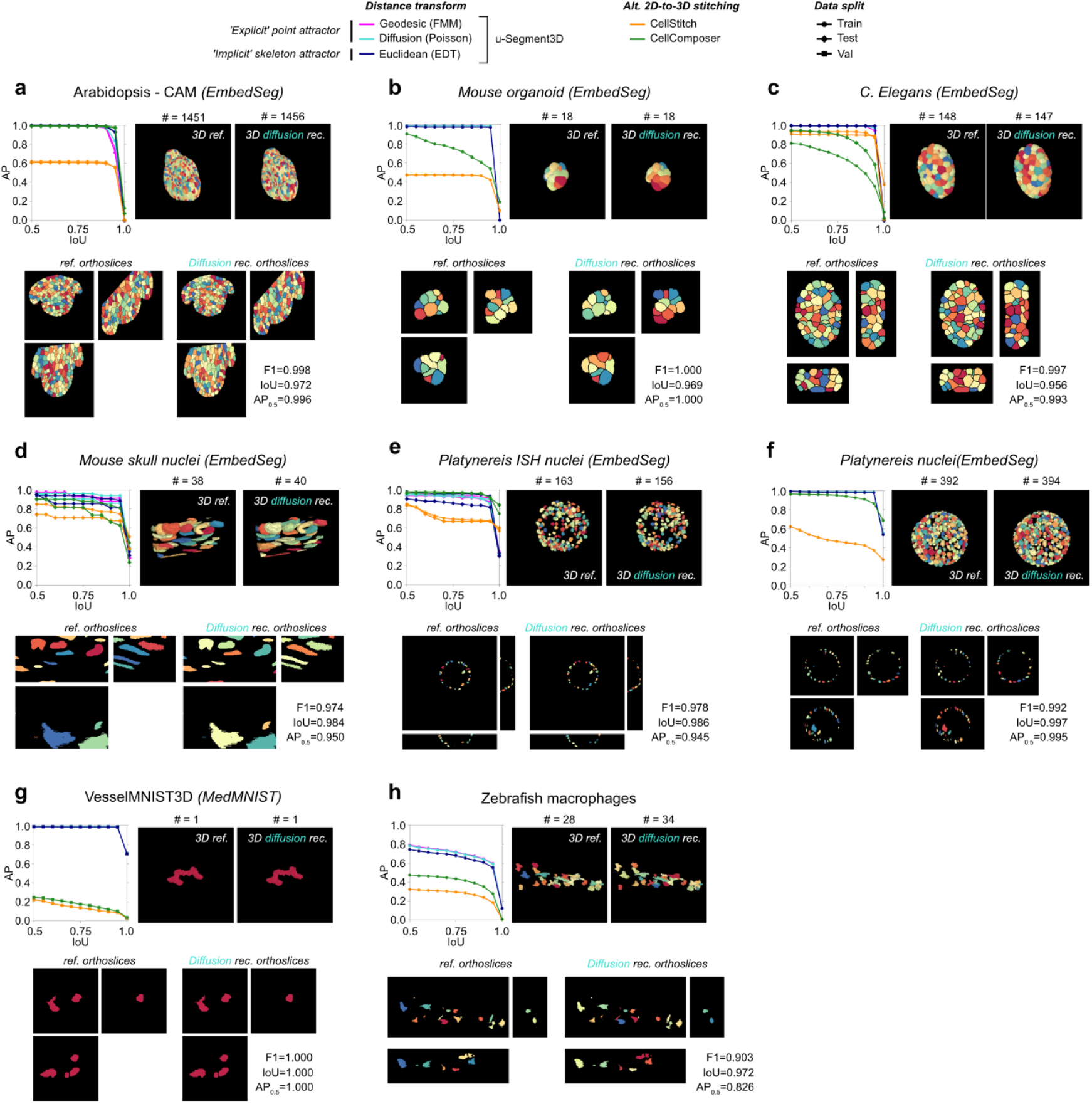
Reconstruction average precision performance of 3D cell shapes from ideal 2D slices sourced from real datasets. Reconstruction performance measured by the mean average precision curve (Methods) using three different 2D distance transforms with u-Segment3D for all datasets not included in Fig. 2d-f in comparison to using CellComposer (green lines) or CellStitch (orange lines) which directly try to stitch 2D segmentations. **a**) Arabidopsis-CAM, **b)** mouse organoid, **c)** *C*.*Elegans*, **d)** mouse skull nuclei, **e)** *Platynereis* ISH nuclei, **f)** *Platynereis* nuclei, **g)** vesselMNIST3D and **h)** zebrafish macrophages. For each dataset, top row, left-to-right: average precision vs intersection over union (IoU) curve; 3D rendering of reference cells, and cells reconstructed using the point-based centroid diffusion distance transform. Bottom row, left-to-right, the respective midplane orthoslices in the three orthogonal views. All available data splits were used for each dataset except VesselMNIST3D for which we used only the validation split. See Suppl. Table 1 for the number of objects and images in each split.

**Extended Data Figure 9.**
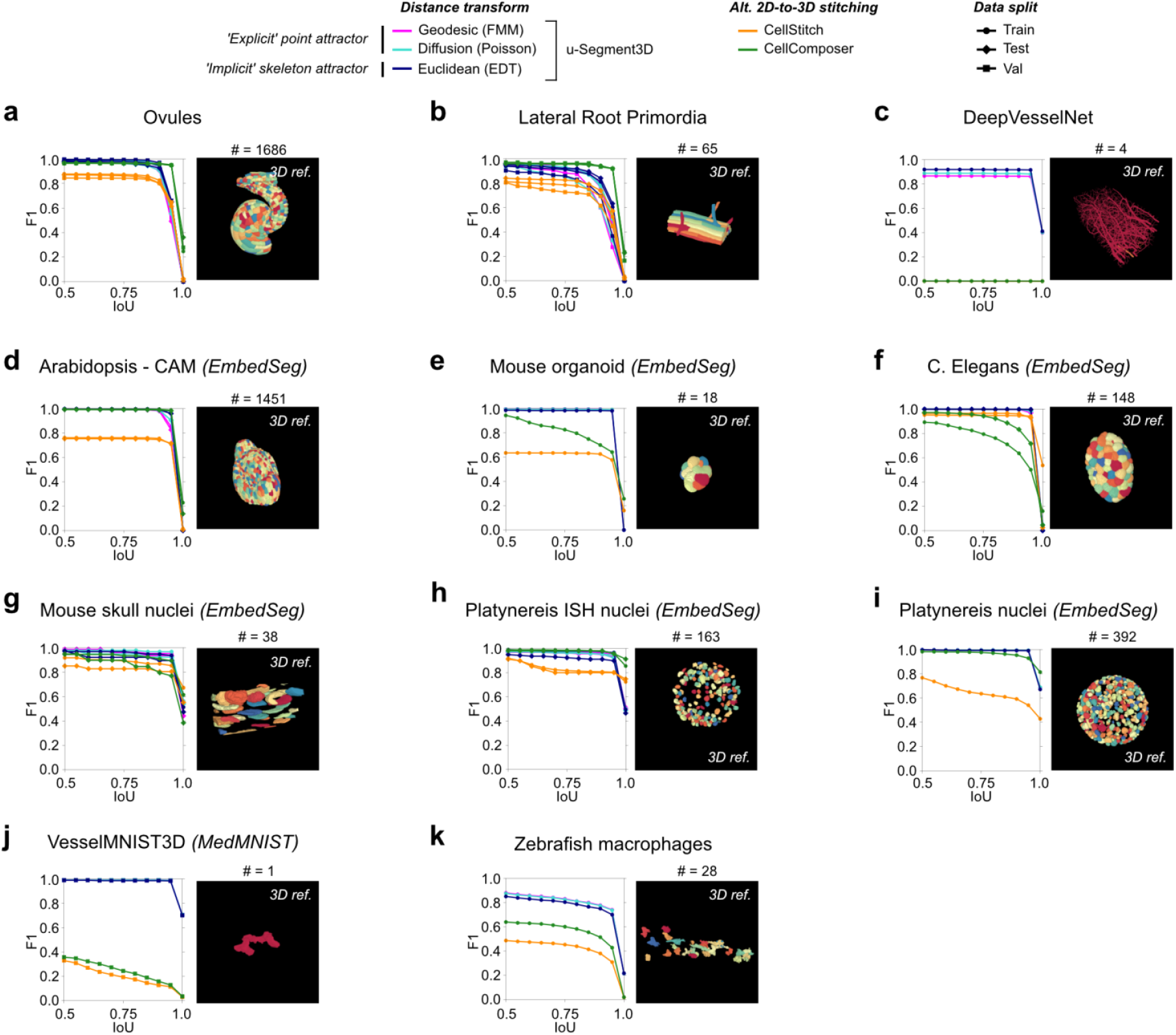
Reconstruction F1 performance of 3D cell shapes from ideal 2D slices sourced from real datasets. Performance of the same shape reconstruction and same methods in Fig. 2 and Extended Data Fig. 8 measured alternatively by F1 score, the harmonic mean of precision and recall (Methods) for the same IoU thresholds as for the average precision curve.

**Extended Data Figure 10.**
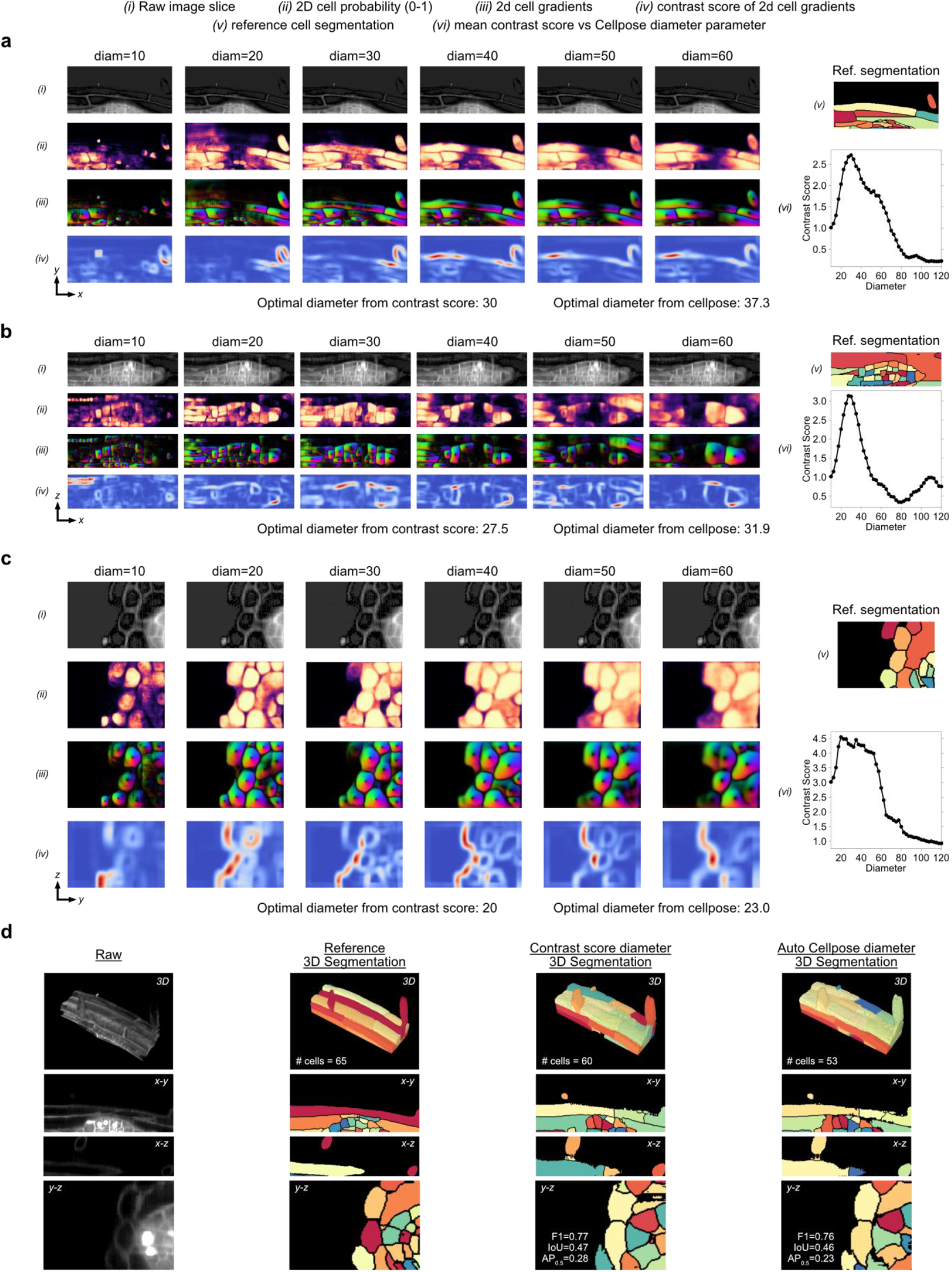
The diameter parameter in pretrained Cellpose models must be individually set in orthogonal views. **a)** Cellpose ‘cyto2’ model outputs and per-pixel u-Segment3D contrast score on the most in-focus (Methods) 2D x-y slice of a Lateral Primordia image as diameter (diam) is increased. (i) Raw input image slice, the same for all values of diameter. (ii) Normalized (0-1) Cellpose 2D pixel probability map colored black=0 to yellow=1. (iii) Unit-normalized Cellpose 2D predicted gradients colored by direction. (iv) Contrast score of predicted Cellpose 2D gradient (Methods, Extended Data Figure 10a). (v) Reference cell segmentation for the raw 2D input. (vi) Mean contrast score averaged over the image for each value of diam. The diameter with maximum contrast score is taken as the optimal diameter by u-Segment3D. **b)** Same as in a) for the most in-focus 2D x-z slice and **c)** the most in-focus 2D y-z slice. **d)** From left-to-right, 3D renderings of the raw 3D volume, reference 3D segmentation, u-Segment3D segmentation using the direct method and diameter in each orthoview set by contrast score or by the Cellpose ‘cyto2’ model (top), and corresponding mid-plane orthoslices in all three views (below).

**Extended Data Figure 11.**
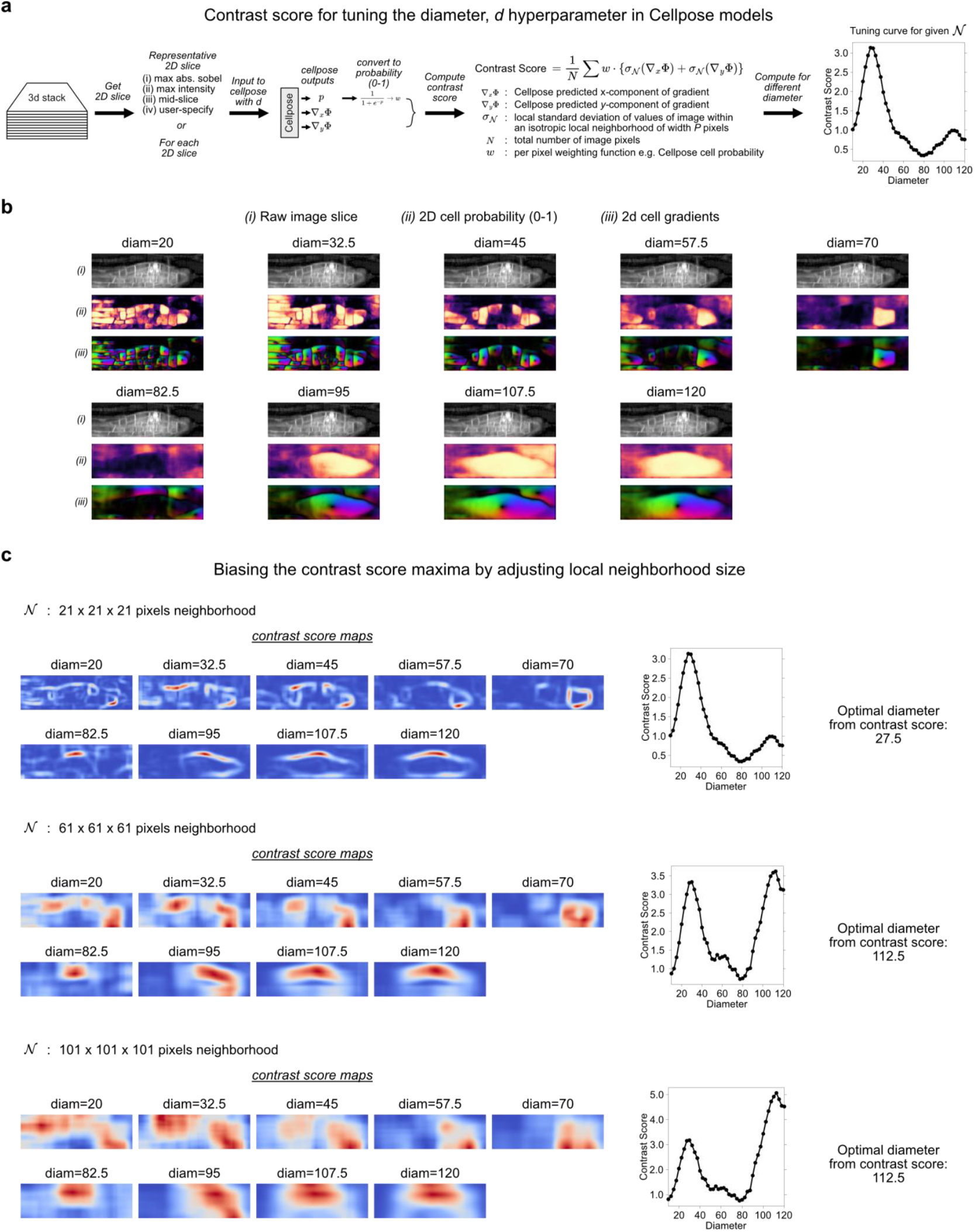
Semi-automatic determination of the diameter parameter in pretrained Cellpose models using local variance. **a)** Computational workflow and definition of a contrast score function using a local neighborhood of width *P* pixels to evaluate 2D Cellpose outputs when the diameter parameter is set to *d*. **b)** (i) Raw input image slice, (ii) normalized (0-1) Cellpose 2D pixel probability map colored black=0 to yellow=1, and (iii) unit-normalized Cellpose 2D predicted gradients colored by direction for 9 equisampled diameters in the range *d* = [20, 120]. **c)** Contrast score maps (colored blue-to-red for low-to-high values) for the same *d* as in b) (left) and the resulting contrast score function and optimal diameter inferred (right), using a neighborhood of width *P* = 21, 61, 101 pixels (top-to-bottom).

**Extended Data Figure 11.**
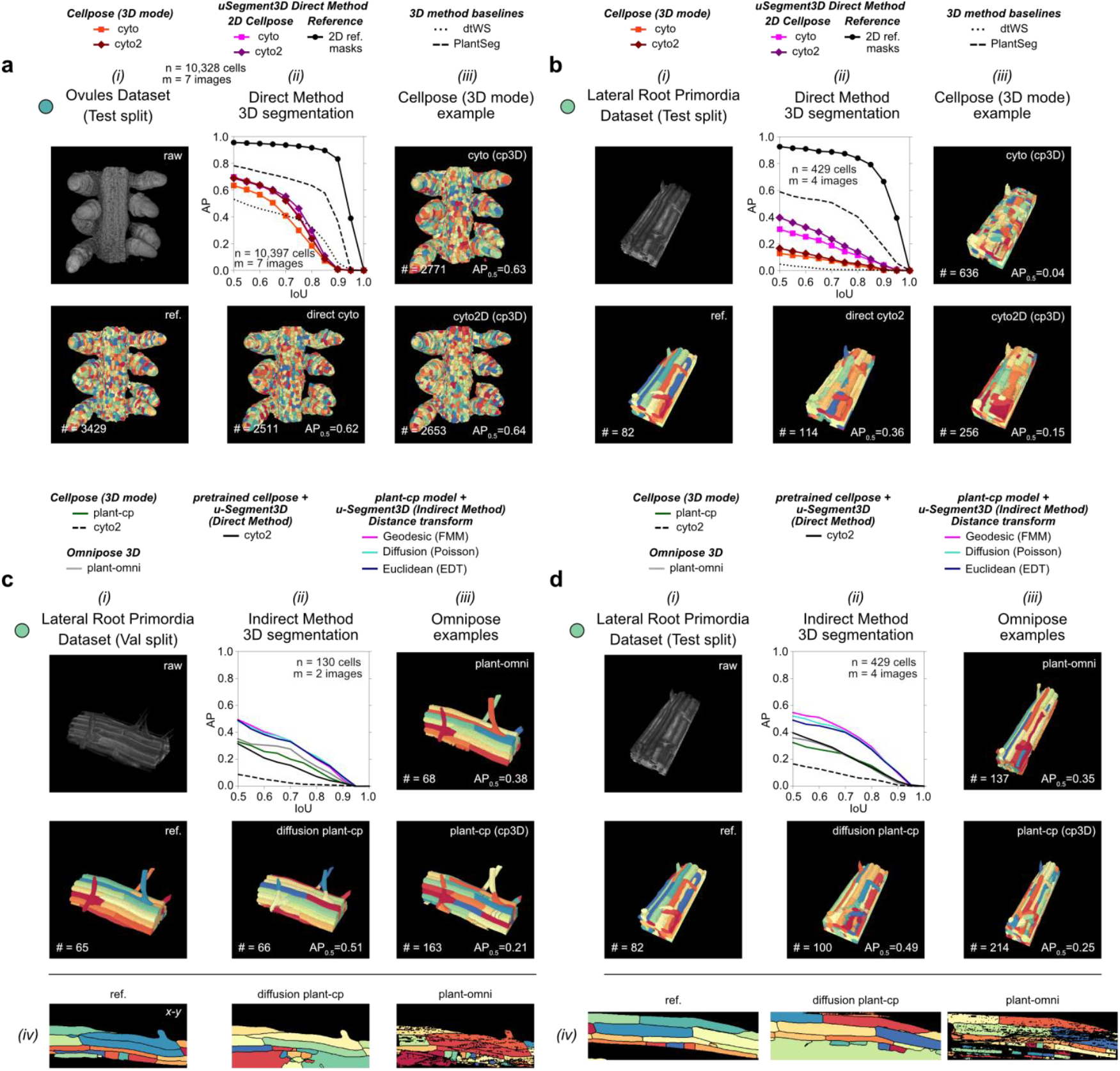
Performance of u-Segment3D using pretrained vs specialized plant Cellpose segmentation models. **a)** Performance of consensus u-Segment3D and Cellpose 3D segmentations using the same pretrained Cellpose 2D models on the test split of Ovules relative to two baselines: classical unsupervised distance transform watershed and PlantSeg, a specialized 3D UNet model trained directly with the 3D masks. (i) example volume and corresponding reference segmentation; (ii) AP curves of all models, with best 3D reconstruction from ideal 2D slices (black line) (top) and segmentation of best pretrained model with u-Segment3D (bottom); (iii) Cellpose 3D segmentations using cyto (top) or cyto2 (bottom) models. **b)** Performance of consensus u-Segment3D and Cellpose 3D segmentations using the same pretrained Cellpose 2D models on the test split of the Lateral Root Primordia (LRP) dataset relative to two baselines: classical unsupervised distance transform watershed and PlantSeg, a specialized 3D UNet model trained directly with the 3D masks. (i)-(iii) similar to a). **c)** Performance on LRP val split using u- Segment3D or Cellpose 3D and plant-cp: a specialized Cellpose 2D model trained on LRP, with plant-omni: a specialized Omnipose 3D model trained on LRP natively in 3D. (i)-(ii) similar to a). (iii) Native 3D segmentation using plant-omni (top) or Cellpose 3D segmentation with plant-cp (bottom). (iv) mid x-y slice of the reference, u-Segment3D diffusion centroid transform plant-cp consensus and 3D plant-omni segmentation (left-to-right). **d)** Same as c) for the test split of LRP.

**Extended Data Figure 13.**
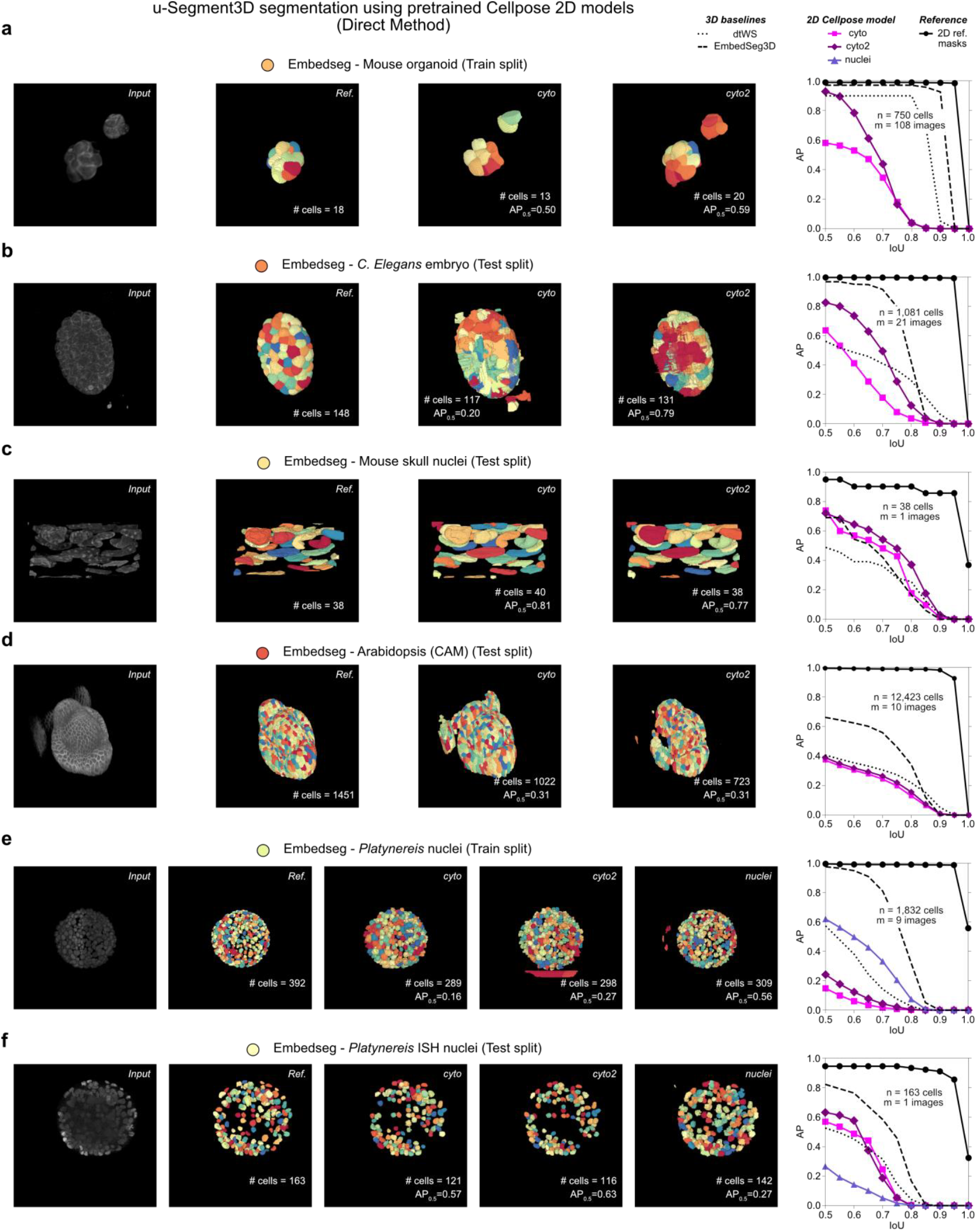
Performance of 2D-to-3D segmentation for real datasets using u- Segment3D with pretrained cellpose2D model outputs. **a)** 3D cell segmentation performance of the mouse organoid (from Embedseg^15^) for the train data split, n=740 cells, m=108 volumes) using pretrained Cellpose 2D models with u-Segment3D and the direct method illustrated in Fig.3a relative to baseline native 3D segmentation: classical unsupervised distance transform watershed (dtWS) (dotted black line) and training an EmbedSeg3D model (dashed black line) (Fig. 4, Methods). Left-to-right: 3D rendering of the raw volume, reference 3D segmentation, generated 3D segmentations for each Cellpose 2D model and the average precision (AP) curve of each model coplotted with the AP curve of the best 3D reconstruction with ideal 2D slices from the three orthoviews (black line with circles). The same as a) for **b)** C. Elegans embryo (test data split, n =1,081 cells, m = 21 images), **c)** mouse skull nuclei (test data split, n=38 cells, m=1 image), **d)** Arabidopsis (CAM) (test data split, n=12,424 cells, m=10 images), **e)** Platynereis nuclei (train data split, n=1,832 cells, m=9 images), **f)** Platynereis ISH nuclei (test data split, n=163 cells, m=1 image). For the Platynereis nuclei which are approximately spherical, we additionally evaluated the performance of the Cellpose ‘nuclei’ 2D model (light purple line with triangles).

**Extended Data Figure 14.**
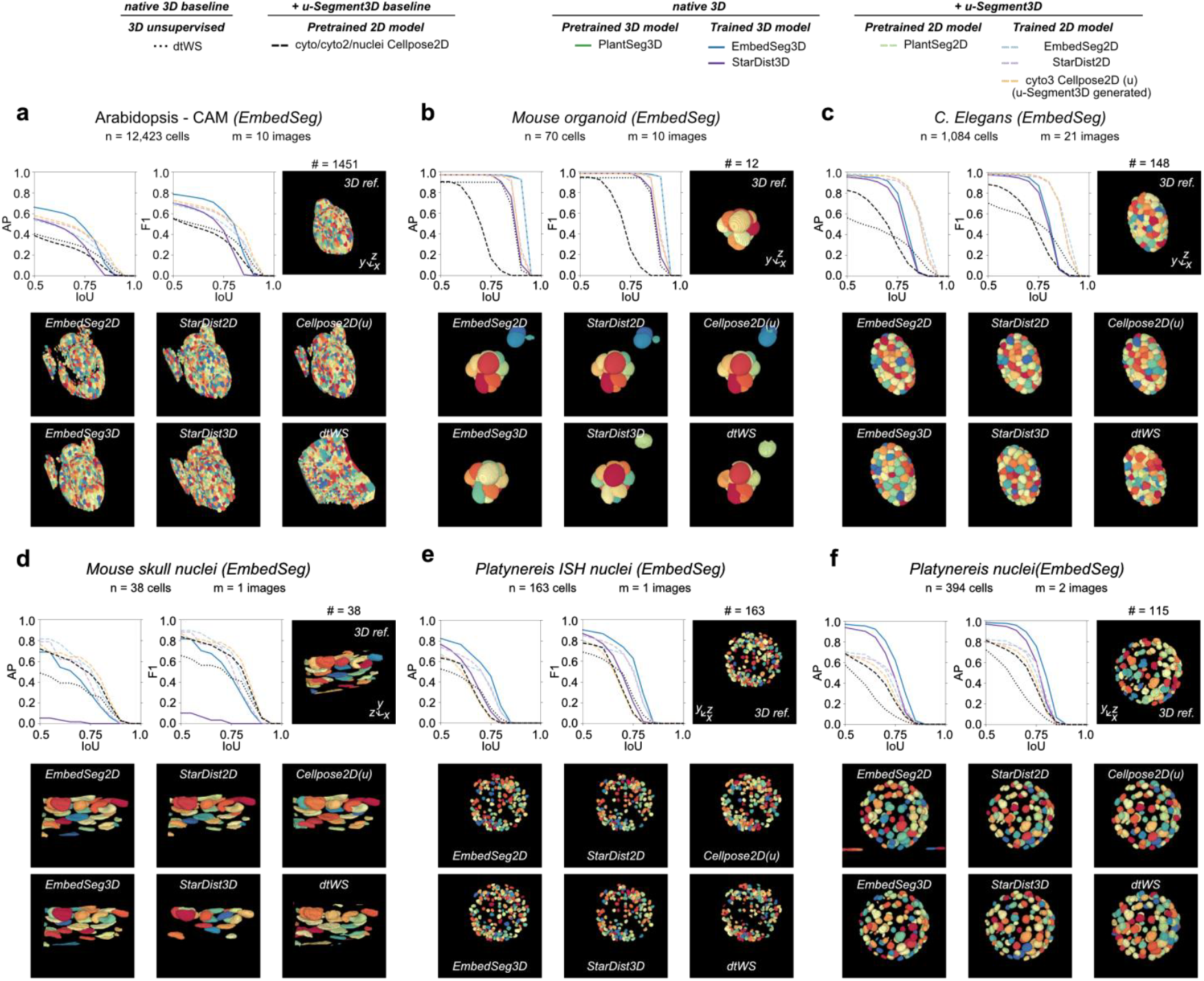
Performance of specialist native 3D trained segmentation models and consensus u-Segment3D segmentation with specialist trained 2D segmentation. **a)** 3D cell segmentation performance of the Arabidopsis (CAM) dataset from Embedseg^15^ based on the test data split, n=12,423 cells, m=10 volumes) for native 3D trained (solid line, darker hue) and u-Segment3D consensus segmentation (dashed line, lighter hue) relative to the baseline of classical unsupervised distance transform watershed (dtWS) (dotted black line, Methods). Top row left-to-right: AP and F1 curve evaluation, and 3D rendering of the reference 3D segmentation. Bottom panels: 3D segmentations from each model with name of 2D model denoting the result from consensus u-Segment3D segmentation. The same as a) for test splits (Methods) of **b)** Mouse organoid (n=70 cells, m=10 images), **c)** C. Elegans embryo (n =1,081 cells, m = 21 images), **d)** mouse skull nuclei (n=38 cells, m=1 image), **e)** Platynereis ISH nuclei (n=163 cells, m=1 image), **f)** Platynereis nuclei (n=394 cells, m=2 images). 2D Cellpose segmentations were generated from network outputs using u-Segment3D’s method (methods), as denoted by the (u) annotation.

**Extended Data Figure 15.**
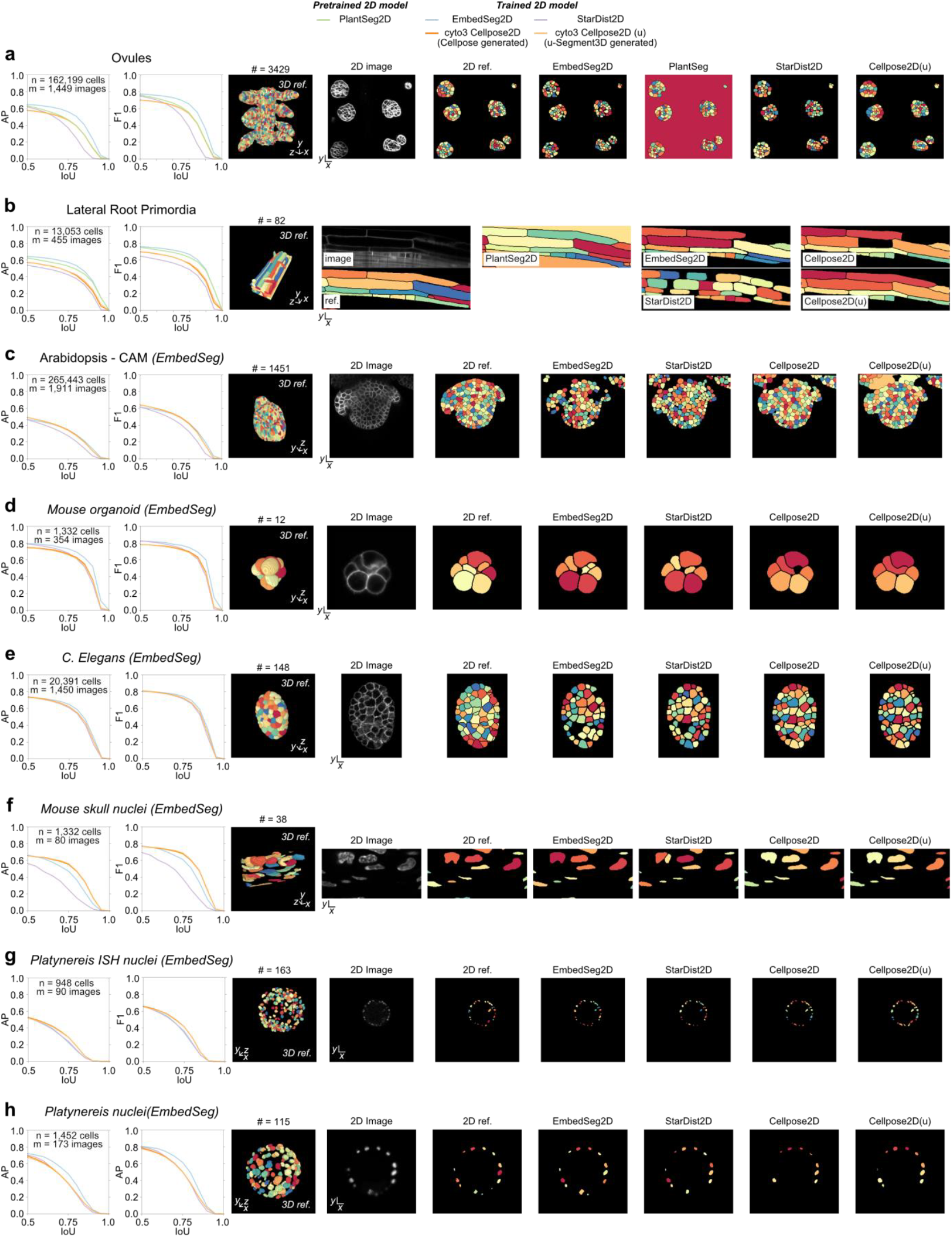
Performance of training specialist 2D segmentation models on 2D x-y, x-z and y-z slices of 3D volumes. **a)** 2D cell segmentation performance of each trained 2D model on sampled 2D x-y, x-z and y-z slice images from the test 3D volumes of the Ovules dataset (n=162,199 cells, m=1,449 images). Left-to-right: AP and F1 curve, 3D rendering of the reference 3D segmentation, raw image of the mid x-y slice and its reference 2D segmentation, and predicted 2D segmentation of the mid x-y slice with trained models. The same as a) of 2D images sampled from the test volumes (Methods) of **b)** Lateral Root Primordia (n=13,053 cells, m=455 images), **c)** Arabidopsis - CAM (n=265,443 cells, m=1,911 images), **d)** Mouse organoid (n=1,332 cells, m=354 images), **e)** C. Elegans embryo (n =20,391 cells, m = 1,450 images), **f)** Mouse skull nuclei (n=1,332 cells, m=80 images), **g)** Platynereis ISH nuclei (n=948 cells, m=90 images), **h)** Platynereis nuclei (n=1,452 cells, m=173 images). Cellpose 2D segmentations were generated from Cellpose outputs using either Cellpose or using u-Segment3D, denoted by the absence or presence of an additional (u) annotation respectively.

**Extended Data Figure 16.**
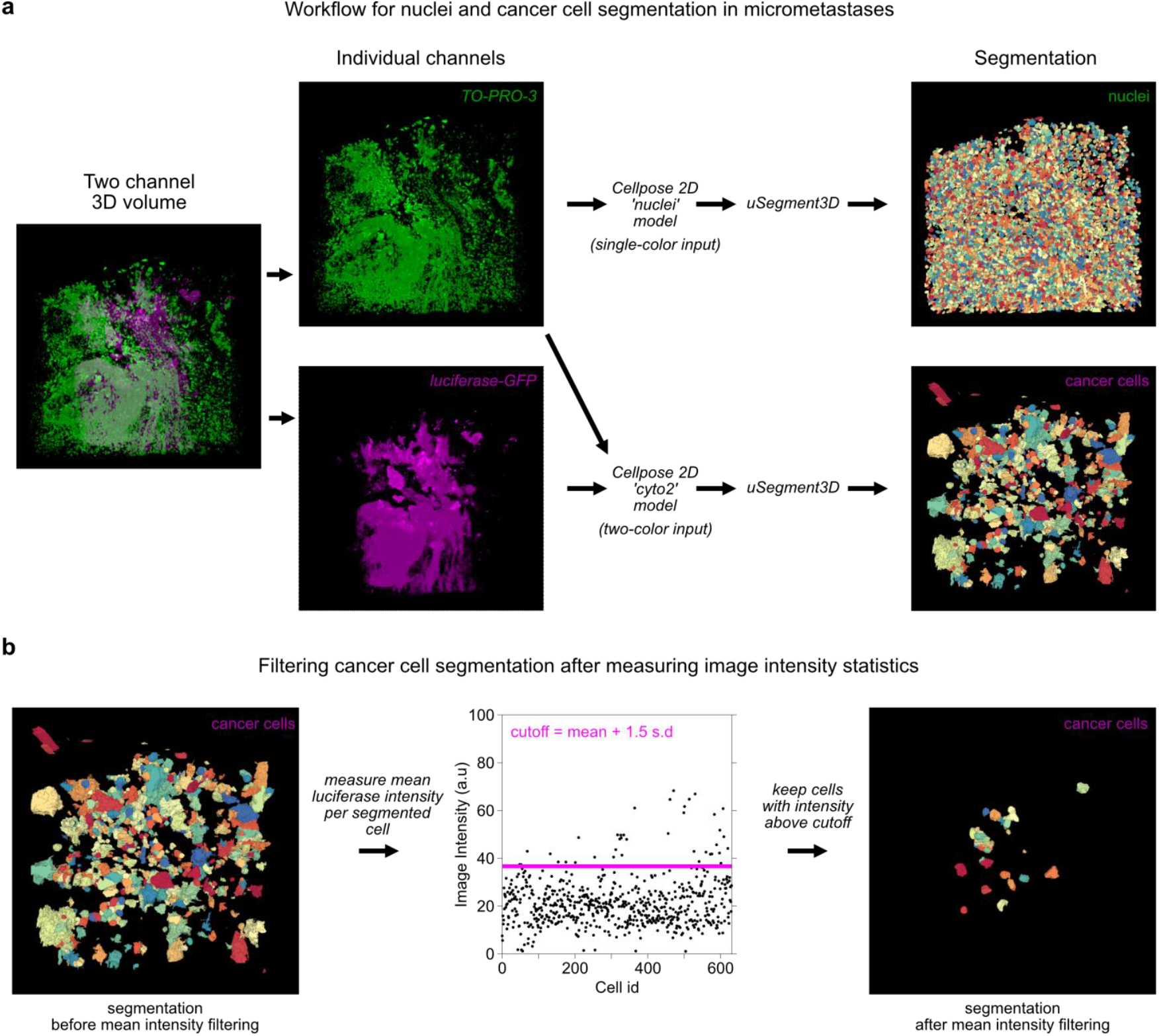
u-Segment3D generated consensus 3D segmentations can be filtered with image statistics to improve specificity for weakly labeled cells. **a)** Workflow and 3D rendering of the resulting segmentation using the nuclei-only stained channel (TO-PRO-3, green) to extract lung nuclei (top branch); using both nuclei (green) and luciferase-GFP (magenta) channel to segment YUMM 1.7 melanoma cells in micrometastatic colonies (bottom branch). **b)** Procedure to leverage the luciferase-GFP intensity to filter cells with a mean luciferase-GFP intensity above a cutoff (magenta) of mean + 1.5 standard deviation across all cells.

## Supplementary Tables

**Supplementary Table 1.** Table of all parameters in u-Segment3D code with explanation and tuning guidance. The variable names correspond to that used in the GitHub source code.

Plants
| Name | Description | Pixel Size (Z,Y,X) [$\mu\text{m}^3$] | Used Microscope | Isotropic Rescaling factor | Total #images (Train/Test/Val) | Total #cells |
| --- | --- | --- | --- | --- | --- | --- |
| Lateral Primordia<sup>7</sup> | *Arabidopsis thaliana* lateral root of the line sC111 were used at 5 day post germination | (0.25, 0.1625, 0.1625) | Multi-view Selective Plane Illumination Microscopy | 0.5 | 21 / 4 / 2 | 2,068 / 429 / 130 |
| Ovules<sup>7,125</sup> | *Arabidopsis thaliana* ovules stained with SR2200 and TO-PRO-3 iodide | (0.235, 0.075, 0.075) * | Confocal laser scanning microscopy | 1 | 22 / 7 / 2 | 23,860 / 10,328 / 2,839 |

\* Resampling according to the published pixel size did not produce the expected isotropic appearance. We find (0.235, 0.15, 0.15) to better generate a more isotropic appearance.
Lateral primordia was downloaded from https://osf.io/2rszy/
Ovules was downloaded from https://osf.io/w38uf/

**Supplementary Table 2.** Summary of datasets used for validation of u-Segment3D. Unless otherwise noted, all images were first resampled to isotropic voxel resolution by appropriate downsampling of x-y slices and then resized isotropically by the indicated rescaling factor.

Embedseg
| Name | Description | Pixel Size (Z,Y,X) [$\mu\text{m}^3$] | Used Microscope | Isotropic Rescaling factor | # images (Train/Test/Val) | Total # cells |
| --- | --- | --- | --- | --- | --- | --- |
| Arabidopsis-Cells-CAM-small<sup>126,127</sup> | *Arabidopsis Thaliana* YFP membrane labelled and imaged between 24 and 28 days after germination | (0.26, 0.22, 0.22) | Confocal Microscopy | 1 | 11 / 10 / - | 12,016 / 12,423 / - |
| C.elegans-Cells-HK<sup>54</sup> | C.elegans embryos membrane labeled. | (0.25, 0.25, 0.25) | Confocal Microscopy | 1 | 54 / 21 / - | 3,479 / 1,081 / - |
| Mouse-Organoid-Cells-CBG<sup>15</sup> | Mouse Embryonic Stem Cells, R1 cell line, labeled membrane | (1.0, 0.1733, 0.1733) | Selective Plane Illumination Microscopy | 0.5 | 108 / - / - | 750 / - / - |
| Mouse-Skull-Nuclei-CBG<sup>15</sup> | Nuclei of the skull region of developing mouse embryos, labeled with DAPI | (0.200, 0.073, 0.073) * | Inverted Zeiss LSM 880 Microscope | 0.5 | 2 / 1 / - | 150 / 38 / - |
| Platynereis-Nuclei-CBG<sup>15</sup> | Nuclei of whole-mount *Platynereis dumerilli* specimens | (2.031, 0.406, 0.406) | Simultaneous Multi-view | 1 | 9 / - / - | 1832 / - / - |
|  | at stages between 0 to 16 hours post fertilization, injected with a fluorescent nuclear tracer |  | Light-Sheet Microscopy |  |  |  |
| Platynereis-ISH-Nuclei-CBG<sup>15</sup> | Nuclei of whole-mount *Platynereis dumerilli* specimens at stage of 16 hours post fertilization, labeled with DAPI | (0.45, 0.45, 0.45) | Laser Scanning Confocal Microscopy | 1 | 2 / 1 / - | 486 / 159 / - |

\* For Mouse-Skull-Nuclei-CBG we found that pretrained Cellpose 2D models and u-Segment3D performed better without resizing to isotropic voxels and applying isotropic rescaling factor.
All of these 3D datasets were downloaded from https://github.com/juglab/EmbedSeg/releases/tag/v0.1.0

**Supplementary Table 3.** u-Segment3D settings and parameters for testing the reconstruction of 3D segmentation from ideal 2D segmentations with public datasets.

VesselMNIST3D
| Name | Description | Pixel Size (Z,Y,X) | Used Microscope | Isotropic Rescaling factor | # images (Train/Test/Val) | Total # cells |
| --- | --- | --- | --- | --- | --- | --- |
| VesselMNIST3D<sup>128,129</sup> | Derived from the Intra dataset. Vessel segments were generated from 103 3D meshes of entire brain vessels. | (1,1,1) | Time-of-Flight Magnetic Resonance Angiography (TOF-MRA) | 9.14 | 1,335/382/192 (*191) | 1,335 / 382 / 191<sup>^</sup> |

<sup>^</sup>We only used the val split as all images were single-component and morphological properties similar across data splits.
\* 1 of the images in the val split was found to be blank.
The dataset was downloaded as .npz from https://zenodo.org/records/10519652.

**Supplementary Table 4.** u-Segment3D settings and parameters for consensus 3D segmentation from the output of pretrained Cellpose 2D predictions with public datasets. The model inversion parameter is for Cellpose2 code. Model inversion is unnecessary for Cellpose3 code.

DeepVesselNet
| Name | Description | Pixel Size (Z,Y,X) | Used Microscope | Isotropic Rescaling factor | # images (Train/Test/Val) | Total # cells |
| --- | --- | --- | --- | --- | --- | --- |
| DeepVesselNet<sup>10,130,131</sup> | Synthetic generated dataset with images of 325 x 304 x 600. Vessel intensities were randomly chosen in the interval [128, 255] and non-vessel intensities from the interval [0 – 100]. | (1,1,1) | - | 1 | 136 (135*) / - / - | 410* / - / - |
|  | Gaussian noise was then randomly applied. |  |  |  |  |  |

\* We could not download and unzip 1 of the raw images in the archive.
The dataset was downloaded from https://github.com/gieseckow/deepvesselnet/wiki/Datasets

**Supplementary Table 5.**
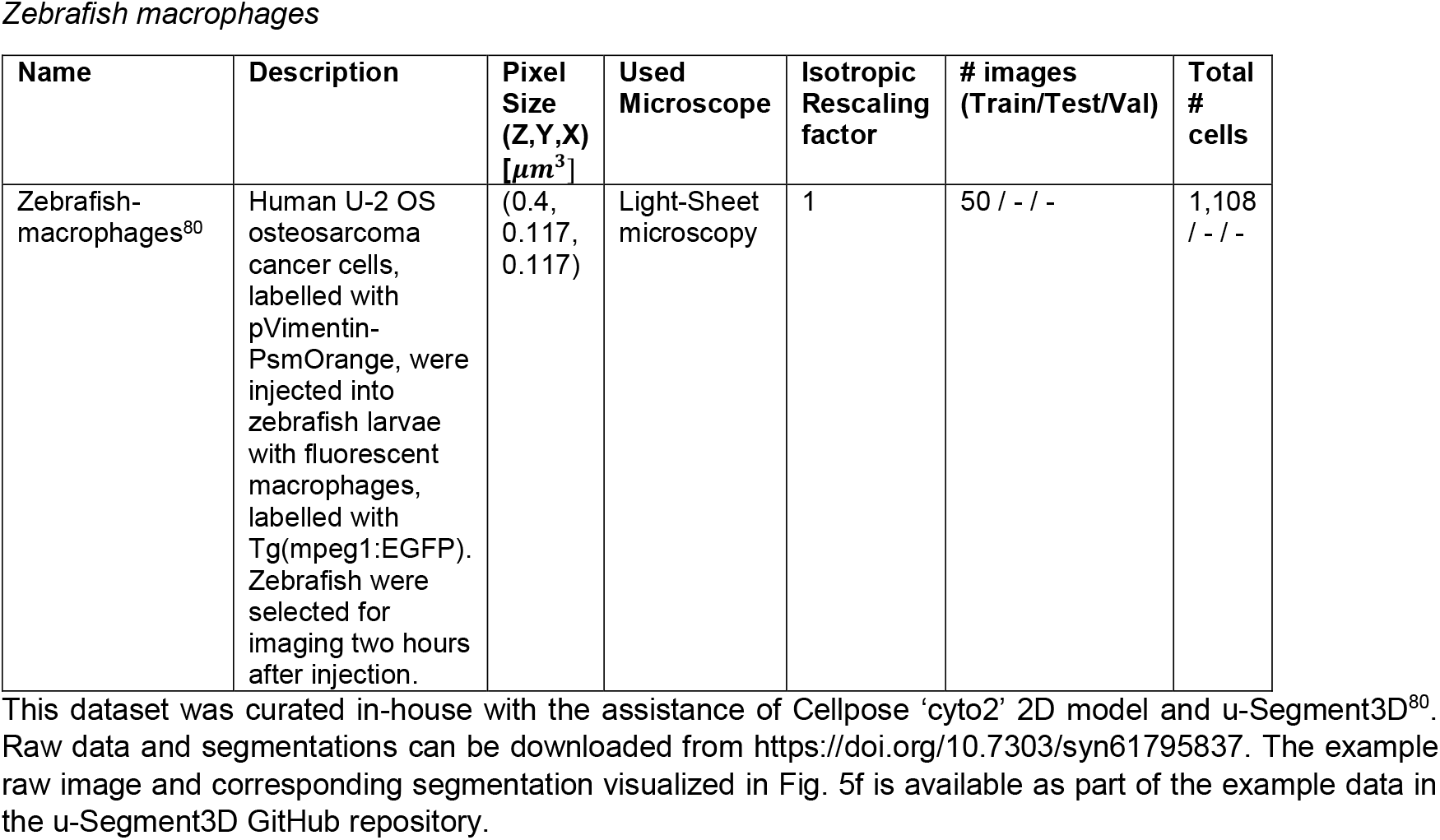
Cellpose2D and u-Segment3D settings and parameters for consensus 3D segmentation on additional demonstration datasets. The model inversion parameter is for Cellpose2 code. Model inversion is unnecessary for Cellpose3 code.

## Supplementary Movies

**Supplementary Movie 1. u-Segment3D is a toolbox to integrate 2D segmentations from orthoviews into a single consensus 3D segmentation**. Illustration of applying a 2D segmentation method to generate instance segmentations of cells within an image volume slice-by-slice from x-y, x-z, and y-z directions. This produces 3 separate 2D stacks which can be integrated using u-Segment3D into a single 3D segmentation. As visualized by exploding the end segmentation, this strategy can successfully extract every cell in dense cleared tissue volumes, without data training.

**Supplementary Movie 2. Gradient descent dynamics of foreground cell coordinates using different 2D transforms**. For each transform, we visualize i) the computed gradients color-coded by direction; ii) the advection of all foreground points (white dots) along the gradients using gradient descent; iii) the gradient descent trajectory (magenta) of equisampled boundary points; and iv) final reconstructed cells. Reconstructed cells are colored using the magma colorscheme from light-yellow to black. The color range is the same for all transforms and is set to be from zero (light-yellow) to 1.5 times the maximum cell id of reference cells (black). Perfectly reconstructed cells such as that using explicit ‘point’ source transforms, should be uniquely colored from light-yellow to purple. In contrast, oversegmentation causes an overabundance of blacks and undersegmentation an overabundance of yellows.

**Supplementary Movie 3. Comparison of the spatial proximity clustering used by Cellpose 3D and u- Segment3D’s image-based connected component analysis on noisy cell tracking challenge datasets with non-optimized model parameters**. For each example, the movie first rotates the rendering of the output 3D segmentation then flies through individual x-y slices where individual cells are delineated by black outlines. Each cell was assigned one of 16 colors from the ‘spectral’ colorscheme. The same cell in 3D render and 2D slices have the same color.

**Supplementary Movie 4. Gradient descent dynamics of foreground cell coordinates under 3D gradients reconstructed from 2D gradients**. Visualization of the propagated foreground points (magenta dots) under suppressed gradient descent with momentum for 200 iterations relative to initial positions (black dots) for the shapes and parameters in Extended Data Fig. 7b.

**Supplementary Movie 5. Gradient descent dynamics during 3D reconstruction of ovules, lateral root primordia and vasculature from ideal 2D segmented stacks**. Visualization of the propagated foreground points (magenta dots) under suppressed gradient descent with momentum for 250 iterations relative to initial positions for the exemplar images from ovules, lateral root primordia, and DeepVesselNet in Fig. 2d-f. Initial foreground positions are colored according to the unique reference cell they belong to. Rotation of the 3D rendering of reference (left) and reconstructed cells using centroid diffusion (middle) and Euclidean distance transforms (right). Whereas centroid point diffusion advects foreground points towards single point attractors, EDT advects foreground points towards the skeleton. When gradient decay, τ is sufficiently high as in DeepVesselNet, the behavior of both distance transforms are similar and points are minimally advected to separate individual vessel networks without fragmentation.

**Supplementary Movie 6. Gradient descent dynamics of 3D reconstruction of lateral root primordial from only y-z, x-y and y-z and x-y, x-z and y-z Embedseg2D 2D segmented stacks**. Visualization of the propagated foreground points (magenta dots) under suppressed gradient descent with momentum for 100 iterations relative to initial positions (black dots) for the exemplar test volume of lateral root primordia in Fig. 4f from using only y-z, both x-y and y-z and all three x-y, x-z and y-z Embedseg2D 2D segmented stacks (left-to-right). Rotation of the 3D rendering of the corresponding output u-Segment3D consensus 3D segmentations.

**Supplementary Movie 7. Segmentation of a movie of thin MDA231 human breast carcinoma cells embedded in collagen from the 3D cell tracking challenge using u-Segment3D with stacks of 2D x-y segmentations only**. Flythrough of x-y slices and 3D rotation of timepoint 0 of raw images and u-Segment3D segmentations. The x-y segmentations were generated using u-Segment3D’s automatically determined optimal Cellpose 2D model parameters. Timelapse of raw image and u-Segment3D segmentations for all 15 timepoints.

**Supplementary Movie 8. Segmentation of unwrapped surface cells of a Drosophila embryo over time using u-Segment3D to aggregate only 2D x-y segmentations only**. Rendering of the surface extracted from binary embryo segmentation of each timepoint using u-Segment3D. Timelapse of the topographic volumes constructed by mapping a surface proximal volume into cartographic coordinates using u-Unwrap3D. Rotation and flythrough of the raw topographic volume at timepoint 25 and corresponding u-Segment3D segmentation. Timelapse of the topographic z-slice that best captures all surface cells and timelapse of the mid-slice y-z cross section (same as Fig. 5m), with corresponding u-Segment3D segmentation.

**Supplementary Movie 9. u-Segment3D postprocessing enables recovery of missing surface protrusions in the 3D segmentation of a HBEC cell aggregate**. Rotating 3D renderings at each part of the workflow depicted in Fig. 6a.; (i) raw image, (ii) deconvolved image, (iii) consensus u-Segment3D segmentation of deconvolved image with Cellpose 2D outputs, (iv) segmentation following label diffusion and (v)segmentation following guided filtering postprocessing. Final volumes were meshed, exploded and rotated to visualize the whole cell segmentations this workflow extracts.

**Supplementary Movie 10. Comparison of the segmentation of vessel sprouting in zebrafish without and with u-Segment3D postprocessing**. Rotating 3D rendering of (i) raw image, (ii) consensus u- Segment3D binary segmentation of raw image using Cellpose 2D outputs and (iii) segmentation after label diffusion and guided filtering postprocessing.

**Supplementary Movie 11. u-Segment3D segmentation of 43**,**779 cells in a** ≈ 35*μ***m x 1.5**mm **x 1.5**mm **CYCIF multiplexed tissue section of metastatic melanoma**. Flythrough of x-y slices of the entire volume and then in a zoomed-in region. Subsequent flythrough of x-z and y-z slices of the entire volume with simultaneous zoomed-in regions. This movie is compressed as MPEG-4 with the HEVC codec and can be viewed, for example, with the VLC media player.

**Supplementary Movie 12. u-Segment3D segmentation enabled detection of weakly fluorescent lung micrometastases in cleared tissues**. Rotations of 3D rendered input image with merged and separated TO-PRO-3 (nuclei, green) and luciferase-GFP (YUMM 1.7 cancer cells, magenta) channels; initial consensus u-Segment3D segmentations using Cellpose 2D outputs for nuclei and cancer cells; and cancer cells after filtering out spurious segmentations by luciferase-GFP intensity.

**Supplementary Movie 13. u-Segment3D segmentation of heterogeneous cell structures in brain tissue labelled using coCATs**. Rotating rendering of input image stack with consensus 3D segmentation of the larger pockets of extracellular space. Slice-by-slice flythrough of the segmentation, colored and overlaid using transparency onto raw x-y, x-z, and y-z slices. Individual pockets are additionally delineated by green outlines.

